# Physical modeling of embryonic transcriptomes identifies collective modes of gene expression

**DOI:** 10.1101/2024.07.26.605398

**Authors:** Dominic J. Skinner, Patrick Lemaire, Madhav Mani

## Abstract

Starting from one totipotent cell, complex multicellular organisms form through a series of differentiation and morphogenetic events, culminating in a multitude of cell types arranged in a functional and intricate spatial pattern. To do so, cells coordinate with each other, resulting in dynamics which follow a precise developmental trajectory, constraining the space of possible embryo-to-embryo variation. Using recent single-cell sequencing data of early ascidian embryos, we leverage natural variation together with modeling and inference techniques from statistical physics to investigate development at the level of a complete interconnected embryo – an embryonic transcriptome. After developing a robust and biophysically motivated approach to identifying distinct transcriptomic states or cell types, a statistical analysis reveals correlations within embryos and across cell types demonstrating the presence of collective variation. From these intra-embryo correlations, we infer minimal networks of cell-cell interactions, which reveal the collective modes of gene expression. Our work demonstrates how the existence and nature of spatial interactions along with the collective modes of expression that they give rise to can be inferred from single-cell gene expression measurements, opening up a wider range of biological questions that can be addressed using sequencing-based modalities.

## I. INTRODUCTION

Development reliably produces complex and highly structured organisms in the presence of external perturbations and intrinsic stochasticity [1]. Strikingly, the dynamics which give rise to these complex structures are also constrained, or canalised in the language of Waddington [2]. Yet phenotypic variation across individuals must be present to facilitate evolution [3]; how the nature of phenotypic variation is shaped by the nature of the dynamical constraints is largely unknown. Single cell gene expression measurements, though limited in quality and an incomplete measurement of a cell’s state, provide a high-dimensional cellular phenotype [4] that has led to an avalanche of cell atlases focused on phenotypic identification of cell types across a multitude of species [5–7]. Phenotypic variation, however, exists not solely at the scale of cells but also at the scale of embryos, producing variation in organismal function. Cells within an embryo interact with one another through intricate mechano-chemical mechanisms, which correlate the states of cellular gene expression. Much like how the couplings between infinitesimal material portions of a violin string sustain large-scale collective oscillations, we expect the complex coupling of cells in embryos to give rise to collective modes of gene expression. Far from producing a pure note, we anticipate that the intricate inter-cellular couplings creates a rich timbre of complex collective modes of gene expression in embryos. Such modes are features, or phenotypes, of the embryo, existing at a scale above the individual cells that comprise it. To our knowledge, the present study is the first attempt to identify and characterize these collective modes of embryonic gene expression using a combination of simple physical modeling and careful statistical analyses of embryonic transcriptomic data. As such, we aim to bring into focus the concept of an embryonic transcriptome and the possibility of doing transcriptomic physics. Practically speaking, we wish to demonstrate that single-cell sequencing data can be leveraged to study a far richer canvas of biological questions than the identification of cell types. In addition we wish to propose that, much like in condensed matter systems, the lower-dimensional phenotypic space spanned by the collective modes of gene expression in an embryo are functionally relevant, encoding a space in which developmental dynamics ensue and where variation acts.

Statistical physics provides frameworks to quantify and investigate the collective self-organization and fluctuations of systems comprised of many individual components, strongly interacting with each other. Such approaches are insightful for the study of complex systems, from spin glasses [8] and bird flocks [9], to gene regulatory networks [10] and biological neural networks [11]. Here, we apply these frameworks to study collective behaviors in the developing ascidian embryo observed using single-cell RNA sequencing data of individuals at distinct stages of development. We follow the tightly controlled program from the initiation of zygotic transcription, where increasing numbers of genes are transcriptionally activated in precise spatial locations as development ensues, finding that variation in gene expression is coupled across cells within the same embryo, even after accounting for differences in staging time and the finiteness of our datasets. Moreover, we find spatial structure in the cell-cell correlations for zygotic transcripts, in contrast to the variation in maternal transcripts, which present no spatial structure in their variation. To ascertain the nature of cell-cell couplings, whether through inheritance of transcripts via cell divisions or inter-cell signalling, we use statistical physics modeling to infer minimal, or sparse, networks of cell-cell interactions that give rise to the observed complex, or dense, set of cell-cell correlations. The inferred network of interactions within the context of a simple statistical physical model allows us to quantify and visualize the collective modes of gene expression, which are not directly accessible from the single-cell sequencing data. This work represents an attempt to combine data-driven and physical modeling to reveal new perspectives on the developing organism and to pave the way for doing “transcriptomic physics”.

## II. BACKGROUND

### Biological Context

In order to ground our goals and approaches in a concrete setting we focus on the early development of the ascidian, or sea squirt, *Phallusia mammillata*. This organism displays several attractive features, which we now discuss, that make it an ideal setting to develop our approach. The ascidian cell lineage is invariant, with the same series of synchronous cell divisions occurring in each embryo [12–14], Fig. 1A, which also provides a consistent way to name cells across embryos [12]. The physical arrangement, or adjacency matrix, of these cells is conserved and even the geometry, such as cell-cell contact areas, is broadly conserved, up to an overall size scaling [15]. The patterning paradigm of these embryos is one of immediate neighboring cells interacting through mechano-chemical mechanisms. The long-range patterning signals, which sculpt the dynamics of fly and frog embryos are believed to be largely absent, or at least, a higher-order effect that are sub-dominant to nearest-neighbor interactions [15]. Providing additional context, initially, a handful of maternal mRNAs are localized to the posterior pole of the zygote, defining the anterior-posterior (head-to-tail) axis. From this relatively unstructured start, and transcriptional inheritance from the individual’s mother, cells self-organize into distinct transcriptomic states, precisely arranged in space; there are 5 such states, referred to as cell-types, as early the 16 cell stage, Fig. 1B. The fate map of each cell, the set of terminal cell types its progeny will become, is restricted [12], Fig. 1A, thus many of these cells remain diverged in gene expression permanently. As development progresses, and the number of cells increases through cell divisions, additional cell types appear and increasing numbers of genes become zygotically expressed [16], again, all in a precise spatial arrangement. The temporal window during which the activity of the embryo becomes dominated by the action of zygotically transcribed genes is referred to as the Maternal-to-Zygotic Transition (MZT), the initiation of zygotic transcription in ascidian happens around the 8-cell stage and zygotic transcripts dominate around the 112-cell stage [17]. A multitude of molecular mechanisms are known or suspected to contribute to the embryo’s feat of self-organization including localized maternal mRNA, FGF signalling, *β*-catenin gradients, and asymmetric mRNA partitioning during cell-division. Major questions about the program of self-organization remain and many would-be insightful experiments, such as a complete space-time recording of live signalling, remain intractable. Yet measuring the gene expression, which is a proxy for the internal “state” of the cell [4], of every cell in an embryo is possible through single-cell sequencing. A central question addressed in this study is whether such measurements can be leveraged to study a broader spectrum of biological phenomena than identification of cell types. As such, the ascidian with its stereotyped cellular topology, geometry, and conserved lineages represents a simpler, but yet authentic, developmental system where general quantitative and physical approaches to identifying the collective modes of gene expression can be developed.

**FIG. 1.**
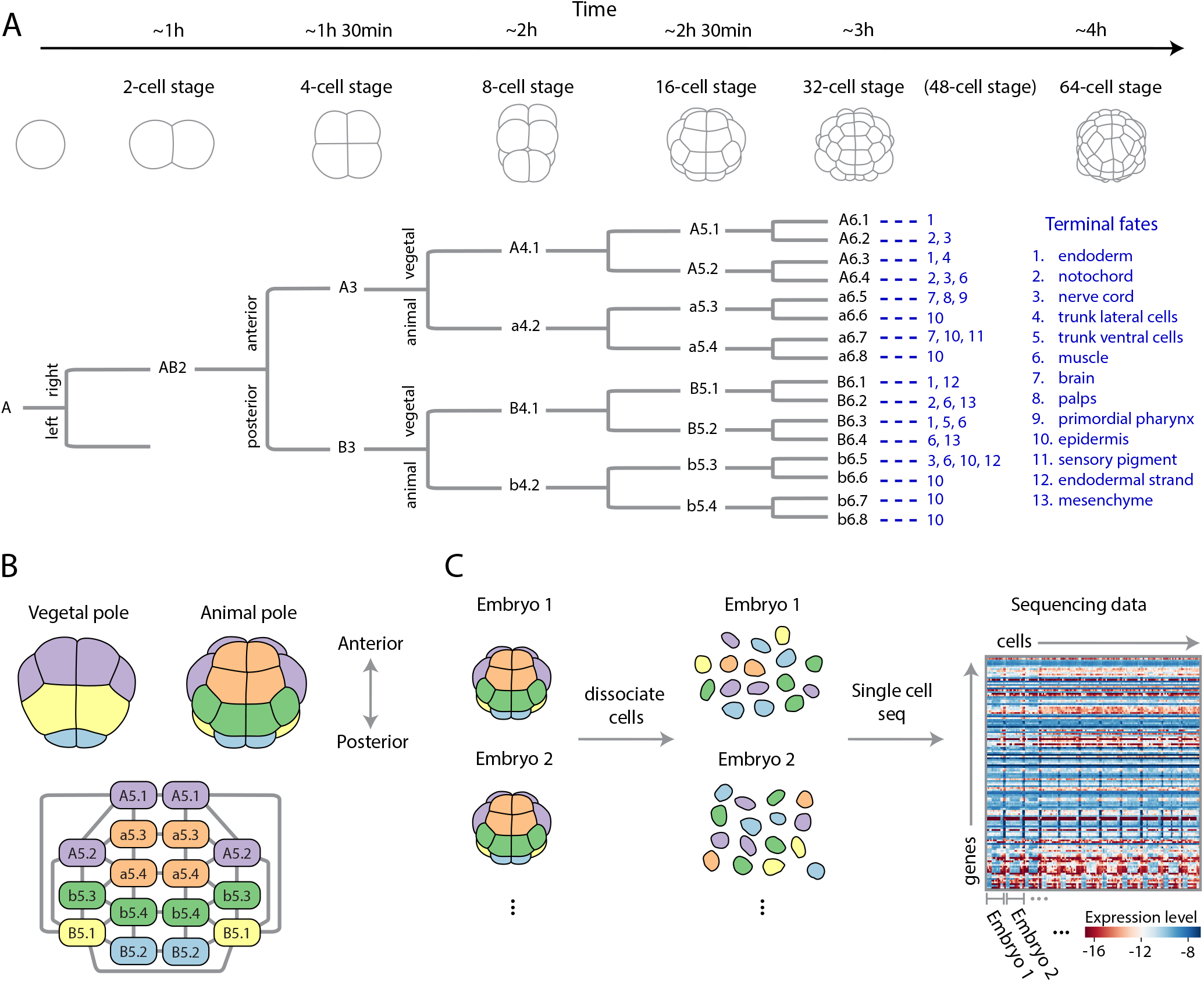
Developmental dynamics of the ascidian embryo: lineage, geometry, and single-cell gene expression data. (A) Early development of the ascidian *Phallusia mammillata*. Representative morphologies (top) shown along with right side of lineage tree (bottom) with the left being mirror symmetric. Cell fates at the larval stage indicated in blue for each 64-cell stage cell (adapted from Ref. [12]). Time is in hours post fertilization at 18^°^ [18]. (B) Representative cell geometry for the 16-cell stage (top), together with the cell-cell contact graph (bottom). Colors indicate transcriptomically distinct cell-states, as discovered in Fig. 2, which are also conserved across embryos. The animal-vegetal, anterior-posterior, and left-right axes form the 3 morphological axes of the embryo. (C) Experimental set up of Ref. [16], where large numbers of embryos at each cell-stage are first dissociated and then single-cell RNA sequencing at high-depth was performed. The precise identity of each cell is lost after dissociation, but the embryonic identity is retained through barcoding [16]. The resultant cell-by-gene expression matrix is shown for the 16-cell stage and for genes that are differentially expressed across the embryo (Methods).

### Data

In addition to the developmental considerations that make ascidian an attractive model system, there exists a publicly available and rich sequencing dataset [16]. Single-cell RNA sequencing was performed on individual *P. mammillata* embryos up to the 64-cell stage, with multiple biological replicates at each stage. Here, we will only consider the 8- to 64-cell stages. Notably, the sequencing platform used in Ref. [16] was Smart-Seq2 [19], which, together with relatively large cells, results in a sequencing depth higher than typical scRNA experiments, with on average 300 reads per gene at the 64-cell stage (Table S1). This depth of sequencing presents a unique opportunity to develop novel theoretical and computational frameworks.

The procedure of single-cell RNA sequencing creates some difficulty in asking what are essentially spatially minded questions that motivate our work. Before sequencing, embryos are dissociated into individual cells, Fig. 1C, and the location and identity of each cell is lost. Nevertheless, the cell type, and thus approximate spatial location, of each cell can be reconstructed from gene expression, as we will detail later (Fig. 2). Moreover, each embryo was barcoded, hence which cell belongs to which embryo is known, Fig. 1C, an essential requirement for identifying embryo-to-embryo variation. While a careful dissection and sorting of cells pre-sequencing is technically possible [20], it is tedious and impractical for more than a handful of embryos. Altogether, the combination of high-quality sequencing, a significant number of biological replicates (Methods), and a relatively constrained space of variation due to invariant cell number and geometry, make the ascidian an ideal model system for exploring collective variation in development.

**FIG. 2.**
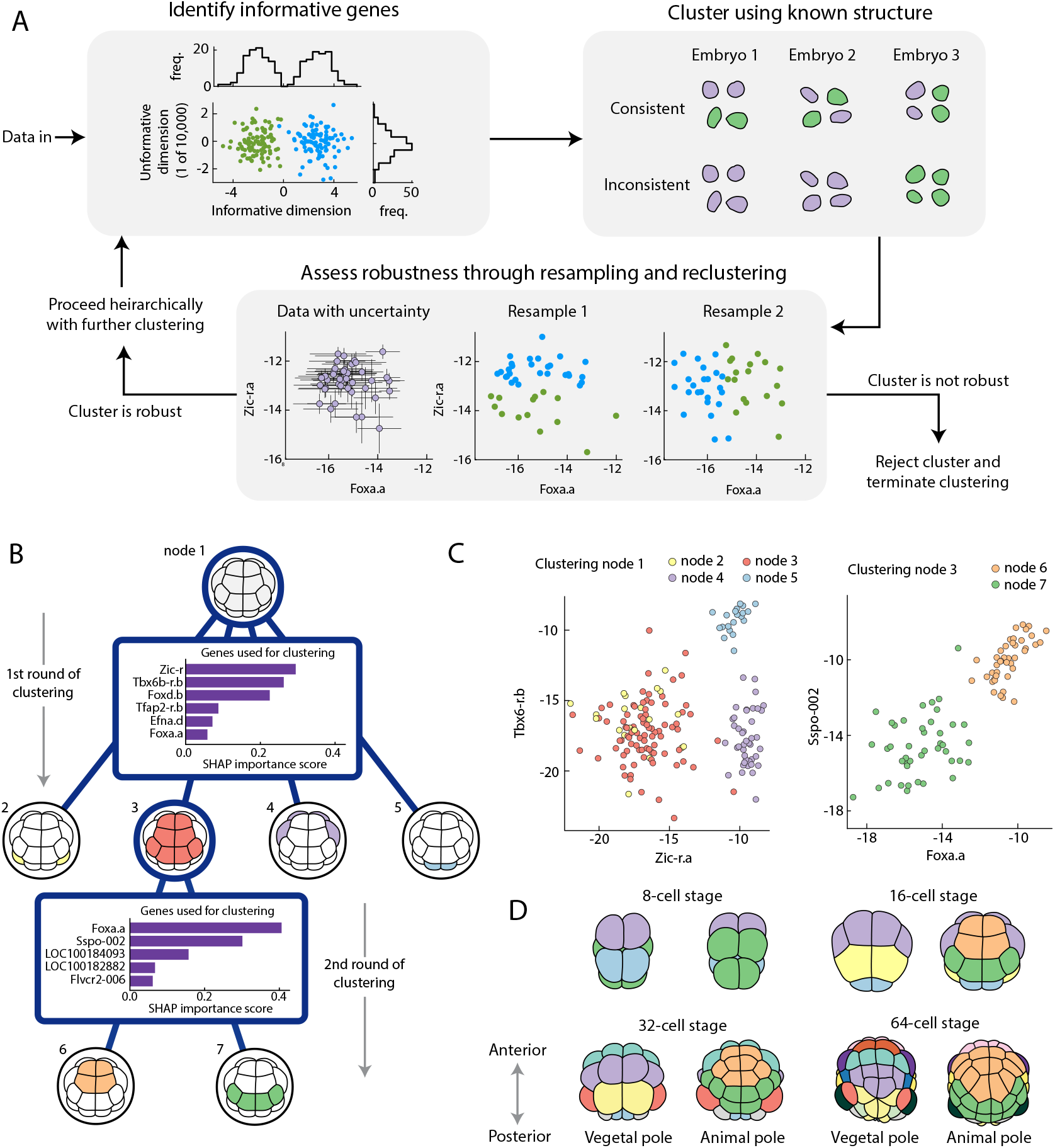
Principled and physically motivated clustering robustly recovers cell types. (A) Overview of clustering algorithm highlighting the three key underlying principles. 1. Identify informative genes. Informative dimensions split the data into separable clusters (green vs blue points, top left), uninformative dimensions do not, and make identifying clusters harder. 2. Cluster using known structural priors. A cell type should have equal numbers of cells in each embryo, a clustering without that property is inconsistent (purple and green cells represent clusters, top right). 3. Assess robustness through resampling and reclustering. We have a posterior for our expression levels [23] (purple points with error bars, bottom), and by repeatedly resampling and clustering (blue vs green points, bottom), we assess how consistent the clusters are and decide whether to keep or reject a cluster. For full details see SM Sec. III. (B) Worked example of the hierarchical clustering for the 16-cell stage. Starting from all cells at node 1, the algorithm splits into 4 clusters using the 6 genes shown. The SHAP importance score is an estimated measure of how useful each gene is when performing this round of clustering [30] (SM Sec. III). The algorithm then splits cluster 3 into two further clusters using a different set of 5 genes. The algorithm terminates here as further clusters are assessed to be not robust (SM Sec. III). (C) The two most important genes shown for both clustering stages, showing separation between identified clusters (node 2 and 3 differ in their expression of *Foxd*.*b*). (D) 8-cell through 64-cell stages colored by cell type, showing increasing transcriptomic specialization in time.

## III. RESULTS

### A robust statistical approach to cell type identification

Before any quantitative analysis, we need to preprocess the data. Starting from the raw transcripts, we align them to a reference genome (Methods), resulting in a cell-by-gene integer matrix of transcript counts, from which we infer the level of gene expression. Specifically, if *a*_*cg*_ is the expected number of mRNA molecules of gene *g* in cell *c*, then *α*_*cg*_ = *a*_*cg*_*/*∑ _*g*′_*a*_*cg*′_ is the fraction of mRNA molecules within that cell given to gene *g*, and we will refer to log *α*_*cg*_ as the “gene expression”, where the log accounts for variation in fold change rather than absolute value and serves to stabilize variance. There is uncertainty in this inference due to fluctuations in the number of mRNA molecules arising from inherent biological stochasticity, and from the measurement process, which can be significant. A host of data processing pipelines are used by the community, each one taking a different approach to normalizing the data in some manner to account for the uncertainty in the measurement [21, 22]. We normalize our gene expression data with Sanity [23], a biophysically motivated Bayesian approach to estimating a posterior distribution for gene expression. The resulting posterior is approximated as an independent Gaussian for each cell and gene, so the estimate for expression is 𝒩(*X*_*cg*_, *ϵ*_*cg*_), effectively giving error-bars to the inferred expression matrix *X*_*cg*_. Throughout, we make use of this uncertainty estimate, finding error bars for statistics, such as covariances, and for robustly identifying cell types. We have found that propagating the uncertainty to downstream analyses greatly strengthens the robustness of our statistical inferences and scientific conclusions.

A major technical challenge is to find the distinct transcriptomic states, or cell types, at each stage and identify which state each cell belongs to. Identification of cell types allows us to partially reconstruct the embryo’s spatial structure; an accurate classification of each cell is crucial to compute quantities like mean expression levels and cell-cell covariances around that mean. Despite this being a relatively mature field, directly applying standard clustering approaches to the expression data does not consistently recover cell types (SM Fig. S14). In order to robustly recover clusters, we developed an algorithm, Fig. 2A, which accounts for known properties of the developmental system in focus, listed as follows. Firstly, most genes are not expressed in an informative way [24], either not expressed at all, or expressed equally in every cell in the embryo. A few genes will be expressed in some cell types but not others, and finding these informative genes is crucial to distinguishing clusters, this information gets lost when using all genes, Fig. 2A. At each stage of clustering, we adapt the algorithm of Ref. [24] to find a set of informative genes by generating trial clusters and searching for genes that discriminate between these trial clusters (SM Sec. III). Secondly, clusters must be consistent across embryos, a cluster with 2 cells in every embryo is conceivably a cell type, a cluster with 4 cells in one embryo and zero cells in the others is not a cell type, Fig. 2A. Throughout, we enforce all clusters to be consistent with the known embryo structure (SM Sec. III). Thirdly, we use uncertainty about expression to assess whether a cluster is robust [25, 26] and is to be kept or not. If a cluster is robust, then that cluster should be consistently identified after perturbing the data and reclustering. Hence, upon repeatedly resampling from the posterior, subsampling the data, and clustering, we should repeatedly find the same clusters, Fig. 2A. Having performed this resampling, we compute a cluster consistency score, which we then compare against cluster consistency scores for null data sets generated by randomly assigning cells to embryos. These null data sets preserve the variance in the expression but destroy the known structure of cell types, a true cluster should have a significantly better consistency score than these null data sets, and this provides us a criteria to assess whether a cluster is robust and should be rejected or not (SM Sec. III). Overall, we combine these three principles into a hierarchical algorithm, motivated by the hierarchical nature of cell differentiation [27]. We search for a small number of well separated clusters before searching these for further subclusters, potentially using a different set of genes to do so. The 16-cell stage is shown as an example in Fig. 2B-C, for full details of the algorithm and results for other stages see SM Sec. III. Having found the clusters, it is straight-forward to identify them with cell identities and their position in the embryo, Fig. 2D, using existing *in situ* hybridization data for select genes [28] (SM Sec. III). We recover cell types previously identified [16, 29], but note that the challenge is not only to identify cell types but to accurately assign a cell type for every single cell including those with seemingly ambiguous expression levels. Altogether, the approach outlined here and described in detail in the SM can be used directly for systems with invariant cell lineage, such as ascidian, and many of the ideas can be adapted broadly for a robust and principled approach to cell type identification. The Bayesian approach to normalization as well as the resampling and shuffling inherent to our method could make it computationally expensive for large datasets, adapting the algorithm to be feasible for large datasets is an outstanding challenge worth overcoming.

### Identifying the dimensionality of phenotypic variation in the embryonic transcriptome

With cell types recovered, we can combine embryos to get an average transcriptomic state for the embryo at each cell stage. This typical state, or wild type (WT), contains information about the system. For instance, measuring activation of the *Otx* gene in a specific cell at the 32-cell stage tells us that cell has received an FGF signal emitted by a neighbouring cell. Probing this usually involves making a significant perturbation, e.g. genetically knocking out or pharmacologically perturbing a signalling pathway, and studying the result which may differ significantly from the WT, Fig. 3A. Here, complementary to this approach, we study the system using only the average WT trajectory and the natural phenotypic variation around it, Fig. 3B. Though cells occupy a vast transcriptomic space spanned by the number of genes in the organism, *n*_*genes*_, embryos occupy an even larger transcriptomic space, *n*_*cells*_ × *n*_*genes*_. This is the dimensionality of the embryonic transcriptome that we intend to build models for and study quantitatively. At a particular developmental time along the one-dimensional (parameterized by time) WT trajectory, embryos have a *n*_*cells ×*_ *n*_*genes*_ − 1 dimensional space to explore, Fig. 3B. We refer to this form of variation in gene expression as natural since each individual will deviate from the consensus trajectory due to genetic and/or environmental causes. Physics has a rich history of studying variation and using it for inference, from correlation functions and phase transitions, to fluctuation and dissipation. Here, we will study the subspace that is accessible to typical biological variation and, from quantitative analyses and physical modeling, infer interactions between cells.

**FIG. 3.**
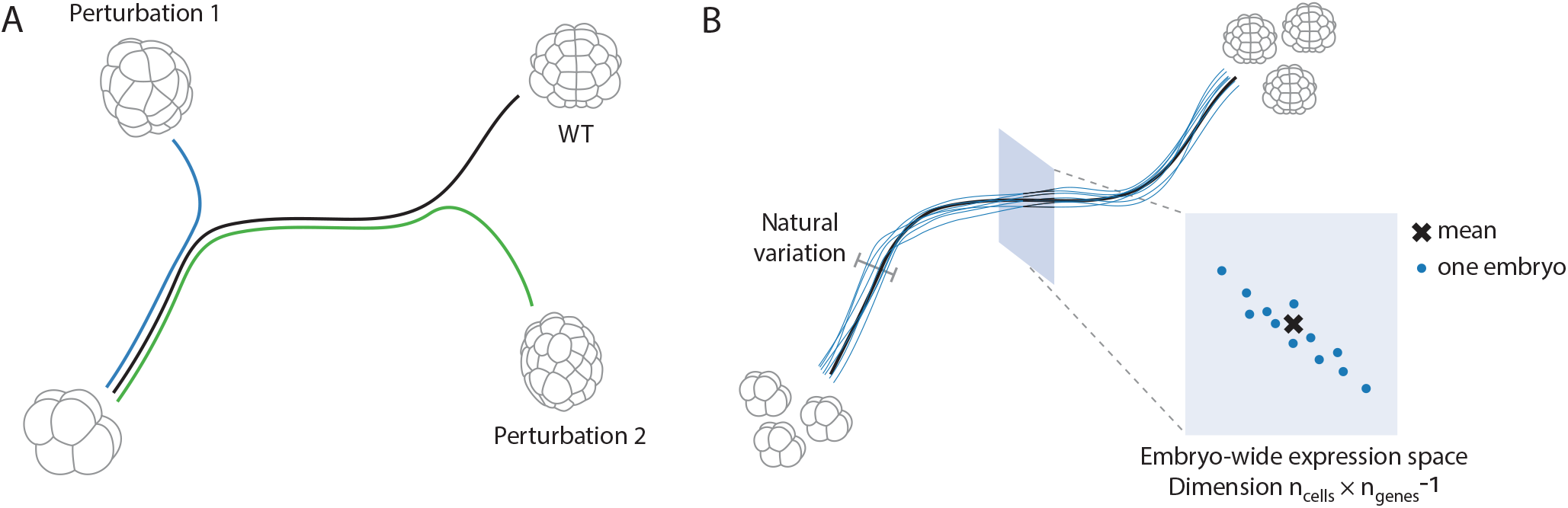
Studying the mechanisms underlying collective modes of gene expression through quantifying natural phenotypic variation. (A) Development is thought to be canalised, implying that embryos robustly follow a typical wild type trajectory (black). Typically, the tools of molecular biology used to study development produce perturbations that are strong enough to kick the system far from it’s canalised trajectory. (B) Even though development is canalised, each embryo will follow a unique trajectory, slightly deviating from the average. Taking a slice at a specific time point along the developmental trajectory, there exists a high dimensional space of variation that embryos can explore.

It is not obvious that the scRNA-seq measurement reveals any form of natural variation at all, perhaps all that can be garnered from such measurements are cell types. Said another way, is there any signal above the noise after cell types are identified? Can we indeed study natural variation and the emergent patterns manifest in space?

Turning again to the experimental data, prior to any modeling, we first ask whether we can even detect signatures of collective variation within an embryo. As a point of comparison, and motivated by the biology at hand, at each cell stage we consider two sets of genes: (1) maternal transcripts, which are present in the zygote but are not zygotically transcribed at the 64-cell stage or earlier (we exclude the handful of maternal transcripts that are localized to the posterior pole which determine the germ cell lineage) (2) zygotic transcripts which are not present in the zygote, but are expressed at subsequent stages of development (Methods). Each of these genetic classes has a corresponding cell-by-gene matrix of expression, *X*_*cg*_. To see whether there is any collective signal in this matrix, we examine it’s singular values, Fig. 4A-B, having subtracted the mean expression levels, *X*_*cg*_ ↦*Y*_*cg*_ = *X*_*cg*_ −∑_*d*_ *X*_*dg*_*/*∑ _*d*_ 1. Before we can evaluate the singular values for signal, we must account for various features of the biology that give rise to variation distinct from the collective modes of phenotypic variation we wish to identify. For the maternal genes, there will be variation arising from mother-to-mother differences in mRNA deposition, which we can subtract from the expression matrix, *Y*_*cg*_ ↦ *Z*_*cg*_ = *Y*_*cg*_ *−* ν_*m*(*c*)*g*_, where ν_*mg*_ is the mother specific mean expression and *m*(*c*) is the mother of cell *c*. Further, we anticipate additional embryo-to-embryo variation, which we subtract similarly, *Z*_*cg*_ *Z*_*cg*_ *− η*_*e*(*c*)*g*_, where *η*_*eg*_ is the embryo specific mean expression and *e*(*c*) is the embryo containing cell *c* (SM Sec. IV). For the zygotic genes, we know that each cell type, whose identification we have described in the previous section, has specific expression patterns, which we must remove, *Y*_*cg*_ ↦ *Z*_*cg*_ = *Y*_*cg*_ *− µ*_*t*(*c*)*g*_, where *µ*_*tg*_ is the cell type specific mean and *t*(*c*) is the cell type of cell *c*. We also expect that each embryo differs in developmental time or how far the embryo is along the WT trajectory [3], even though care was taken to stage them at a consistent time [16]. As we want to study variation around the WT trajectory at a fixed time, not variation along the WT trajectory itself, we remove the temporal variation by projecting onto one fixed developmental time via fitting an embryo specific time *τ*_*e*_, and a local cell-type specific tangent vector to the developmental trajectory *α*_*tg*_ by minimizing the variance left in the matrix *Z*_*cg*_ ↦ *Z*_*cg*_ − *τ*_*e*(*c*)_*α*_*t*(*c*)*g*_, Fig. 4C (SM Sec. IV).

**FIG. 4.**
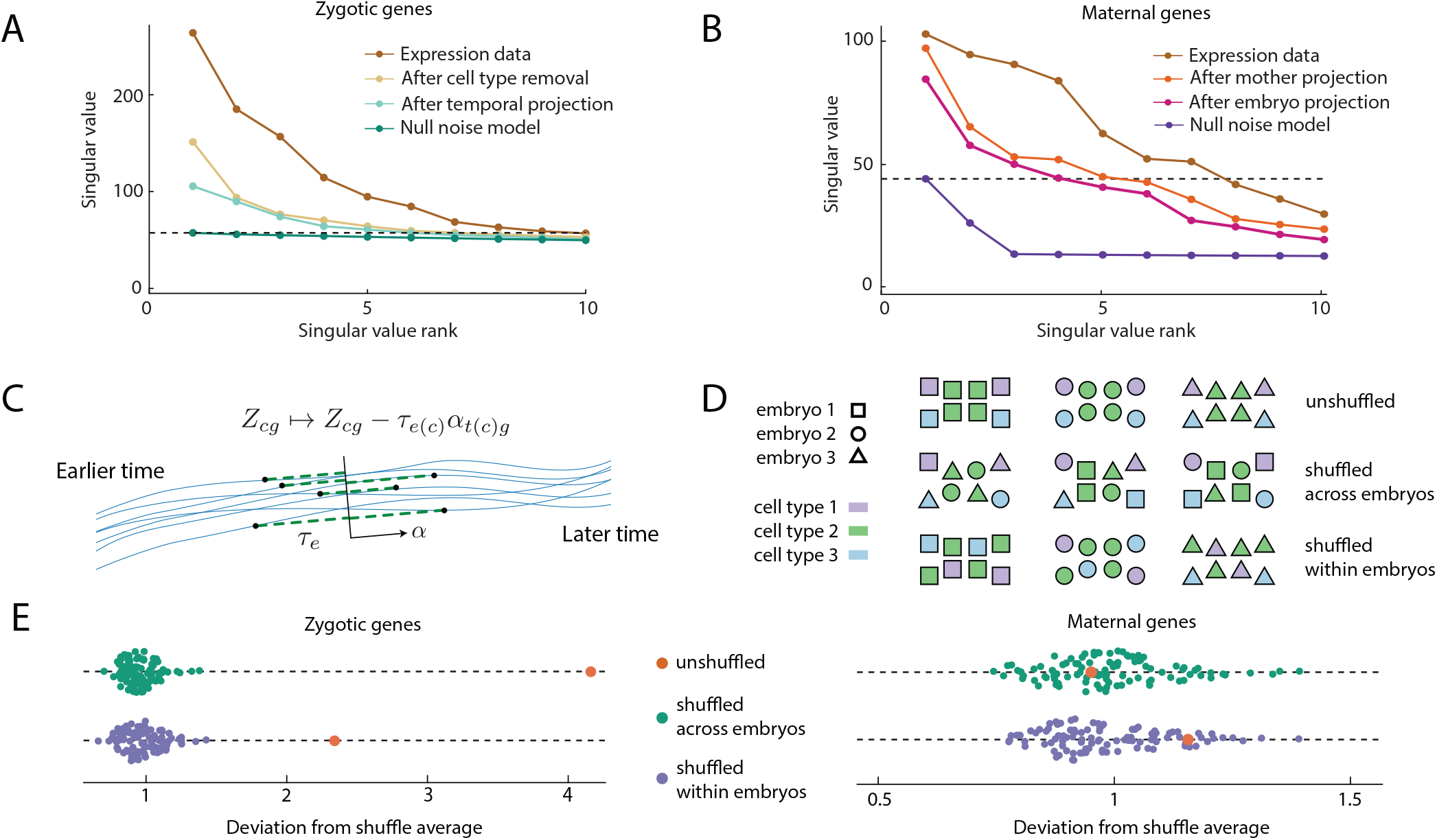
A quantitative assessment of single-cell gene expression data demonstrates correlated expression between cells within an individual embryo. (A, B) Singular values of the gene expression matrix, for zygotic (A) and maternal (B) genes at the 32-cell stage before and after known sources of variation, such as cell types, variations from mother-to-mother or embryo-to-embryo, and variations in developmental staging (see C), is removed. To demonstrate that significant variation remains, a null noise model is shown for comparison. The horizontal dotted line shows the largest singular value of the null statistical model and provides an estimate for how many singular components to keep (SM Sec. IV). Other stages are shown in SM Fig. S15. (C) Local developmental time, *τ*_*e*_ for each embryo (black points, full trajectories are blue lines) will vary owing to slight discrepancies in staging. We remove this variation by projecting each embryo onto a common time (black line) by identifying the axis tangent to the average trajectory, *α*, and then subtracting the temporal variation, *τ*_*e*_*α* from the gene expression matrix *Z*. (D) To test for cell-cell correlations within an embryo we shuffle cells of the same cell type between embryos, as well as shuffling cells within an embryo to test whether such correlations have spatial structure. Color is cell type, shape is embryo identification. (E) Results of shuffling within and across embryos as in (D) for both maternal and zygotic genes at the 32 cell stage (other stages in Fig. S17). Each parameter configuration has 100 random shuffle trials. In every case, all covariances were measured and the deviation Δ from the average covariance values is plotted, note that while the covariance vector remains the same for the unshuffled covariance matrix (orange) it has different deviation for within and across embryo shuffles as it is being compared to a different average. Shuffling is not significant for maternal genes, but significantly affects zygotic genes, indicating collective signal across and within embryos.

Having accounted for these known sources of variation, we are in a position to address whether the maternal and zygotic matrices have any remaining collective signal, or whether it can be explained by each entry being an independent random variable. To do so, we compare against a null distribution constructed by permuting the rows of the matrix, preserving the marginal distribution of each gene, but destroying the correlation structure. For the zygotic genes at the 16-cell stage we find that all the variation could be explained by the null distribution, although for 32- and 64-cell stages this is not the case; the data has non-trivial singular values, and hence signal, Fig. 4A-B, (SM Sec. IV). For the zygotic genes, the cell-type specific expression accounts for much of the variance with the WT trajectory projection typically accounting for the largest remaining singular value, Fig. 4A. For the maternal genes, we find non-trivial singular values at each stage. Additionally, the null model informs us how many singular values we should keep, the rest plausibly being noise [31] (SM Sec. IV). We project the expression levels for each cell onto these principal components (PCs), effectively setting to zero the singular values that were plausibly noise, and work in this low dimensional space from here onwards.

We have not yet shown that genuine spatially-ordered collective embryo-to-embryo variation is present, since correlations between genes in the same cell could give a matrix with significant singular values. To test this, we compute covariances for cells within the same embryo and examine how this changes after shuffling cells between embryos, Fig. 4D. Note that we cannot compute the covariance between, say, neighboring a5.3 and a5.4 cells since we only know that a cell is in the a5.3-a5.4 cell type, not its exact location, Fig. 1B. Yet, we can still compute the average covariance between a cell in the a5.3-a5.4 cell type and one in the B5.1 cell type, within the same embryo. Specifically, we can measure the covariance

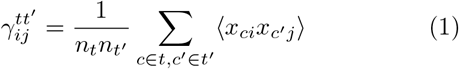

for cell types *t, t*^*′*^, principal components *i, j*, where *n*_*t*_ is the number of cells in cell type *t*, and *x*_*ci*_ is the expression after known variance has been removed. We can also measure covariance within a cell for each cell type,

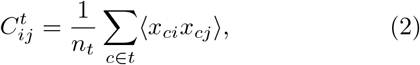

Together, we say these make up the set of empirical covariances {*C*_*α*_}. For a robust empirical estimate of 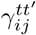, and 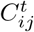, we repeatedly resample expression from the posterior to account for uncertainty from the expression measurement, as well as bootstrapping to account for uncertainty from having finite data, see SM Sec. IV for details.

To test for collective variation, we compute the inter-cell covariance statistics, 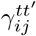, before and after shuffling the data, Fig. 4D. Taking 100 shuffles as well as the un-shuffled data, we compute the component wise average covariance, 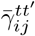, as well as the component wise standard deviation, 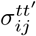, from which we calculate the deviation Δ,

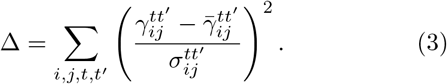

If the value of Δ for the unshuffled data is significant compared with values for the shuffled data, then there is significant collective variation, see SM Sec. IV for full details. For maternal genes, shuffling within an embryo does not affect the covariance statistics at any stage, Δ is not significant, suggesting a lack of spatial structure, Fig. 4E. Further, shuffling across embryos is not significant after embryo-to-embryo variation is removed. Hence, the variation can be explained by embryo-to-embryo variations in initial mRNA deposition, without needing to invoke spatial distributions or coupling through cell divisions. The same is not true for the zygotic genes, where cell types and temporal projection are insufficient to explain the within embryo covariance, the shuffle tests reveal genuine collective variation in gene expression within and across embryos, Fig. 4E (see SM Fig. S16 for all stages). This agrees with our biological intuition, that the space of zygotic transcripts is tightly controlled and thus the fluctuations that do occur are coupled across cells within an embryo, in contrast to the maternal transcripts which can vary from cell to cell within an embryo and are not as tightly restricted in their variation. Altogether, our detailed statistical approach set out to ask whether there was measurable collective fluctuations having accounted for known sources of variation. We find that for the set of zygotic genes there exists embryo-wide variation with spatial structure.

### Statistical physics modeling of phenotypic variation

So far, we have established the existence of collective fluctuations within an embryo for zygotic genes, in contrast to the maternal transcripts. We now attempt to understand the nature of this collective behavior. Constructing and then studying the empirical joint probability distribution of an embryo’s state of gene expression, the embryonic transcriptome, is impractical, since each embryo counts as only one data point in a high-dimensional space and we only partially know a cell’s identity. Despite these limitations, statistical physics gives us a way to rationalize statistics of this system, making minimal assumptions about the nature of interactions and using the statistics we can estimate, in particular the covariances between different cell types. In short, we suppose there are certain important statistics which are prescribed by the system. For instance, the variance of a gene within a cell is constrained by the gene regulatory network, or the covariance between two sister-cells is fixed by the mechanism of division. There are infinitely many distributions that are consistent with these statistics, but out of all such distributions there is one that is makes minimal further assumptions: the maximum entropy distribution [32–34]. Here, the first non-trivial statistics we can consider to be constrained by the system are covariances (including variances), as we have already subtracted the mean.

For instance, we could take the average covariance between PC *i* and PC *j* for cells in cell type *t*,

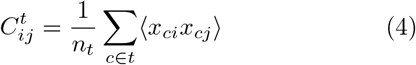

to be fixed by the system, where *n*_*t*_ is the number of cells in cell type *t*. Similarly, we could consider the covariance between cells *p* and *q*, Λ_(*pi*)(*qj*)_ = ⟨*x*_*pi*_*x*_*qj*_⟩ to be fixed. Since there is a left-right symmetry in the embryo at this early stage, there is no biological reason to vary the left and right covariances independently. Instead if *p* and *q* are on the left side of the embryo, with 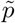 and 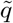 their equivalent cells on the right side, then we may fix 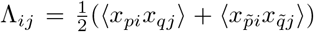. In general, we can introduce an asymmetric directed adjacency matrix, Ω_*rs*_, to fix covariances of the form

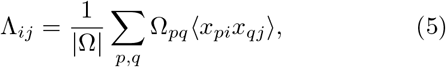

with Ω_*sr*_ ∈ {0, 1} keeping track of which cells are being included.

Since ⟨*x*_*pi*_*x*_*qj*_⟩ ≠ ⟨*x*_*pj*_*x*_*qi*_⟩, covariances between cells can be directed, for example, the covariance between PC 1 in A5.1 and PC 2 in A5.2 can be different than the covariance between PC 2 in A5.1 and PC 1 in A5.2. Indeed, we might directly fix the covariance between PC 1 in A5.1 and PC 2 in A5.2, but not fix the covariance between PC 2 in A5.1 and PC 1 in A5.2, implemented by having an asymmetric adjacency matrix Ω. However, in certain cases we might explicitly fix symmetric covariances, for instance the covariance between PC *i* in one cell and PC *j* in a neighboring cell, averaged over all pairs of neighbors without choosing a directional arrow between these pairs. In this case we denote the adjacency matrix as Ψ to explicitly indicate that it is symmetric. In general there will be a set of such symmetric constraints and a correponding set {Ψ^*k*^} of symmetric adjacency matrices, along with a set of asymmetric constraints and corresponding set {Ω^*k*^} of directed adjacency matrices that we consider to be fixed.

Whatever set of covariances we elect to fix, the distribution with maximum entropy that is consistent with those covariances is a Heisenberg-like model, *p*(***x***) = 𝒩 (0, *J*^−1^) with ***x*** = (*x*_*pi*_) with *i* indexing over principal components, *p* indexing over cells within an embryo, ***x*** is flattened into a vector, and

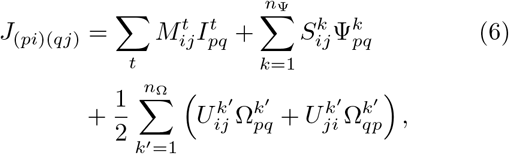

where 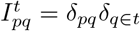 is an indicator function with *t* a cell type, Ψ^*k*^ and Ω^*k*^ are adjacency matrices of which there are *n*_Ψ_ and *n*_Ω_ respectively, and *M, S, U* are interaction parameters; *M* couples genes within the same cell, *S* symmetrically couples different cells, *U* is a general coupling between different cells, all arise as Lagrange multipliers enforcing the constraints (SM Sec. V). This distribution arises as it is the simplest one consistent with biologically and physically motivated constraints on the system, namely that principal components within the same cell are coupled with a cell-type specific coupling, and that principal components between cells can be coupled through an adjacency matrix.

As an example, suppose the developmental dynamics directly sets the intra-cell covariance matrix for each cell type, as well as the covariance matrix for cells in spatial contact, Fig. 5A. The interaction matrix, 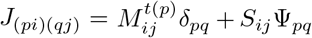 is sparse, where Ψ_*pq*_ is the spatial adjacency matrix of an embryo at a given stage, Fig. 5B. Yet the covariance matrix, *J*^−1^ is dense, Fig. 5C, as is the matrix of observed covariances between cell types Fig. 5D. The only assumption is that cells in spatial contact have correlated gene expression, the form of the distribution follows from the maximum entropy principle and the parameters are fit to experimental measurements of covariances (SM Sec. V). We can ask how well this simple model explains the data, and compare against a model which assumes sisters are interacting, so Ψ_*pq*_ = 1 when *p* and *q* are sister cells, or a model where all cells are equally interacting, so Ψ_*pq*_ = 1 for any pair *p* ≠ *q*, Fig. 6A. We quantify the quality of the fit by comparing the empirical covariances, {*C*_*α*_}, the set of covariances measured from the data, to the model predicted covariances 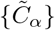, which we can directly compute from the model (Fig. 5D). A perfectly fitting model would have 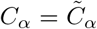, ∀*α*, and we quantify the fit by measuring the variance in {*C*_*α*_} explained by 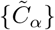, namely the quantity 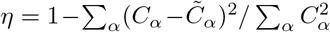. At the 32-cell stages, the embryo-wide model and neighbor model can reasonably explain the experimental data (*η* = 0.89 and 0.87 respectively), Fig. 6B, with the spatial model fitting the data better at the 64 cell stage (*η* = 0.79 embryo-wide vs 0.92 for spatial). In contrast, the sister model can not explain the experimental observations, Fig. 6B, as this model can never cause embryo-wide correlations, only correlations between sister cells (*η <* 0.63).

**FIG. 5.**
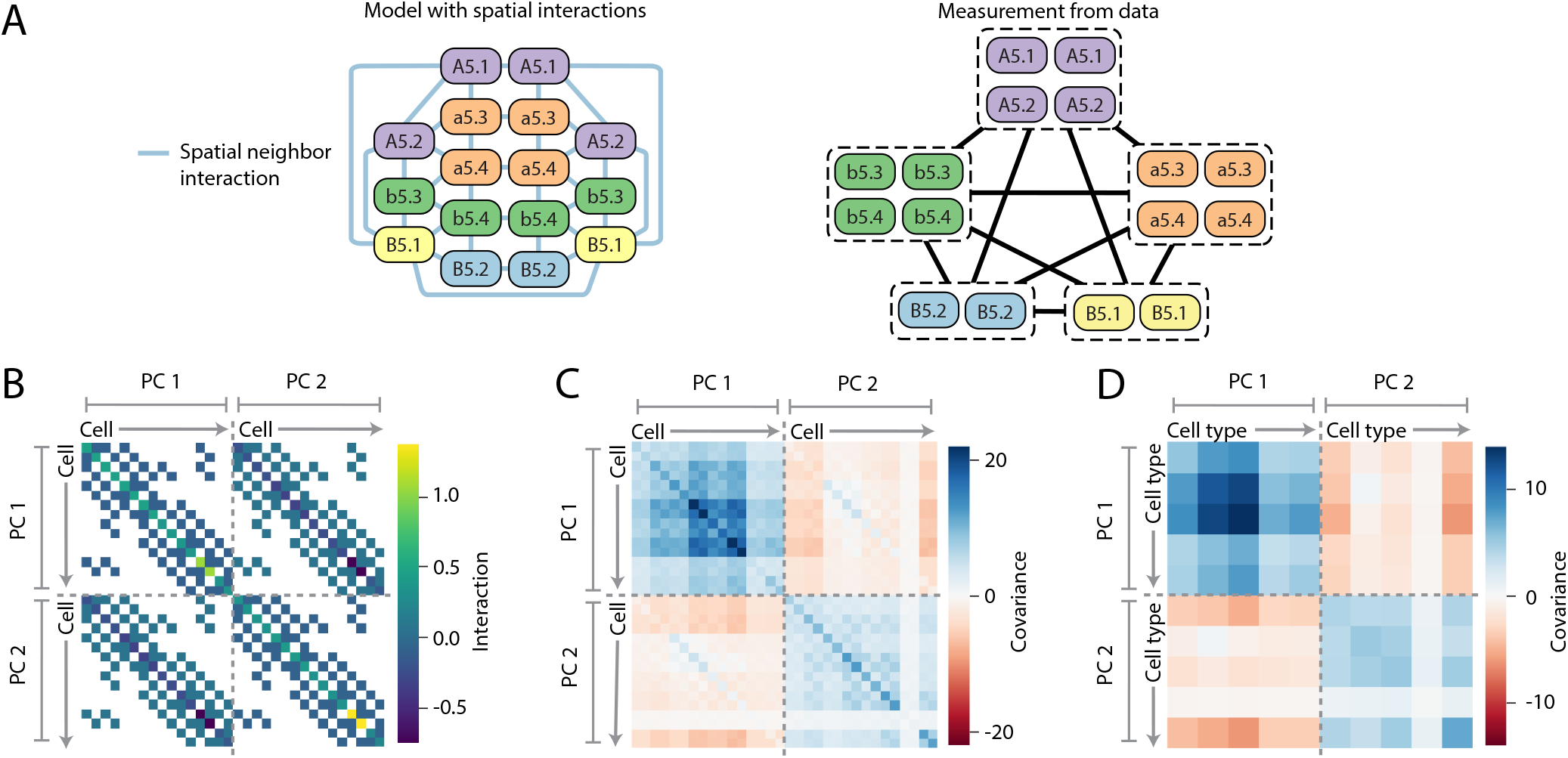
Structure of statistical physics model with sparse interactions. (A) Model of the 16-cell stage, where spatial neighbors are directly interacting. In the model we know every cells identity and the covariance between any two cells (left). However, in the data we only know the cell type of each cell and not it’s exact identity (right), hence we can only compute the covariances between cell types. (B) The sparse interaction matrix *J*, for model *p*(*x*) = 𝒩(0, *J*^−1^) where only spatial neighbors have direct interactions, shown for the 16-cell stage with 2 principle components as an example. (C) Dense covariance matrix *J*^−1^. Note that while *J*^−1^ is always symmetric, (*J*^−1^)_(*pi*)(*qj*)_ ≠ (*J*^−1^)_(*pj*)(*qi*)_ in general, even if *J*_(*pi*)(*qj*)_ = *J*_(*pj*)(*qi*)_. (D) Dense matrix of cell type to cell type covariances, corresponding to the model *J*, which, unlike (C), can be directly compared to measurements from data.

**FIG. 6.**
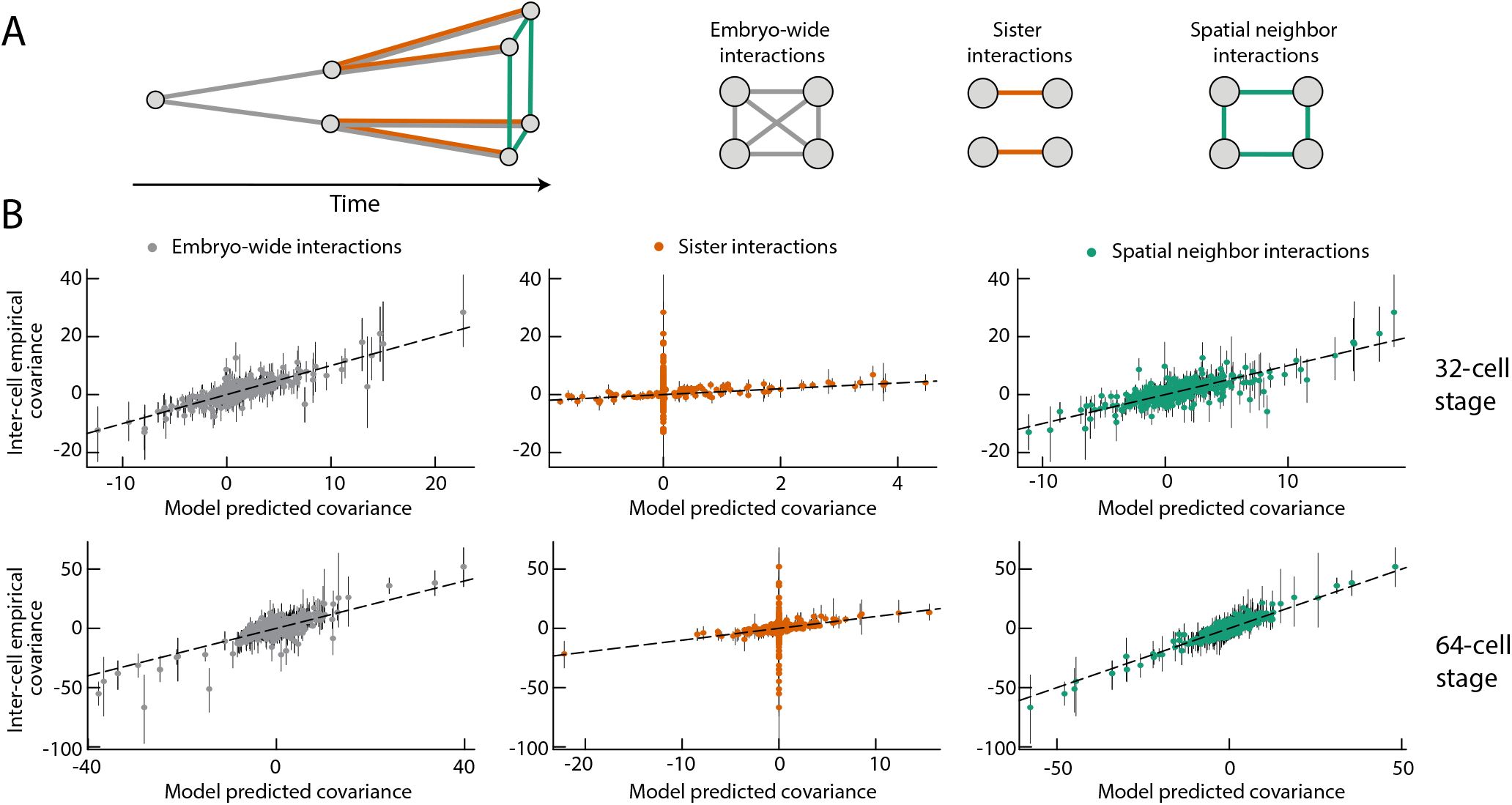
Comparing models with data identifies plausible families of statistical physical models for interactions. (A) Correlations between different cells could arise from signaling resulting in interactions between spatial neighbors, cell division resulting in interactions between sister cells, or variations in the zygote resulting in all-to-all coupling across the embryo. We consider how well these different hypotheses explain the experimental data. (B) The inter-cell empirical covariances (IEC) are covariances between different cell types measured from data. The model predicted covariances (MPC), are covariances between different cell types in the maximum entropy model (Fig. 5D). A model which explains the data should have MPC values equal to the IEC values. Here, we test models which assume that all cells within an embryo directly interact (gray) only sisters interact (orange), or only spatial neighbors interact (green). Both the embryo-wide and spatial model are able to reasonably explain the data, with points lying around *x* = *y* (black dashed line), whereas sister-cell interactions alone are not sufficient. Without interactions between non-sister cells, many covariances are zero in the model despite being non-zero in the data. Error bars are from the 25^*th*^ to 75^*th*^ percentile of covariance estimates.

While comparing these hypotheses for interactions is revealing, there are embryo-wide, neighborhood, and sister cell interactions at play, not to mention interactions between specific cells representing cell-cell signalling. Consider a model which admits embryo-wide, spatial neighbor, and sister cell interactions, together with specific cell-cell interaction terms which respect the left-right symmetry of the embryo for all spatial neighbors. Such a model has too many parameters (1080 *U* and *S* values for the 32-cell stage with 5 principal components) and will overfit the data. Therefore, and motivated by the biology of the system, we assume the nature of interactions are sparse and use regularization [35, 36], together with uncertainty estimates for the covariances, to identify which of the many possible interactions are important and can robustly explain the experimental measurements. Specifically, we apply a *L*_1_ regularization penalty for each interaction parameter (*U* and *S* terms) with strength *λ*. We draw *N* = 100 empirical covariance matrices from the posterior distribution, and fit a regularized model to each of them. We then assess which parameters are a consistent and non-zero size across different fits, and use this to rank terms by importance. We sample across a number of *λ* values, making a ranking for each, and then aggregate the rankings with the Schulze method [37], resulting in an overall ranking of terms by importance. Full details in SM Sec. VI. Given this ranking, starting from the most important terms, we include sufficient terms in the model to reach *η >* 0.9, finding a 17 term model at the 32-cell stage, Fig. 7B, and a 12 term model at the 64-cell stage (SM Fig. S19).

**FIG. 7.**
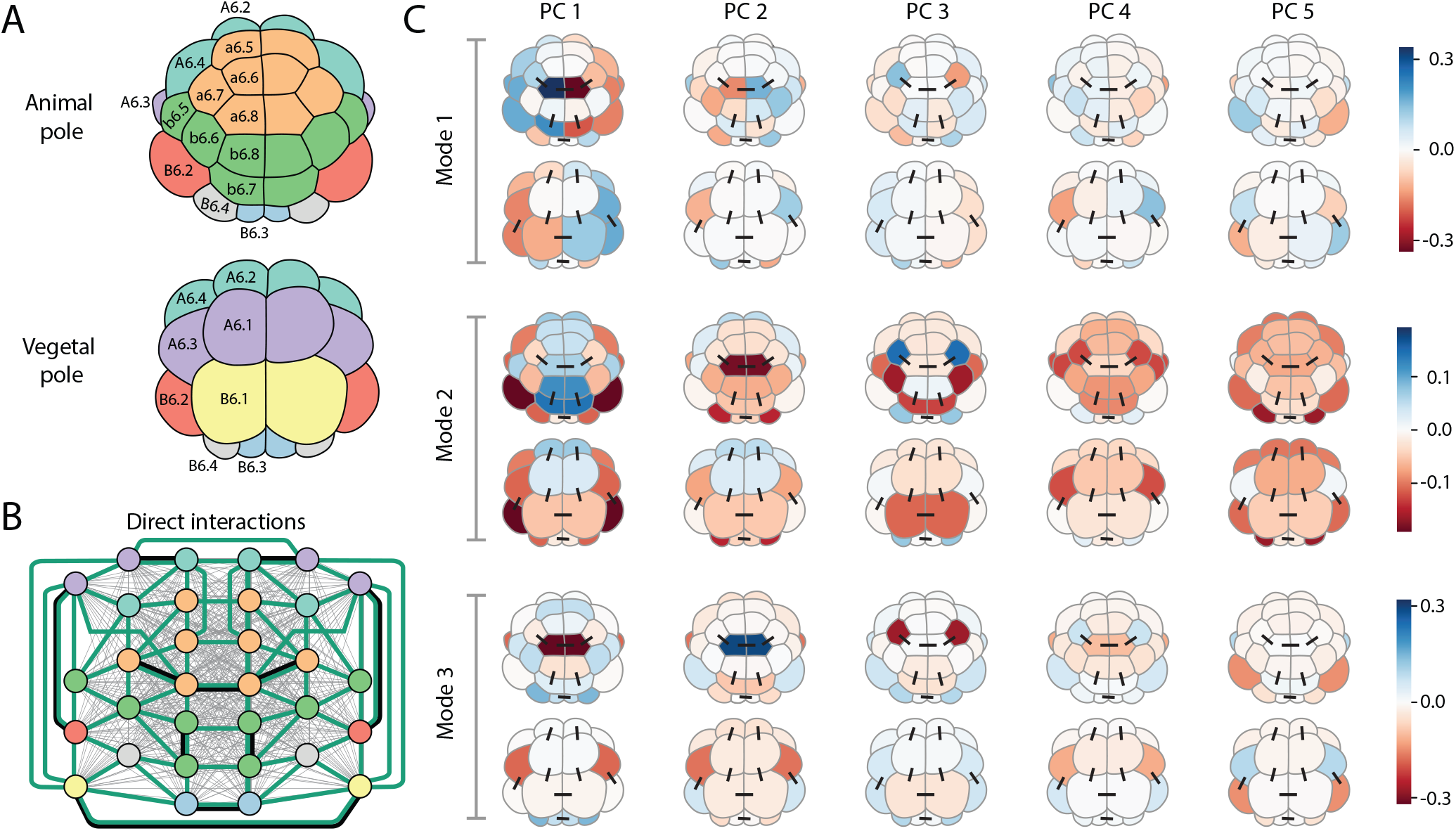
Statistical physics modeling identifies spatially ordered collective modes in expression data and reveals underlying interactions. (A) Schematic of the 32-cell stage embryo colored by cell type. (B) Graph representation of the 32-cell stage embryo, colored by cell type with edges representing inferred direct interactions in regularized statistical model. Embryo-wide (grey lines) and neighbor (green) cell-cell interactions are present in model, as well as specific cell-cell interactions (black lines). A full ranking of terms is in Table S3. (C) Principal modes of variation from inferred regularized model. The inferred model from (B) is a Gaussian and hence the distribution can be interpreted as a multidimensional ellipsoid defined by the covariance matrix *J*^−1^. Here, the first three principal directions of that covariance matrix (called Mode 1-3) are shown, which together capture over 80% of the variance of the model. Each such direction, or mode *ϕ*_*cg*_, has both spatial and PC components and is visually represented by five columns (PC’s = 1, …, 5) where column *g* shows an embryo (both animal and vegetal view shown) with cell *c* colored by the value *ϕ*_*cg*_. Black lines show direct interactions between specific cells. The first mode represents a left-right mode of variation, whereas the second mode appears to somewhat correspond to an animal-vegetal mode.

The interactions we identify must be cross-referenced with what is known and suspected about the signaling interactions at this stage of development. These interactions have been studied through the tools of molecular embryology, which are wholly distinct from the approach we have pursued here. Neural induction, where FGF signalling causes a6.5 and b6.5 cells to adopt a neural fate, occurs at the 32-cell stage and is the first known cell-cell signalling that causes a change of state. However, levels of the FGF-target gene *Otx* were not found to be elevated in potential a6.5 or b6.5 cells in the sequencing data, suggesting that the sequenced cells were dispersed before the increase in *Otx* expression, and hence before induction occurred. In general, cell-cell communication induces correlations in time, with the transcriptomic machinery taking a finite time to respond to new information. A static snapshot may capture some of those temporal couplings, but only after the signal is received. Neighbor, embryo-wide, and a handful of specific cell-cell interactions, three of which are between sister cells reflecting interaction through inheritance, appear in the top inferred terms (Table. S3). All of the inferred specific cell-cell interactions are between animal cells or between vegetal cells with no animal-vegetal interactions which indeed should not have occurred yet. For the remainder of non-sister inferred interactions, there is no known molecular mechanism for them, say left and right B6.1 cells, to communicate at this stage in a way analogous to FGF signaling. Instead, perhaps they could reflect embryo-to-embryo genetic differences in levels of expression for expressed genes, such as *nodal* or *snail* in the case of B6.1, which would correlate expression between these cells across embryos. While cell-cell inductions have occurred by the time of the 64-cell sequencing, the data has fewer and less complete embryos, as well as noisier sequencing which limits our ability to robustly infer interactions. Seemingly, embryo-wide and neighbor interactions, in addition to a coupling between the germ cells, are sufficient to explain the measured collective variations at the 64-cell stage, the rich landscape of specific cell-cell interactions made inaccessible by limitations in the data. Altogether, at the 32-cell stage, as the data is pre-*Otx* expression unsurprisingly we do not see interactions corresponding to FGF signaling, but we do infer some embryo-wide, neighbor and specific sister cell interactions. At the 64-cell stage the lack of data prevents robust inference of cell-cell interactions.

Whilst we do not know the ground truth of cell-cell interactions, in addition to the results being biologically plausible, we can check to see whether our inference procedure is self-consistent. Specifically, given a particular sparse model, we can sample from it to create a data set of the same size and quality as the experimental data (SM Sec. VII). After putting this simulated data through the inference procedure, we do not recover exactly the same ranking, but true sparse interactions are identified as amongst the most important (SM Fig. S21), showing that there is robustness to the inference pipeline despite the limitations of the data.

### Identifying the collective modes of gene expression

Having found a model with a sparse set of interactions, which explains the observed covariances, we can examine this model and ask what the spatially ordered modes of collective variation look like in this system. Given any Gaussian model, *ρ*(***x***) ∼ 𝒩 (0, Σ), one can picture the distribution as an ellipsoid defined by the principal axes of the covariance matrix Σ = *J*^−1^. Specifically, since Σ is symmetric, we have the standard mode decomposition

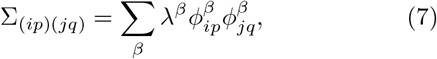

where the *ϕ*^*β*^ are eigenvectors of Σ with eigenvalues *λ*^*β*^. The *ϕ*^*β*^ corresponding to the largest eigenvalues give the directions of the principal modes of variation. For our sparse model at the 32-cell stage, over 80% of the variation is captured in the first three principal axis or modes of variation, shown in Fig. 7C. These modes represent coordinated embryo-wide variations in gene expression, in stark contrast to the modes found for a model without cell-cell interactions (SM Fig S20). We emphasize that each mode (rows of Fig. 7C) corresponds to a set of zygotic genes, which collectively express in particular spatial patterns across the cells of an embryo. The pattern of a specific gene is some linear combination of the PCs (columns of Fig. 7C). Importantly, since we do not know the exact identity of each cell, only which cell type it belongs to, these modes reflect an inference of the model. Moreover, the modes appear somewhat robust to which interaction terms appear in a model, for instance these first three modes explain 20% of the variation for the model inferred from synthetic data, which has a slightly different set of interaction terms, SM Fig. S22. This is notable as these modes exist in a high dimensional space (*n*_*cells*_ × *n*_*PC*_ = 160 for the 32-cell stage), and hence two random vectors are typically almost orthogonal. Simply applying a random rotation to the original regularized model results in the first three unrotated modes explaining only 0.17% of the rotated model’s variance: if the mode inference was sensitive to the model terms, this would be the typical level of variance explained by the first three modes. No collective modes, of the nature we have identified through a careful statistical study, have been inferred previously.

Intriguingly, the primary collective modes in Fig. 7C approximately correspond to the major morphological axes of the embryo, which are coming into being and being maintained, at these stages of development. The first mode represents a left-right mode of variation, which agrees with biological intuition that left and right sides are relatively uncoupled and hence can vary independently. Further, the second mode appears somewhat animal-vegetal like, which, again, would be expected as FGF signaling between animal and vegetal poles has not occured yet, leaving animal and vegetal cells largely coupled at this stage. These modes represent a testable prediction of the theory. Were expression levels measured without losing cell identity, say with *in situ* hybridization or scRNA-seq with careful dissection of embryos, then one should observe the modes of variation, as identified in Fig. 7C, as the primary source of variation around the mean levels.

## IV. DISCUSSION

Our goal was to study variation in ascidian development at the level of the embryo, both to understand what typical variation looks like as well as to construct generative models of the variation. Starting from high quality transcriptomic data [16], across multiple stages and multiple embryos, we developed a new statistically-robust approach to characterize the WT transcriptional states at successive stages and to accurately classify every cell into one of those states. For zygotic transcripts, we found that embryo-to-embryo variation along the WT time axis, as well as expression noise at the level of single cells, was sufficient to explain variation at the 8- and 16-cell stages. At the 32- and 64-cell stages, we show that there are additional collective degrees of freedom in the data which vary from embryo-to-embryo. To understand the source of the collective variation, we introduced statistical physics models, allowing us to infer both interactions as well as the collective modes of variation from limited experimental observations. Our findings shows the maternal to zygotic transition in action, which begins at the 8-cell stage as zygotic transcription turns on in the embryo, with a handful of genes transcribed at the 16-cell stage. As increasing numbers of zygotic genes are expressed at later stages, and different cells become coupled through inheritance and cell-cell communication, non-trival collective variation in the embryo appears.

A central goal of this work was to identify these spatially organized collective modes of variation at the scale of the whole embryo. By searching for such modes, we reveal natural variation present in an otherwise tightly controlled developmental program, in contrast to strong perturbative approaches typically taken to study development. As such, our work represents the desire to go beyond cell typing and leverage single-cell expression measurements to address a broader scope of biological questions, including analyzing the collective behavior of multiple cells rather than one cell at a time. Further, we propose that, since selection acts on the organism rather than its individual cells, the collective modes we identify are also phenotypic traits that naturally vary and are under selection. We hope that our work could open new avenues to study the evolution of developmental programs.

To perform much of our quantitative analysis, we required a high quality identification of cell type from single cell data, and as such we formulated a statistically robust approach to cell typing which can be leveraged by the community broadly. Central to this was the emphasis on utilizing a posterior distribution for gene expression that permits resampling for downstream statistical tasks. Gene expression data is widely recognized as noisy, yet often the only statistical challenge is seen as estimating the most likely value of expression. Instead, by accepting that these measurements have potentially large uncertainty and including this uncertainty in our analysis, we can strengthen our ability to perform not only cell typing, but improve our estimation of covariances and inference of interactions along with related statistical problems. The sequencing data used here was of high depth, but we hope that future studies can investigate a diverse array of developmental programs with shallower sequencing, using some of the techniques developed here.

To build statistical models and remove noise from the data, instead of using hundreds of zygotic genes, we work with a handful of principal components. The principal components span across many genes, with genes often contributing to multiple components. While the building blocks of the developmental process are undoubtedly genes, by working with principal components we are searching for a level of description beyond the single gene and at the collective level of multiple genes, much as in statistical physics we work with collective variables like density rather than individual molecules. Here the choice was data-driven principal components, although one could coarse grain over biologically motivated collections of genes, say in different regulatory pathways. Recent deep learning and large language model approaches [38], trained on a vast corpus of single cell sequencing amongst other biological data, find a non-linear transcriptome latent space. It would be intriguing to take such a latent space as the input degrees of freedom to then be modeled within a statistical physical approach, which could provide an interpretable model for such data.

Statistical physics models have proved powerful at explaining related biological phenomena such as pattern formation in cell layers [39], with maximum entropy models in particular used to study gene regulatory networks [10], protein dynamics [40] and antibody diversity [41]. However, to the best of our knowledge, statistical physical models and the principle of maximum entropy, have not previously been applied directly to scRNA-seq data in order to study the collective expression of multiple cells. Here, our knowledge of the stereotyped spatial connectivity and mean expression level of cells within an embryo allows us to statistically model the joint probability distribution of gene expression for all the cells in an embryo at a given stage - an embryonic transcriptome - with one sample from the generative model representing one embryo. As statistical physics suggests, interacting systems, which cells in an embryo are, have rich collective behavior and should be modeled as such.

Finally, we remind the reader that our approach does not model temporal dynamics. Each stage of the embryo is treated separately despite the fact that they are connected in time. This is a consequence of the destructive nature of scRNA-seq together with the significant length of time between measurements. Expression profiles between stages are seemingly time-discontinuous and so the data is intrinsically non-dynamical. Including deeper lineage statistics then just sister-cell interactions, or creating a joint distribution of all stages at once could begin to incorporate more dynamical information into a model. Perhaps more dynamical measurements are required to fit a more dynamical model, but a dynamical perspective is worth considering as these collective modes are ultimately a spatio-temporal feature of development.

## METHODS

### Raw scRNA data

All experimental data comes from the single cell RNA sequencing experiments of Ref. [16], available with accession numbers, ArrayExpress: E-MTAB-6506, E-MTAB-6508, E-MTAB-6528, E-MTAB-6530. We only consider cell stages 8 to 64 as there is no zygotic expression before stage 8 [16]. In total there are 8 embryos at the 8-cell stage, 11 embryos at the 16-cell stage, 14 embryos at the 32-cell stage, and 8 embryos at the 64-cell stage. Not all embryos are complete.

### Aligning transcripts

We use the reference transcriptome of Ref. [16], ENA accession number PRJEB3758, which was assembled from scRNA transcripts and validated against a *Ciona robusta* reference. We also compared against the Aniseed genome [28] which has a more complete set of genes but does not have untranslated regions, limiting the reads and hence quality of expression estimate of important genes (SM Sec. II). Aligning was performed using the STAR method [42].

### Differentially expressed genes

Given a particular gene, we test whether it is differentially expressed in two cell types, by collecting expression values for all cells in belonging to the two cell types, and performing the standard Wilcoxon rank-sum test [43]. To find all genes that are differentially expressed across any two cell types, we perform the above test but adjust p-values to account for testing across multiple genes and cell type pairs. We consider genes at significance *p <* 0.01 for adjusted p-value as differentially expressed. Fig. 1C shows the differentially expressed genes at the 16-cell stage.

### Zygotic genes

At the *k*-cell stage, we identify a set of purely zygotic genes. First we take all genes which have fewer than 10 reads in every cell at the 4-cell stage, which identifies non-maternal genes which are not expressed at early stages. Of those genes, if any have more than 100 reads in any cell at the *k*-cell stage, we include them in our set of zygotic genes at that stage. This criteria excludes genes that have maternal transcripts present which are then additionally are transcribed zygotically, as well as lowly expressed zygotic genes. It correctly identifies the major zygotic genes and excludes any maternal factors. Next, we exclude any zygotic genes that are not differentially expressed across cell types. We do this as certain genes are expressed in every cell in some embryos and in no cells in others. While interesting, these genes are not involved in the self-organizational program, and so we exclude them from the analysis.

### Maternal genes

To select a set of purely maternal genes, we take all genes with more than 500 reads in any cell at the 4-cell stage that are also not differentially expressed in any subsequent stage. This gives us 2268 genes that are purely maternal factors and not zygotically transcribed.

### Determining cell-cell contact network

From light-sheet microscopy images of embryos segmented with the ASTEC framework [15], two cells were determined to be in contact if, on average, the area of contact represented at least 5% of one of the cells surface area. This average was taken over the left and right side of the embryos, assuming a symmetric contact map, as well as across two embryos.

## ACKNOWLEDGEMENTS

We thank Kilian Biasuz for providing data and assistance in determine cell-cell contacts, and Christelle Dantec for discussions about aligning transcripts. D.J.S. thanks Eric Siggia and Dillon Cislo for helpful discussions and valuable comments on the manuscript. M.M. and D.J.S. were supported by The National Science Foundation-Simons Center for Quantitative Biology at Northwestern University and the Simons Foundation grant 597491. M.M. is a Simons Investigator. This project has been made possible in part by grant number DAF2023-329587 from the Chan Zuckerberg Initiative DAF, an advised fund of the Silicon Valley Community Foundation. This work was supported by NICHD 1R01HD088831-01 to M.M., a binational “NSF-ANR: Collaborative Research: A mechanical atlas for embryogenesis at single-cell resolution.” (National Science foundation [2204237] and ANR ANR-21-CE13-0046) to M.M. and P.L.. This research was supported in part by grants NSF PHY-1748958 and the Gordon and Betty Moore Foundation Grant No. 2919.02 to the Kavli Institute for Theoretical Physics (M.M.).

