## Supplementary Material for "Physical modeling of embryonic transcriptomes identifies collective modes of gene expression"

### CONTENTS

I. Introduction 2

II. Preprocessing of sequencing data 3
- A. Alignment 3
- B. Normalization 3

III. Recovering cell types 4
- A. Identifying important genes 5
- B. Core clustering algorithm 5
  - 1. Initialization 7
  - 2. Modified K-means 7
  - 3. Merging clusters 7
- C. Cluster consistency 8
  - 1. Testing cluster significance 10
- D. Overall clustering algorithm 10
- E. Results 11
  - 1. 64-cell stage, caption for Fig. S10 15
- F. Overall clustering consistency 19
- G. Comparison with standard pipeline 19

IV. Analysis of data post clustering 21
- A. Singular values 21
- B. Null model for singular values 23
- C. Covariances 23
- D. Shuffle tests 24

V. Maximum entropy framework 25
- A. Maximum entropy for embryos 27
- B. No cell-cell interactions 28
- C. Full generality 29
- D. Initial guess 30

VI. Inference 30
- A. Inference results 32

VII. Synthetic data and inference testing 32
- A. Synthetic data 32
- B. Robustness of inferred modes 34
- C. Gaussian assumption 35

A. Maximum entropy Gradients 37

References 38

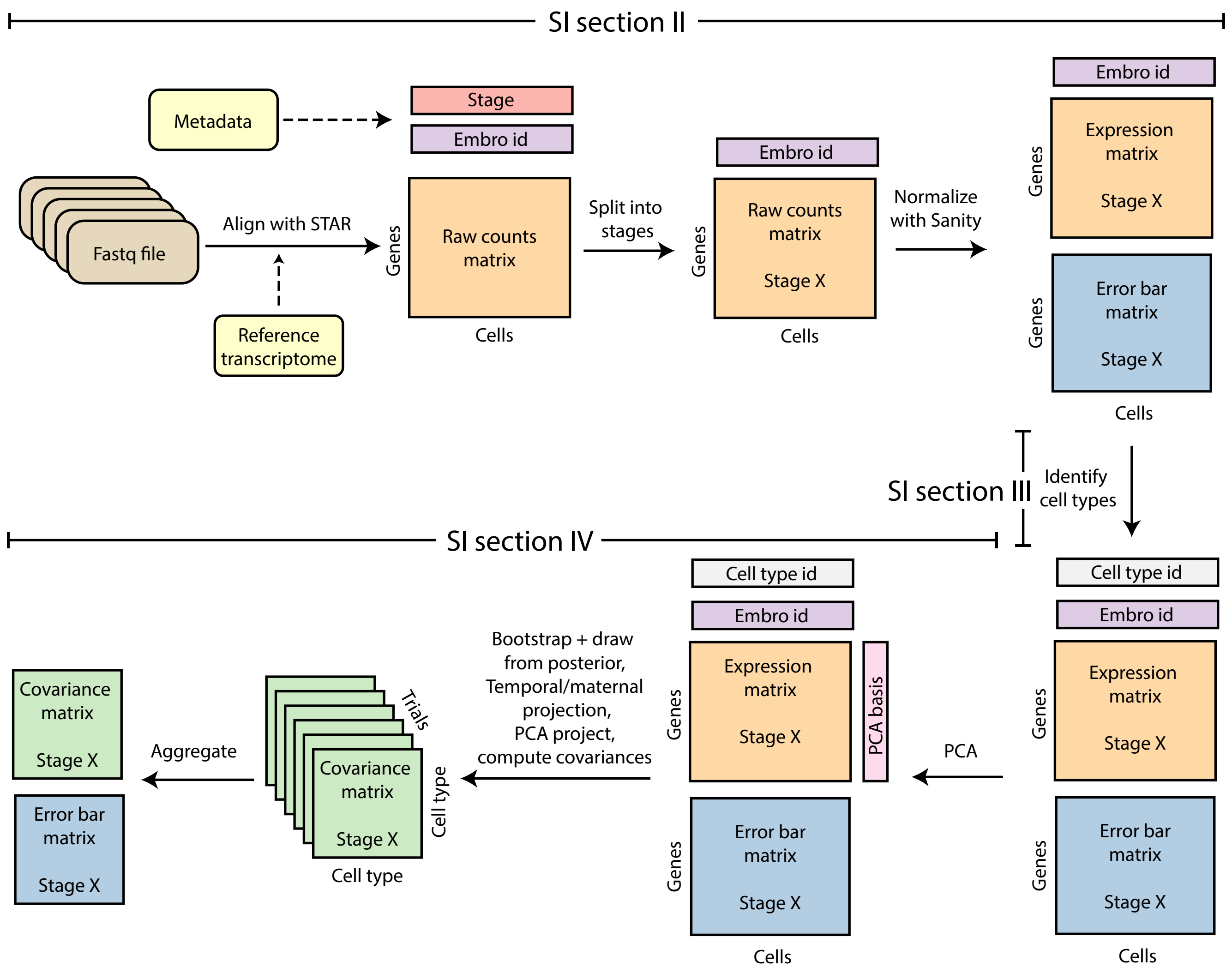


FIG. S1. Summary of overall workflow for experimental data, starting with experimental fastq files (raw sequencing output) and a reference transcriptome, ending with covariance matrices which are the input data to statistical models. The initial preprocessing, including alignement and normalization is discussed in Sec. II. The recovery of cell types is discussed in Sec. III. The temporal projection, PCA analysis, and computation of covariances is discussed in Sec. IV.

### I. INTRODUCTION

This supplementary material contains additional details, including complete numerical and algorithmic procedures, which were outlined in the main text. In Section II we describe the preprocessing of the sequencing data, specifically alignment and normalization. In Section III, we describe how the cell types are recovered. In Section IV, we discuss the temporal projection, PCA analysis, shuffling tests, and covariance calculations. The overall workflow of data processing, from raw data files to the computed covariance matrices is summarized in Fig. S1. In Section V we discuss the maximum entropy framework and mathematical modeling, in Section VI we discuss the inference procedure, and in Section VII we discuss how we validate the inference procedure with synthetic data.

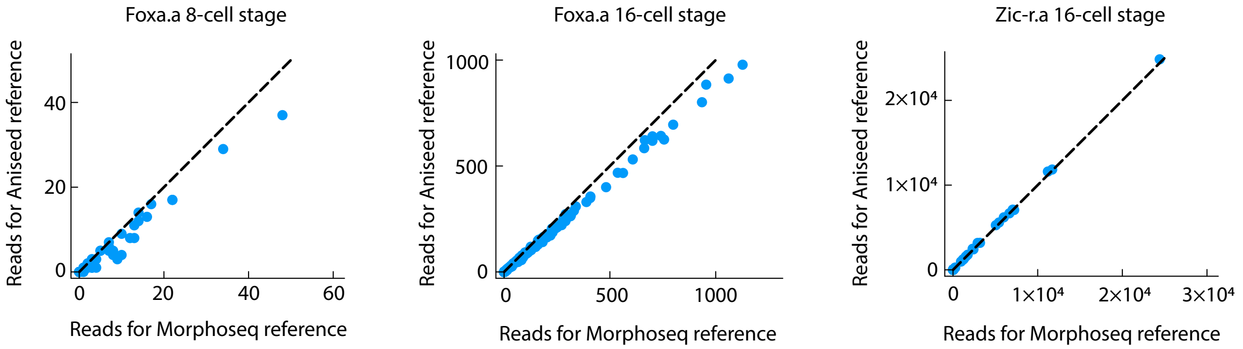


FIG. S2. Comparison of Aniseed and Morphoseq references aligned with STAR. Some genes like *Foxa.a*, (left and middle), have significant untranslated regions. The Morphoseq reference contains these regions and hence has higher numbers of aligned reads than the Aniseed reference which does not contain these regions. For other genes, like *Zic-r.a*, the two perform equivalently (right).

### II. PREPROCESSING OF SEQUENCING DATA

#### A. Alignment

In order to estimate gene expression, raw transcripts must be aligned with a reference genome or transcriptome and then normalized. Multiple computational methods for aligning and normalizing exist, and two references for *P. mammillata* exist. Here, we will briefly compare algorithms and references, and justify the final choices we used throughout.

In Ref. [18], a transcriptome was assembled from raw transcripts and validated against a *Ciona robusta* reference, which we will refer to as the Morphoseq reference. Alternatively the Aniseed [3], database for tunicates contains a reference genome for *P. mammillata*. The Aniseed reference contains a more complete list of genes (19,467) than the Morphoseq reference (9656), and a larger number of transcripts can be aligned to the Aniseed reference than to the Morphoseq reference, see Table S1. However, the Morphoseq transcriptome contains untranslated regions (UTRs) which Aniseed does not. Therefore, for many important genes, more transcripts can be found when using Morphoseq then when using Aniseed, Fig. S2. For this reason, and because the Morphoseq transcriptome by construction contains genes which are expressed during these early cell stages, we use the Morphoseq reference.

To align transcripts to a reference, we tried Bowtie [7], used in Ref. [18], as well as STAR [4]. Both are standard methods. Overall, STAR found more alignments than Bowtie, and the additional alignments do not appear to be spurious upon manually inspection. For this reason, we will use the STAR aligning method throughout.

TABLE S1: Comparison of aligning choices

| Parameter combinations | Total reads | Non-zero genes | Reads per cell (8-cell stage) | Reads per gene (8-cell stage) | Reads per cell (64-cell stage) | Reads per gene (64-cell stage) |
|---|---|---|---|---|---|---|
| STAR align, Aniseed reference | $4.27 \times 10^9$ | 19178 | $5.45 \times 10^6$ | 328 | $3.06 \times 10^6$ | 162 |
| STAR align, Morphoseq reference | $3.96 \times 10^9$ | 9656 | $4.92 \times 10^6$ | 515 | $2.85 \times 10^6$ | 295 |
| Bowtie align, Morphoseq reference | $3.24 \times 10^9$ | 9656 | $3.98 \times 10^6$ | 418 | $2.32 \times 10^6$ | 241 |

#### B. Normalization

Having chosen the Morphoseq reference and STAR aligning tool, we now have a cell by gene matrix of reads for each cell stage. This matrix has to be further processed, or normalized, in order to do statistics with it. Typically, such normalization procedures have two major components [1]. Firstly, rather than work with absolute counts, $X_{cg}$, take the relative fraction of counts, $X_{cg}/\sum_i X_{ci}$. This relative fraction means cells of different sizes but the same mRNA composition are considered to have the same expression, and also accounts for the difference in mRNA capture

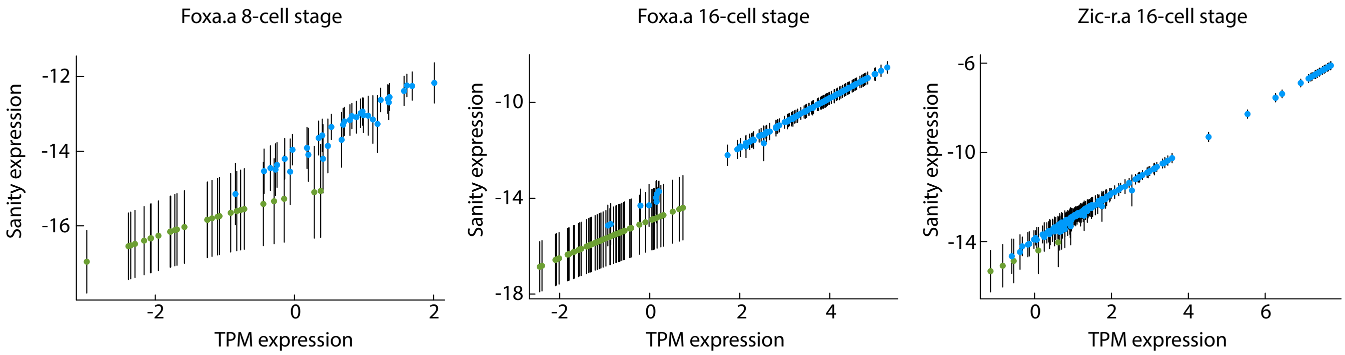


FIG. S3. Comparison of Sanity and TPM normalization methods. Error bars are ±1 standard deviation. Each point is one cell, points are green when zero transcripts were present. When expression is high, both methods find similar results. For low expression Sanity generally identifies cells with zero reads as having lower expression than cells with non-zero reads, but more importantly recognizes the uncertainty in expression when actual numbers are small. Note that we will only ever compare variation in expression around the experiment wide average, the average value (e.g., around -14 in Sanity, 0 in TPM) is not important.

rate during sequencing which can vary from cell to cell. Secondly, take the log of this relative fraction, as the fold change, not the change in absoulute counts, is important. Going from 2 to 200 mRNA counts for a particular gene is a meaningful change, but 2002 to 2200 is likely not. Effectively, this performs a variance stabilization [1], an important statistical step for comparing variation across different variables. The simplest way to combine these two steps is in the transcripts per million reads (TPM) method,

$$Y_{cg} = \log\left[\frac{X_{cg}+1}{\sum_i X_{ci}} \times 10^6\right], \tag{S1}$$

where the one is taken to avoid log of zero when there are no reads for a particular gene. Variations on this include accounting for gene length, and estimating a normalizing factor using a small number of consistently expressed genes, rather than summing over all genes [13].

Sanity normalization attempts to infer the log fraction of total mRNA molecules corresponding to a particular gene, $\alpha_{cg}/\sum_d \alpha_{dg}$, where $\alpha_{cg}$ is the expected number of mRNA molecules for a particular cell and gene. By constructing a probability distribution for raw counts $X$ given a particular state of the cell, $\mathbb{P}(X|\alpha)$, Bayesian inference allows for the computation of a posterior distribution of the parameters given data, $\mathbb{P}(\alpha|X)$. The posterior of Sanity estimates an most likely expression level $Y_{cg}$, together with a independent Gaussian, $\mathcal{N}(Y_{cg}, \epsilon_{cg})$, where the Gaussian effectively gives an error-bar to the expression. Gene-wide uncertainty is not considered here as the overall level of a gene is never directly used, only the variation between different cells and embryos.

Either Sanity or TPM performs similarly at estimating expression, Fig. S3, indeed, as recent work indicates [1]. For a particular cell stage, cells from all embryos at that stage were normalized together. However, we again stress that the main reason we use Sanity is to ensure that we have an estimate of the uncertainty about any measurement.

### III. RECOVERING CELL TYPES

From the normalized transcriptomic data, a major technical challenge is to recover the cell type of each cell and thus gain information about it's position in the embryo. To do so effectively, we devise a clustering scheme based on the following principles:

1. While many genes are expressed, only a handful of genes are differentially expressed across the embryo and thus informative about cell types. These genes alone should be used to cluster the data. Genes that are not differentially expressed add unwanted noise, which obfuscates genes with useful signal and makes recovering cell types more difficult.

2. We know the structure of an embryo, and we use that information. Specifically, every embryo has at most two cells for each cell type, assuming left-right symmetry in mean expression (an assertion we will test later) and accounting for the fact that some cells were lost during sequencing. A clustering that assigns 3 cells in the same embryo to a cluster, and no cells in any other embryo must be incorrect.

3. There is uncertainty in the data, both in the expression levels and also from the finite number of experiments. We ensure the clustering is robust, and to assess our confidence in the clustering based on this uncertainty.

4. We should proceed hierarchically and not all informative genes need to be used at once. For instance, the expression of *Pen1* gene can distinguish B4.1 germ cells from somatic cells at the 8-cell stage. Yet its expression is unhelpful for distinguishing the remaining A4.1, a4.2, and b4.2 cells. Instead, after using of *Pen1* to identify B4.1 cells, we should use different genes to distinguish the remaining cells.

Below, we outline the technical specifics which employ these principles. We describe the method to select informative genes, the core clustering algorithm, and the cluster consistency score, before describing the overall algorithm which utilizes these component parts.

### A. Identifying important genes

Consider a data set which can be separated into some number of clusters by typical clustering algorithms. If one adds enough additional dimensions in which the data values are randomly distributed, and thus contain no information about the clusters, these same algorithms will fail to recover the clusters [11]. In developmental systems, there are a large number uninformative genes, that may be crucial to cell survival, yet are not differentially expressed across cell types. Including these uninformative genes may result in the principal components of all expression levels essentially fitting to noise and containing no information about cell types. Instead, we look for the small number of informative genes from which cell types can be identified.

If we knew the cell types ahead of time we could easily identify the genes that are differentially expressed across different cell types. The challenge is that the cell types or clusters are exactly the information we are trying to learn. However, what Ref. [11] show is that by averaging over many generated trial clusters, the small number of informative genes can be recovered. Their method first splits the data into some number $K$ of trial clusters. Given two of these clusters they find a sparse linear separation between them, making a note of which dimension contribute most to that linear separation. This is repeated for all pairs of clusters, and the whole procedure is repeated for many possible clusterings. The dimensions which most often contribute to linear separations between clusters are the informative dimensions.

To adapt this method to the expression data, we note that the trial clusters should be consistent with the known structure of the true clusters. While we do not know the true clusters or cell types *a priori*, we do know that a cluster representing a cell type will have, say, 2 cells in every embryo rather than 4 cells in one embryo and 1 in another. With this knowledge, we generate a trial cluster by taking exactly two cells from each embryo. It does not matter whether the actual cell type contains 2 or 4 cells per embryo (one expects an even number due to left-right symmetry), as this method only seeks to identify meaningful dimensions before cluster sizes are determined.

Running this procedure tells us, for each gene, how many times it was the dimension which contributed most to a linear separation. Effectively, this gives us a importance ranking of genes and allows us to select the, say, 10 most important genes. The exact steps of the algorithm are outlined in Algorithm 1.

**Algorithm 1** Gene selection algorithm, modified from Ref. [11]

**Input:** Expression levels $X_{ij}$ with uncertainty $\epsilon_{ij}$, cells per embryo $S$, $N_{trials} = 50,000$, gene ranking vector $\vec{g} = 0$, $\lambda = 0.05$
**for** $1 : N_{trials}$ **do**
  $Y_{ij} \leftarrow$ draw from posterior $X_{ij} + \mathcal{N}(0, \epsilon_{ij})$
  $C \leftarrow$ Randomly assign $Y$ into $S/2$ clusters each with at most 2 cells per embryo
  **for** each pair $l$, $m$, of clusters **do**
    $\vec{\theta} \leftarrow \arg\min_\theta \sum_{p\in C^l} [1 - \vec{Y}_p \cdot \vec{\theta}]_+ + \sum_{p\in C^m} [1 + \vec{Y}_p \cdot \vec{\theta}]_+ + \lambda||\vec{\theta}||_1$, with $\vec{Y}_p = \{Y_{pj}\}$
    $j^* \leftarrow \arg\max_j \theta_j$
    $g_{j^*} \leftarrow g_{j^*} + 1$
  **end for**
**end for**
  **return** $\vec{g}$

### B. Core clustering algorithm

Suppose we have chosen a small number of genes, and have some cells, with known embryo origin, that we wish to sort into $K$ clusters. Later, we will discuss how we select the genes, the number of clusters, as well as how we apply this clustering hierarchically. We take a standard clustering algorithm, K-means, and adapt it to our specific system.

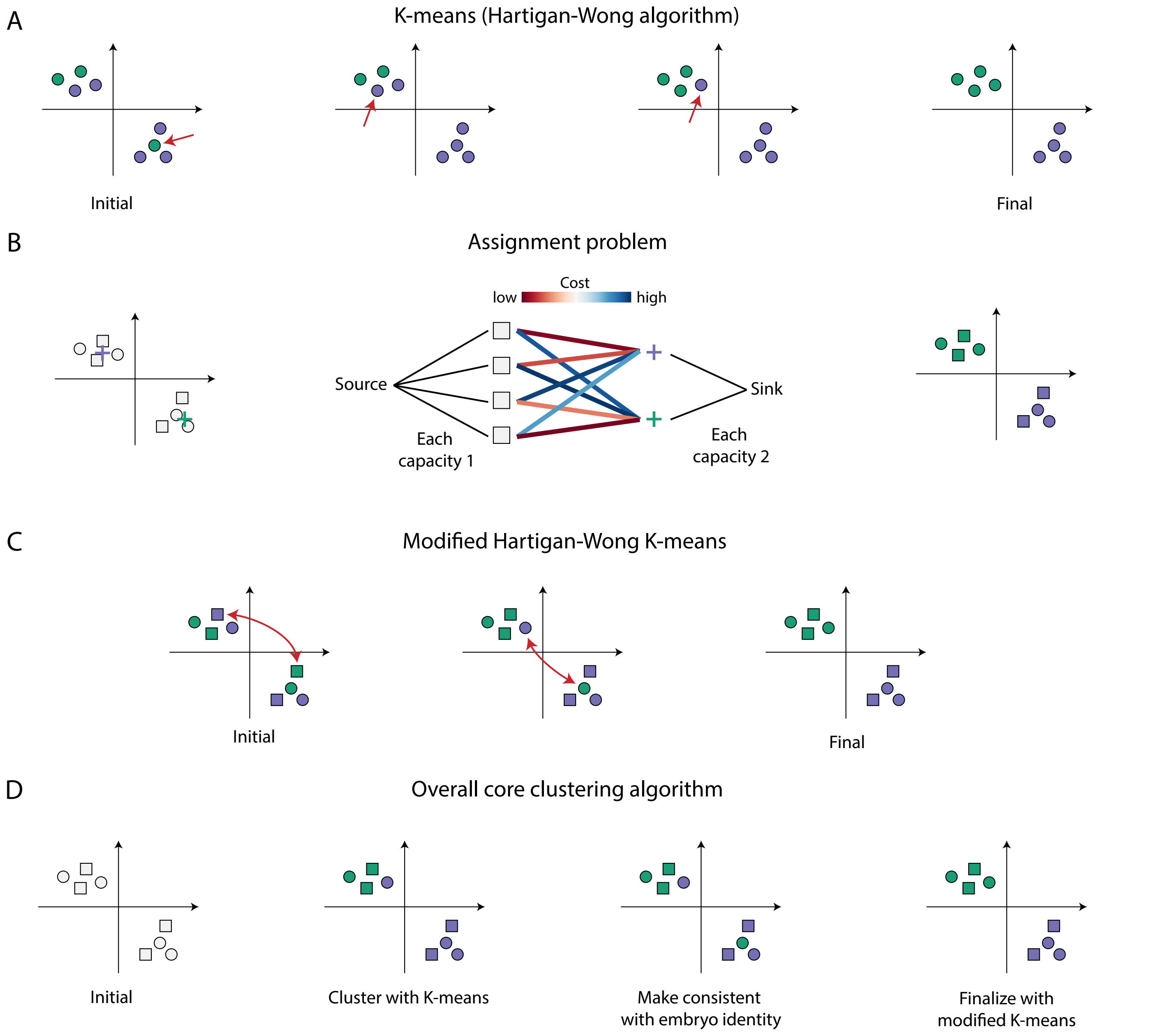


FIG. S4. Summary of overall workflow for the core clustering algorithm. (A) K-means clustering using the Hartigan-Wong algorithm. Starting from a random assignment, we find the best point (red arrow) to reassign clusters in order to minimize the K-means score (cluster moment of inertia). The algorithm terminates when any reassignment would increase the score. (B) To find a cluster consistent with embryo identity constraints, we solve a transport problem. For each embryo, we try to assign cells to a nearby cluster center (colored + signs, left) subject to the constraint that no cluster can have more than $c_k$ cells per embryo (in figure $c_k = 2$), resulting in a transport problem (center). After solving this for each embryo, we have a feasible clustering (right). (C) To be consistent with embryo constraints, we modify the Hartigan-Wong algorithm so that it can only swap cell's cluster identity within the same embryo. (D) Overall, we find an initial clustering with K-means, solve a transport problem to find a clustering consistent with the embryo constraints, and then iterate this clustering with a modified K-means algorithm.

In particular, we assume that cluster $k$ contains at most $2c_k$ cells from each embryo, so that each cluster is spread across all embryos, with $\sum_k 2c_k$ equalling the total number of cells. Note that for the experimental data, certain cells can be missing from a particular embryo due to not making quality control [18], and so a cluster of the experimental data may have 2 cells in embryos 1, 2, 3, and 1 cell in embryo 4 (say). The algorithm is summarized in Algorithm 2.

#### 1. Initialization

The modified K-means, which we will describe in detail later, is slow without a good initialization. A standard clustering algorithm can run rapidly and produce good initial guesses for clusters, but these clusters will not conform to the known structure of the embryo. However, we can solve a transport problem that takes the cluster guesses and transforms them into a clustering that obeys the known structure and can serve as the initial clustering to run the modified K-means on.

At the $2s$-cell stage of the embryo, we initialize with $s$ cell types. If we seek fewer clusters, $K < s$, we merge resulting clusters, as described later. We start by clustering of the data into $s$ clusters using K-means (although any other algorithm could be employed here), getting a clustering $\mathcal{C}_0$. This cannot serve as a plausible clustering as it may not obey the constraint that at most 2 cells from each embryo are in a cluster, but we will transform it into a plausible cluster. We do this by considering each embryo one at a time, and finding the best way to divide this embryo between the $s$ existing clusters of $\mathcal{C}_0$, whilst obeying the constraints.

First, compute the mean value of each cluster of $\mathcal{C}_0$, namely $\bar{X}_1, \ldots, \bar{X}_s$. We wish to divide the expression levels, $\{\vec{x}_c\}$ for one particular embryo amongst the $s$ clusters, where $c$ indexes the (at most) $2s$ cells. This may not be as simple as assigning each cell to it's existing cluster, or the cluster with the nearest mean value, as this may put more than two cells in a cluster. Instead, we consider the cost of putting cell $c$ in cluster $l$ as $T_{cl} = ||\vec{x}_c - \bar{X}_l||_2$. We then have a formal transport problem where sources, $c$, of strength 1 are connected to sinks $l$ of strength 2, and the cost of transporting $c$ to $l$ is $T_{cl}$, Fig. S4B. We can solve this linear program with standard algorithms, and the sparse solution determines which cells should go into which clusters whilst obeying the constraints. Performing this for every embryo gives us an initial clustering $\mathcal{C}_1$ that we can pass into the modified K-means algorithm.

#### 2. Modified K-means

Starting from the initialized clustering, $\mathcal{C}_1$, satisfying our biological constraint on cluster membership, we proceed with a modified Hartigan–Wong method [6], where the unmodified Hartigan-Wong is an algorithm for implementing a K-means clustering, Fig. S4A.

In short, for a cluster $S_j$, we define the function $\varphi(S_j)$ as

$$\varphi(S_j) = \sum_{\boldsymbol{x} \in S_j} (\boldsymbol{x} - \boldsymbol{\mu}_j)^2, \tag{S2}$$

where $\boldsymbol{\mu}_j$ is the mean taken over all $\boldsymbol{x} \in S_j$. Then for all potential swaps, where cells $\boldsymbol{x}$ and $\boldsymbol{y}$ are in the same embryo but in different clusters $S_n$ and $S_m$, and so could be interchanged, we define a quantity

$$\Delta(\boldsymbol{x}, \boldsymbol{y}, n, m) = \varphi(S_n) + \varphi(S_m) - \varphi((S_n \backslash \{\boldsymbol{x}\}) \cup \{\boldsymbol{y}\}) - \varphi((S_m \backslash \{\boldsymbol{y}\}) \cup \{\boldsymbol{x}\}), \tag{S3}$$

and also

$$\Delta(\boldsymbol{x}, n, m) = \varphi(S_n) + \varphi(S_m) - \varphi(S_n \backslash \{\boldsymbol{x}\}) - \varphi(S_m \cup \{\boldsymbol{x}\}), \tag{S4}$$

representing the case when $S_m$ has fewer cells than $2c_m$ in a particular embryo than $S_n$ and so $\boldsymbol{x} \in S_n$ can be moved without swapping whilst not violating our biological constraint on cluster membership.

For all feasible swaps or moves, we can compute the quantity $\Delta$, and implement the swap with the largest value of $\Delta$. This continues until all values of $\Delta$ are negative, at which point the algorithm has converged, and we are left with a clustering $C_2$, Fig. S4C.

#### 3. Merging clusters

So far, $C_2$ is a clustering with $s$ clusters at the $2s$-cell stage. Suppose instead that we want $K < s$ clusters, optionally with known sizes $c_1, \ldots, c_K$. We proceed as follows. First we take two clusters that are closest, as measured by the Ward distance and merge them, resulting in a clustering with $s - 1$ clusters. Next, we rerun the modified K-means algorithm, since this new clustering may not be a local minimum of the modified K-means. We repeat this overall procedure iteratively until we reach $K$ clusters.

If we have target sizes for the clusters, $c_1, \ldots, c_K$, then the above merging procedure is not guaranteed to result in the required cluster sizes. In this case, we introduce an additional criteria to the merging. Given two clusters that are closest in Ward distance, they are only merged if the resulting merger either gives the target cluster sizes, or could plausibly reach the target cluster sizes after further merges. If they cannot reach the target sizes, then try merging the second closest pair of clusters and so on until one that can be merged is found. Computing whether a merge is feasible can be solved efficiently with dynamical programming, see Algorithm 3.

**Algorithm 2** Core clustering algorithm

**Input:** Data $X$, embryo identity $\vec{e}$, cells per embryo $S$, number of clusters $K \leq S/2$, (optional cluster sizes $c_k$ for $k = 1 : K$).
$\mathcal{C} \leftarrow$ K-means clustering of $X$ into $S/2$ clusters.
$\vec{x}_k \leftarrow$ cluster centers from $\mathcal{C}$
$\mathcal{C} \leftarrow$ assign $X$ to clusters with no more than 2 cells per cluster using $\vec{x}_k$ and linear programming.
$\mathcal{C} \leftarrow$ run modified K-means with $\mathcal{C}$ as input.
**while** #clusters($\mathcal{C}$) $> K$ **do**
  $(p, q) \leftarrow \arg\min_{p \neq q} d(\mathcal{C}_p, \mathcal{C}_q)$, where $d$ is the Ward distance between clusters. If $c_k$ specified, only minimize over $p$, $q$ that can be merged consistently with $c_k$.
  $\mathcal{C} \leftarrow$ merge clusters $p$ and $q$.
  $\mathcal{C} \leftarrow$ run modified K-means with $\mathcal{C}$ as input.
**end while**
  **return** $\mathcal{C}$

**Algorithm 3** Feasibility check to see whether target cluster sizes can be achieved from merging current cluster sizes

**Input:** Proposed cluster sizes $\vec{n} = (n_1, \ldots, n_R)$, target cluster sizes, $\vec{n}_t = (c_1, \ldots, c_K)$.
$\vec{n} \leftarrow sort(\vec{n})$
$\vec{n}_t \leftarrow sort(\vec{n}_t)$
$d \leftarrow map(\vec{n}_t \rightarrow 1)$
**function** FEASCHECK($\vec{m}$)
  **if** $\vec{m}$ is in $d$ **then return** $d[\vec{m}]$
  **end if**
  **if** $length(\vec{m}) = K$ **then return** 0
  **end if**
  **for** $1 \leq p < q \leq length(\vec{m})$ **do**
    $\vec{r} \leftarrow sort(\{m_1, \ldots\} \backslash \{m_p, m_q\} \cup \{m_p + m_q\})$
    $f \leftarrow$ FEASCHECK($\vec{r}$)
    **if** $f = 1$ **then return** $d[\vec{m}] \leftarrow 1$
    **end if**
  **end for**
  **return** $d[\vec{m}] \leftarrow 0$
**end function**
  **return** FEASCHECK($\vec{n}$)

#### C. Cluster consistency

For any set of genes and any cluster sizes, our algorithm will produce a clustering. Therefore, we need a way to determine from the data how much to trust this clustering, which in turn can inform us how to choose our cluster sizes and input genes. We will do this by computing a consensus cluster score [12], using the fact that we have an estimate for the uncertainty of our expression values. In short, we resample the data from the posterior, then subsample by a factor of 0.9 (probability of a cell being kept is 0.9), and then cluster. We repeat this $n$ times. Two cells, $p$, $q$ then have a score $M_{pq} = S_{pq}/I_{pq}$, where $S_{pq}$ counts the number of times cells $p$ and $q$ were in the same cluster, and $I_{pq}$ counts the number of times they were in the same subsampled data set. To compare two clusterings, with different numbers of clusters or different input genes, we use the proportion of ambiguous clustering (PAC) score of $M$ [17]. In short, if the clustering was perfectly consistent, $M_{ij}$ would take values of 0 or 1, if inconsistent it would take intermediate values. The PAC score is defined as $\text{PAC}(M) = P_{90}(M) - P_{10}(M)$, where $P_i(M)$ is the $i^{th}$ percentile of all values in the set $\{M_{pq}, p < q\}$, and a lower PAC score represents a more consistent clustering. Starting with the full data set, we run the gene ranking algorithm and take the top $g_{max}$ genes. We next choose a cluster size $K$. For this choice of $g_{max}$ and $K$ we calculate the PAC score using $n = 500$ trials, each time sampling data from the posterior of expression and subsampling by a factor of 0.9. This score is a function of the cluster number $K$ and the set of $g_{max}$ genes $\{g\}$, $\text{PAC} = \text{PAC}(K, \{g\})$. Consider now removing a gene $g_i$ from this set, and recomputing the

PAC score. If

$$\min_i \ \mathrm{PAC}(K, \{g\} \backslash g_i) < \left(1 + \frac{2}{\sqrt{n}}\right) \mathrm{PAC}(K, \{g\}), \tag{S5}$$

then we reduce the gene set to $\{g\} \backslash g_j$ where $j$ is the argmin and iterate. The $\sqrt{n}$ term is to promote sparsity where the scores are approximately equivalent up to finite sampling effects.

After iterating some number of times, suppose we have some reduced set $\{g\}$ and have computed $\mathrm{PAC}(K, \{g\} \backslash g_i)$ for all $g_i \in \{g\}$, but no single gene reduction improves the score. We then compute $\mathrm{PAC}(K, \{g\} \backslash \{g_1, g_2\})$, where $g_1$ and $g_2$ are the genes that have the smallest and second smallest PAC scores after removal. If this results in an improvement we remove these genes and begin iterating again. If it does not, we try removing the 3 genes with smallest PAC scores and so on. This is necessary as sometimes removing any one gene increases the PAC score, but removing 3 or 4 genes massively decreases it. Ideally we would try all subsets of genes but this is not computationally feasible, so we employ the iterative approach above which works well in practice.

When this iterative process is finished and no more genes can be removed without increasing the PAC score, we are left with the best PAC score possible for $K$ clusters, or $\mathrm{PAC}(K)$. We then compare the scores for $K = 2, \ldots K_{max}$, and selecting the $K^*$ with minimal score. Overall, the above procedure for selecting the best genes and cluster size is summarized in Algorithm 4.

**Algorithm 4** Find the most robust set of genes and clusters for a given set of data.

**Input:** $X$,$\epsilon$, $K_{max}$, $\{g\}$, $n = 500$

**function** PAC($\{g\}$,K)
  **for** $s = 1 : n$ **do**
    $Y_{ij} \leftarrow$ draw from posterior $X_{ij} + \mathcal{N}(0, \epsilon_{ij})$
    $Y \leftarrow$ subsample $Y$ with factor $p_Y = 0.9$
    $\mathcal{C}_s \leftarrow$ cluster $Y$ into $K$ clusters (Algorithm 2)
  **end for**
  $S_{pq} \leftarrow$ number of times cells $p$ and $q$ are in same cluster $\{\mathcal{C}_s\}$
  $I_{pq} \leftarrow$ number of times cells $p$ and $q$ are in sample
  $M = \{S_{pq}/I_{pq}, p < q\}$
  **return** (90% percentile of $M$) – (10% percentile of $M$)
**end function**

**function** BestGenes($\{g\}$,K)
  $\alpha \leftarrow \mathrm{PAC}(\{g\}, K)$
  **for** $g_j$ in $\{g\}$ **do**
    $\alpha_j \leftarrow \mathrm{PAC}(\{g\} \backslash \{g_j\}, K)$
    **if** $\alpha_j < (1 + 2/\sqrt{n})\alpha$ **then**
      **return** BestGenes($\{g\} \backslash \{g_j\}, K$)
    **end if**
  **end for**
  $\{g\} \leftarrow$ sort $\{g\}$ by $\alpha_j$ values
  **for** sorted $g_j$ in $\{g\}$ **do**
    $\beta \leftarrow \mathrm{PAC}(\{g\} \backslash \{g_1, \ldots, g_j\}, K)$
    **if** $\beta < (1 + 2/\sqrt{n})\alpha$ **then**
      **return** BestGenes($\{g\} \backslash \{g_1, \ldots, g_j\}, K$)
    **end if**
  **end for**
  **return** {g}
**end function**

**for** $K = 2 : K_{max}$ **do**
  $\{\tilde{g}\}_K \leftarrow$ BestGenes($\{g\}, K$)
  $\alpha_K \leftarrow \mathrm{PAC}(\{\tilde{g}\}_K, K)$
**end for**
**return** $\{\tilde{g}\}_{K^*}, K^*$ for $K^* = \arg\min_K \alpha_K$

#### 1. Testing cluster significance

Our clustering algorithm, (Algorithm 2) will always find $K$ clusters, whether the data supports it or not, and whatever the PAC score may be. One could set a tolerance for the PAC score; if the PAC score is too high the clustering is rejected. Instead of this somewhat arbitrary threshold, we wanted to check whether the PAC score of a clustering was significant relative to a null clustering, and reject or accept the clustering based on that. To obtain a null, we take the cells and shuffle the embryo labels. If there was truly genes differentially expressed within embryos then this shuffling should make the cluster consistency worse, if there are no such genes then the shuffling will not make a difference. We do this shuffling $m = 200$ times, and keep the round of clustering if it is in the $5^{th}$ percentile or smaller of PAC scores. It may well be the case that the clustering is statistically significant, but has a fairly high PAC score indicating uncertainty in the clustering.

**Algorithm 5** Test whether a clustering is significant. Note that the PAC function from Algorithm 4 implicitly depends on embryo identity from the clustering step. We make the dependence explicit here.

**Input:** $X$, $\epsilon$, $K$, $\{g\}$, embryo identity $\vec{e}$, $m = 200$,
$\alpha \leftarrow \text{PAC}(\{g\}, K, \vec{e})$
**for** $s$ in $1 : m$ **do**
  $\vec{e} \leftarrow \text{permute}(\vec{e})$
  $\alpha_s \leftarrow \text{PAC}(\{g\}, K, \vec{e})$
**end for**
**if** $\alpha < 5\%$ percentile of $\{\alpha_g\}$ **then**
  **return** Cluster significant
**end if**
**return** Cluster not significant

### D. Overall clustering algorithm

As outlined, we have

- A way to rank informative genes, Algorithm 1.
- A way to cluster data into $K$ clusters, Algorithm 2.
- A way to find a set of genes and number of clusters that is maximally robust, Algorithm 4.
- A way to decide whether to cluster or not, Algorithm 5.

We combine these ingredients and work hierarchically to obtain an overall clustering.

Starting with the full data set, we can use Algorithm 1 to select the $g_{max}$ most informative genes, and then Algorithm 4 to find the number of clusters, $K^*$, to keep and a more restrictive gene set $\{g^*\}$ to use for clustering. To actually compute the final cluster assignment, we perform the clustering on data drawn from the posterior a further $n$ times, selecting the most typical clustering to keep as the final clustering. The similarity between two clusters is determined through counting how often pairs of cells $p$, $q$ are either in the same cluster or in different clusters across both clusterings. The cluster with the highest average similarity score is kept.

We have introduced parameters, which we take as $g_{max} = 15$, $K_{max} = 5$, $n = 500$. Ideally all of these parameters would be taken to be as large as possible, while remaining numerically feasible. With this choice, the total number of genes used is always less than $g_{max}$, the number of clusters is less than $K_{max}$, suggesting that increasing them would not make a difference, and the selection does not appear to be sensitive to $n$ either.

We now have a way to cluster the data in the most robust way possible. However, this is not the final clustering. Indeed, at the 8-cell stage, it can be robustly determined whether cells are B4.1 or not, and therefore the most robust clustering is into 2 clusters; one of 2 cells and one of 6 cells. However, it is possible to take the remaining cluster, of A4.1, a4.1, b4.1 cells and further cluster these. The overall clustering then proceeds hierarchically, taking each cluster and splitting it into smaller clusters, following the procedure outlined above. If a cluster cannot be divided into further clusters (Algorithm 5), then the algorithm moves on to a different cluster. If no more clusters can be divided into further clusters then the algorithm stops. This hierarchical procedure is outline in Fig. S5.

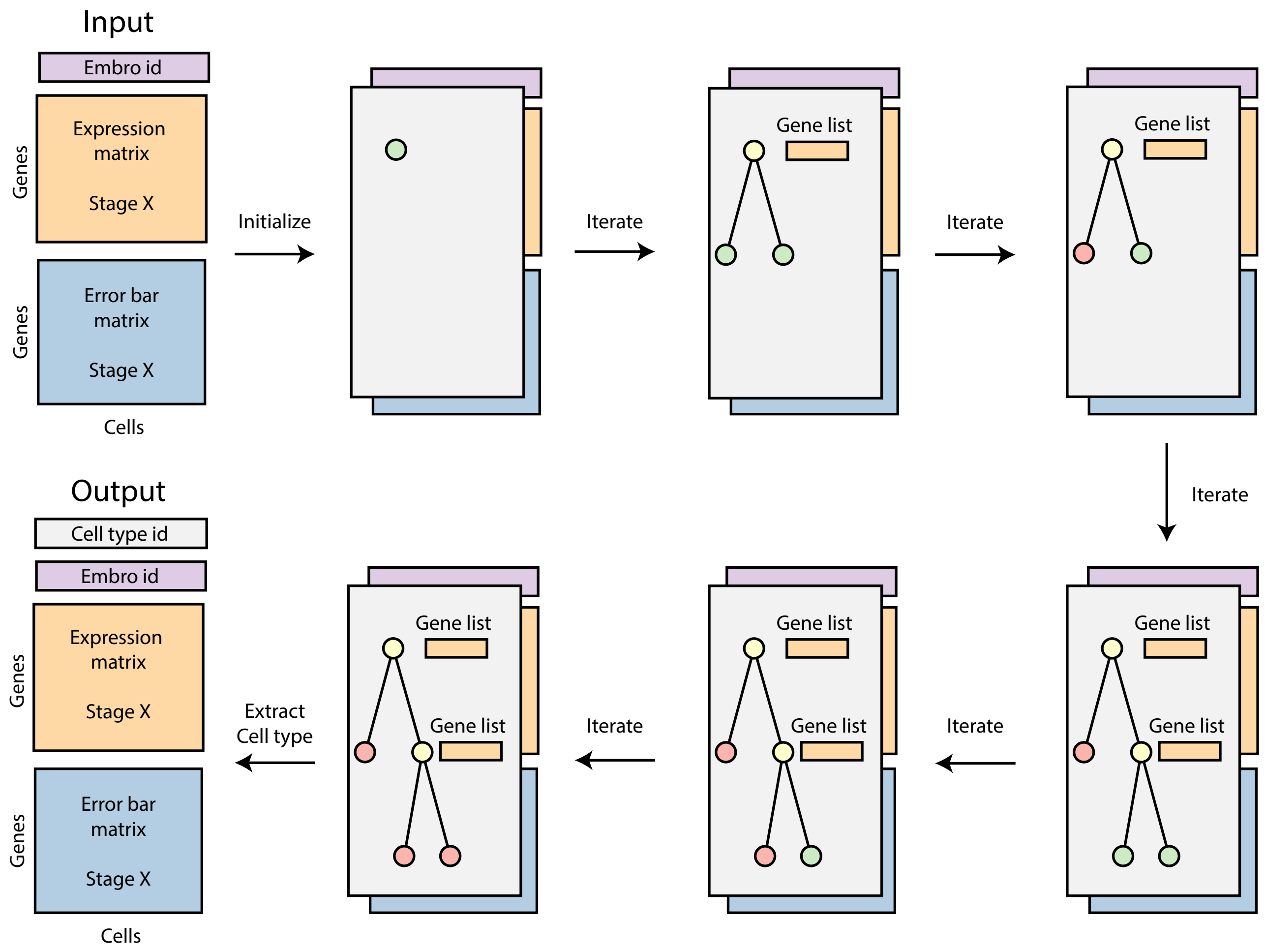


FIG. S5. Summary of data workflow for clustering. We input a matrix of expression with uncertainty and embryo identity. We start with a single active node (green). Running the clustering algorithm, we split into, say, two clusters. We save the genes that were used as well as which points were put into each cluster, create an active node for each cluster, and set the original node to inactive (yellow). If we find that no further clustering is necessary we set a node to terminal (red). We iterate until all nodes are terminal. We output the cell type identity along with the input data, and can optionally save the list of genes used as well as the tree structure.

### E. Results

Results for the overall clustering algorithm are shown for the 8-cell stage in Fig. S6, the 16-cell stage in Fig. S7, the 32-cell stage in Fig. S8 and Fig. S9, and the 64-cell stage in Fig. S10, Fig. S12, and Fig. S11.

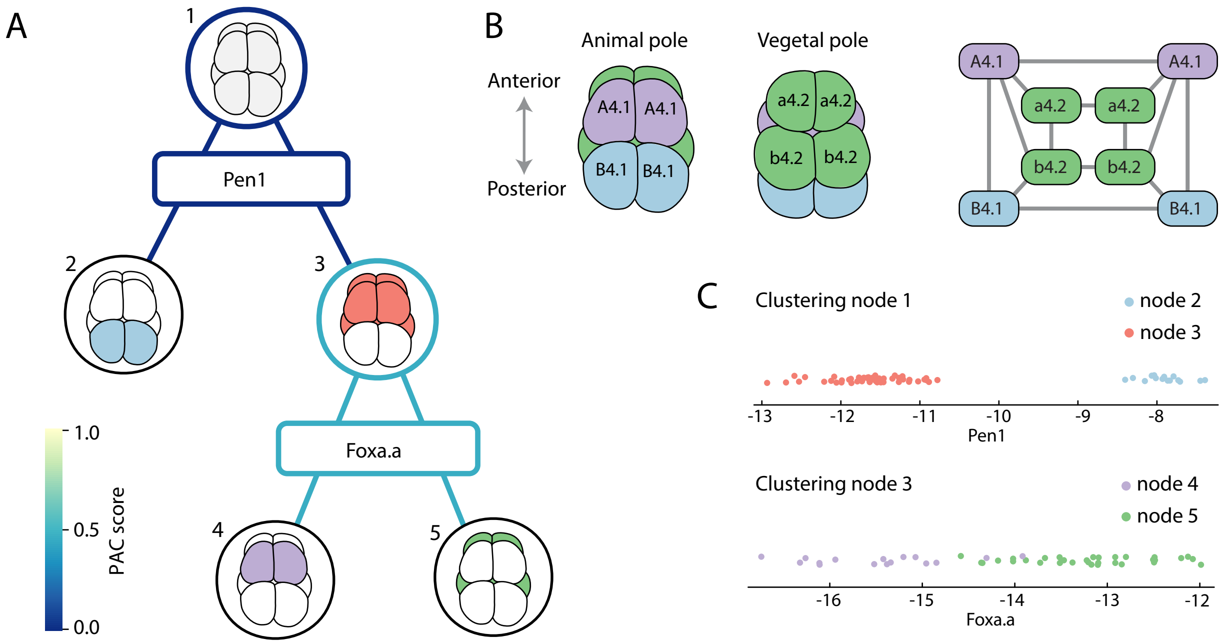


FIG. S6. Hierarchical clustering at the 8-cell stage identifies three cell types. (A) Starting from all cells, the gene *Pen1* is used to split the data into clusters of 2 and 6 cells. Then the gene *Foxa.a* is used to split the 6 cells of node 3 into clusters of 2 and 4. Nodes are either terminal (black) or colored by their PAC score, with the first decision at node 1 resulting in a highly consistent clustering, and the second decision at node 3 resulting in a fairly consistent clustering. (B) Having found the clusters, it is straightforward to identify them with their position in the embryo from in-situ hybridization experiments [10, 18] The gene *Pen1* is a maternal factor localized to the germ line cells, B4.1, whereas the nascent *Foxa.a* expression occurs in the vegetal cells a4.2,b4.2 [10, 18]. The spatial arrangement (left) as well as cell-cell contact network (right) is shown. (C) Gene expression data at the two decision branches of the clustering. Node 1 is clustered using *Pen1* expression (top) and there is a clear separation between clusters. Node 2 is clustered using *Foxa.a*, which has a less clear separation but still results in a statistically significant clustering.

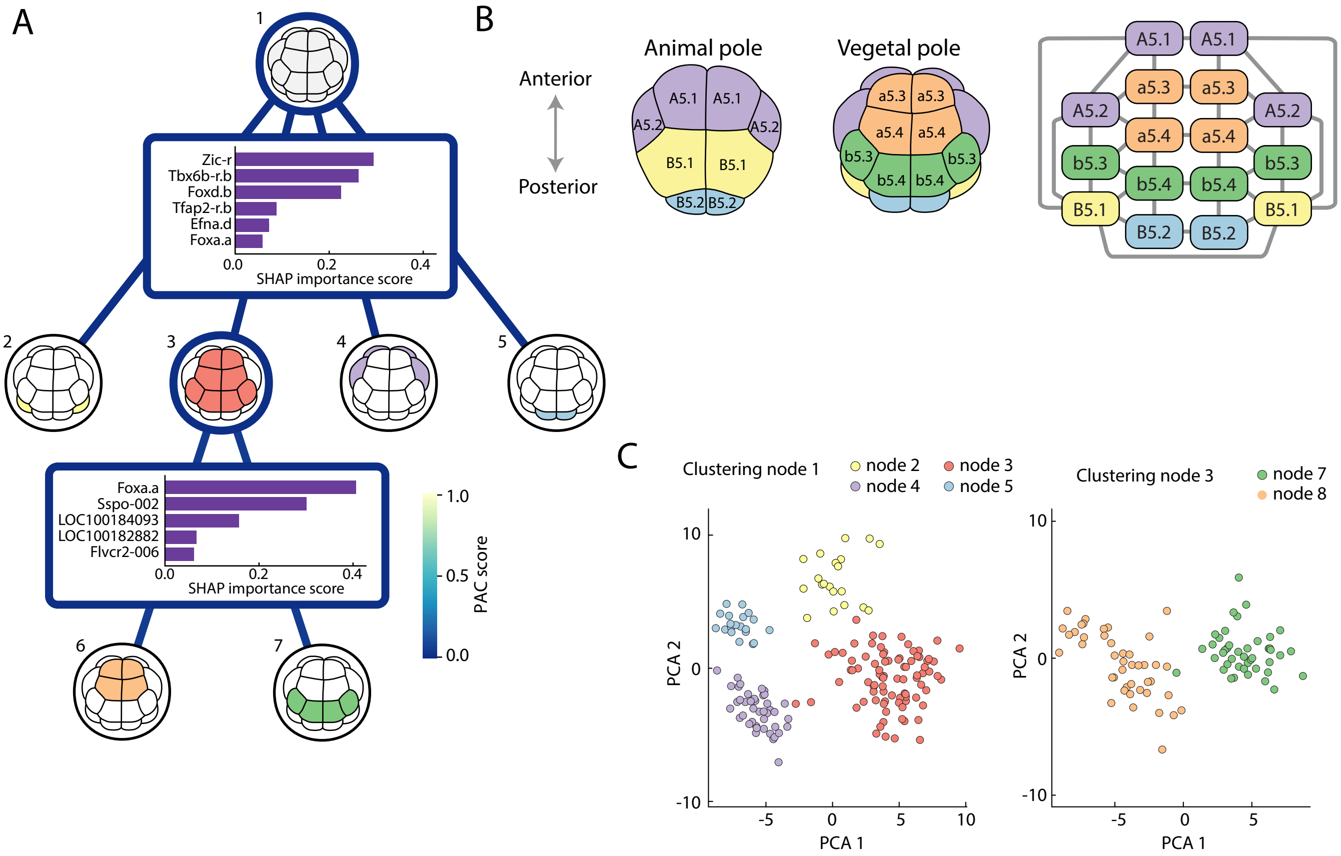


FIG. S7. Hierarchical clustering at the 16-cell stage identifies three cell types (Fig 2B-C of the main text, reproduced here for consistency). (A) Starting from all cells, the 6 genes are used to split the data into 4 clusters. The largest of these clusters is then split into two further with a different set of 5 genes. Nodes are either terminal (black) or colored by their PAC score, with both decisions resulting in a highly consistent clustering. The contributions of each gene used for a split is evaluated using the SHAP importance score [9], genes that contribute more than 5% towards a clustering are shown at each round of clustering. (B) Having found the clusters, we identify their position in the embryo from in-situ hybridization experiments [10, 18]. Localized maternal factors, such as *Pen1*, identify the germ-line cells B5.2 [10, 18]. Lack of *Foxa.a* expression identifies cells b5.3, and b5.4 [10, 18]. Of the remainder, lack of *Foxd.b* identifies a5.3, a5.4 cells [10, 18]. The remaining clusters correspond to the animal cells A5.1, A5.2, and B5.1, which are distinguished by the expression of *Tfap2-r.b* in B5.1 and not the others [18]. The spatial arrangement (left) as well as cell-cell contact network (right) is shown. (C) Gene expression data at the two decision branches of the clustering. Plotting the first two principal components of the 6 genes used for node 1 and 5 genes used for node 2 reveals clear separation between different clusters.

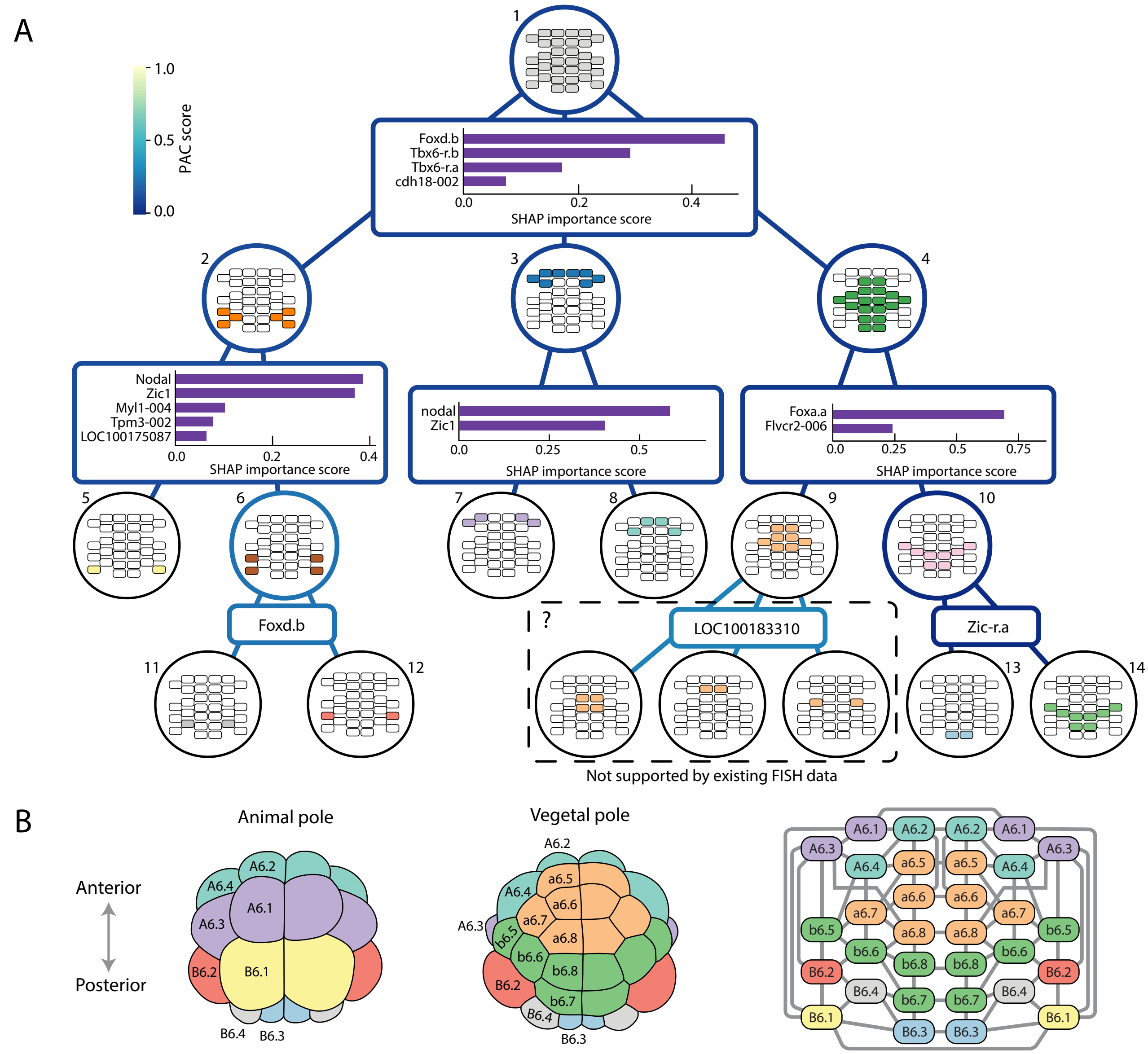


FIG. S8. Hierarchical clustering at the 32-cell stage identifies ten cell types, of which 8 are supported by in-situ hybridization data. (A) Starting from all cells, the data is hierarchically split, first into 3 clusters each of which are then split into further clusters and so on until the algorithm terminates. Nodes are either terminal (black) or colored by their PAC score, with decisions resulting in a highly consistent clustering, with the potential exception of node 9. The contributions of each gene used for a split is evaluated using the SHAP importance score [9], genes that contribute more than 5% towards a clustering are shown at each round of clustering. (B) Having found the clusters, we identify their position in the embryo from in-situ hybridization experiments [10, 18]. In the first round of clustering, high *Tbx6-r.b* identifies B6.1, B6.2, and B6.4 cells (node 2). Of the remainder, high *Foxd.b* identifies A6.1-4 cells (node 3) from a5,b5 and B6.4 cells (node 4). At node 2, high *Nodal* identifies B6.1 cells, and then high *Foxd.b* identifies B6.2 cells from B6.4 cells (node 6). At node 3, A6.1, A6.3 cells express *Nodal*, whereas A6.2, A6.4 cells express *Zic1*. At node 4, *Foxa.a* is expressed in a6 cells, but not b6 or B6.3 cells. Finally, localized maternal factors, such as *Zic-r.a*, identify the germ-line cells B5.3 from b5 cells [10, 18]. Curiously, the algorithm splits the a6 cells into 3 clusters using the uncharacterized gene *LOC100183310* (node 9). Whilst there is uncertainty in this clustering, the algorithm determines it to be significant. However, without supporting in-situ hybridization data to assign cell types to these clusters, we merge them in the remainder, treating node 9 as a terminal node. The overall spatial arrangement (left) as well as cell-cell contact network (right) is shown for the identified cell types.

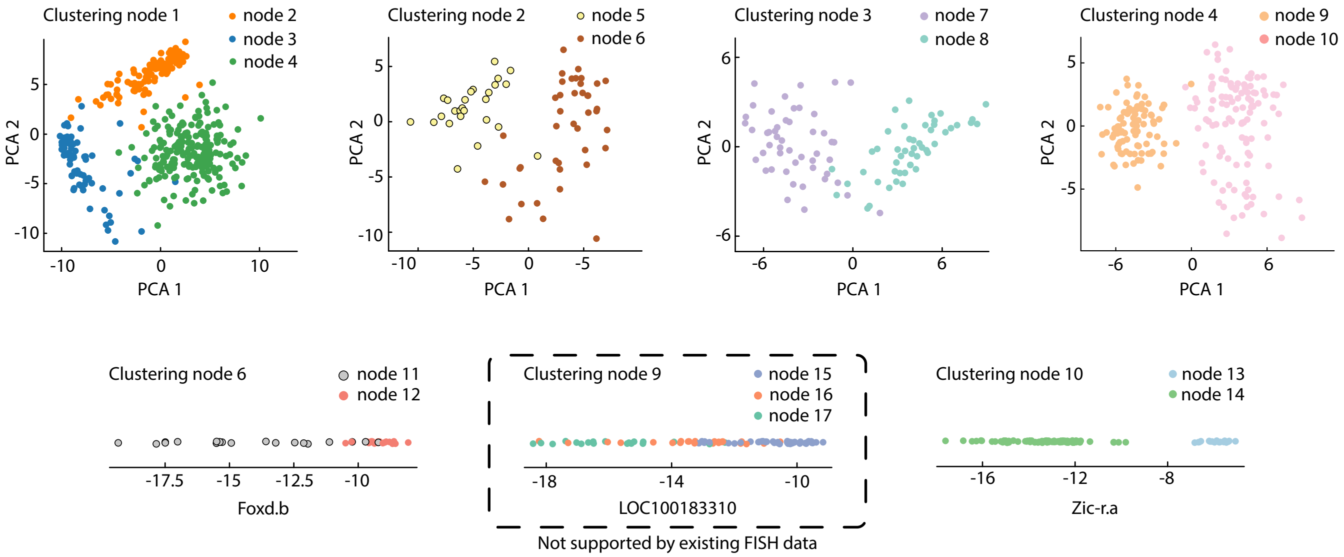


FIG. S9. Hierarchical clustering at the 32-cell stage continued from Fig. S8. The first two principal components are shown from the space of gene expression used when clustering nodes 1, 2, 3, and 4. For nodes 6, 9, 10 only a single gene was used. In every case, with perhaps the exception of node 9, clear clusters are visible.

#### 1. *64-cell stage, caption for Fig. S10*

Hierarchical clustering at the 64-cell stage identifies 20 cell types, of which 15 are supported by in-situ hybridization data. Starting from all cells, the data is hierarchically split, first into 2 clusters each of which are then split into further clusters and so on until the algorithm terminates. Nodes are either terminal (black) or colored by their PAC score, with decisions resulting in a fairly consistent clustering. Unlike in previous rounds of clustering, all 64-cell embryos are missing significant number of cells, in addition to having nosier expression for the cells that are present. The contributions of each gene used for a split is evaluated using the SHAP importance score [9], genes that contribute more than 5% towards a clustering are shown at each round of clustering.

Having found the clusters, we identify their position in the embryo from in-situ hybridization experiments [3, 10, 18]. The first split in the clustering separates cells B7.1-B7.5 and B7.7,B7.8, from the rest (B7.6 are germ line cells), and they are distinguished by the expression of *Tbx6-r.a*. Among these, B7.4 and B7.8 express *Acta1*, of which only B7.4 expresses *Smyd1*. Then, B7.3 and B7.7 are distinguished from B7.1, B7.2, B7.5 by their expression of *Chrd*, and further B7.3 expresses *Mnx*, unlike B7.7.

In addition, animal and vegetal cells can be divided by *Tfap2-r.b*, amongst other genes. For the animal cells, a vs b cells can be distinguished by expression of *Trabd2a*. Then, a7.9,a7.10 can be distinguished from a7.11-a7.16 through the lack of *Mfge8-005* expression. Also, b7.9, b7.10 can be distinguished from b7.11-b7.16 through the expression of *Zic1*.

For the remaining, A7.1-A7.8 and B7.6 cells; only A7.1, A7.2, and A7.5 express *Chrd*. Next, A7.3 and A7.4 are distinguished through their expression of *Bra*, and of these only A7.4 expresses *Chadl-002*. Of the remaining, A7.6, A7.8, B7.6 cells, only A7.8 expresses *Mytf*, and finally A7.6 expresses *NoTrlc* whereas B7.6 does not.

Further splits identified by the clustering, such as splitting a7.9 and a7.10 cells with the *Plg* gene are not supported by existing in-situ hybridization data. It is likely that, due to the many missing cells in each embryo, spurious genes can be found that appear to well separate the data into clusters. If the embryos were more complete, or even if there were more embryos, a spurious separation into clusters would not be possible, as is the case at earlier cell stages. Merging the few clusters not supported by in-situ hybridization data results in a clustering that is consistent with the known biology.

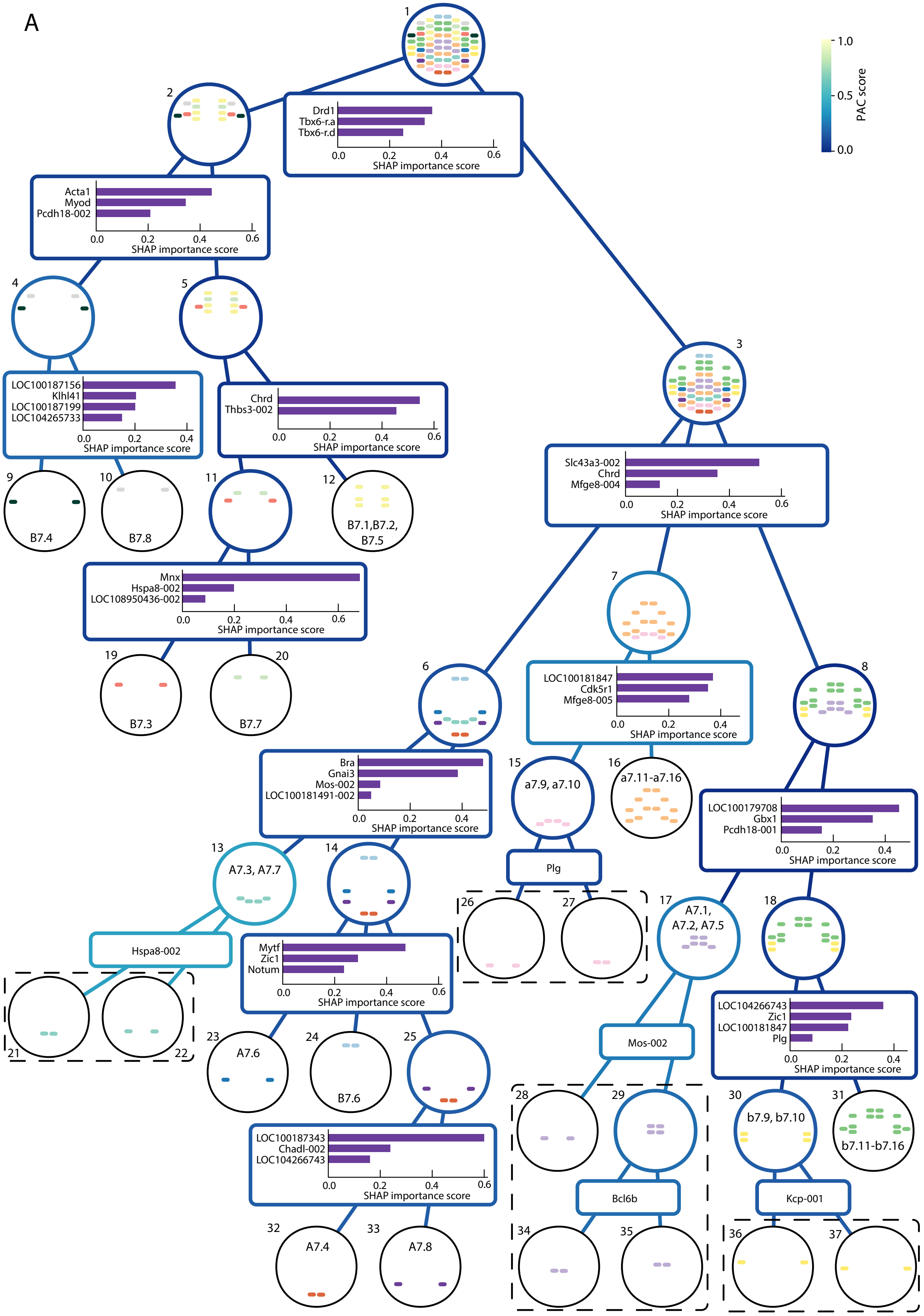


FIG. S10. Hierarchical clustering at the 64-cell stage, see section III E 1 for figure caption.

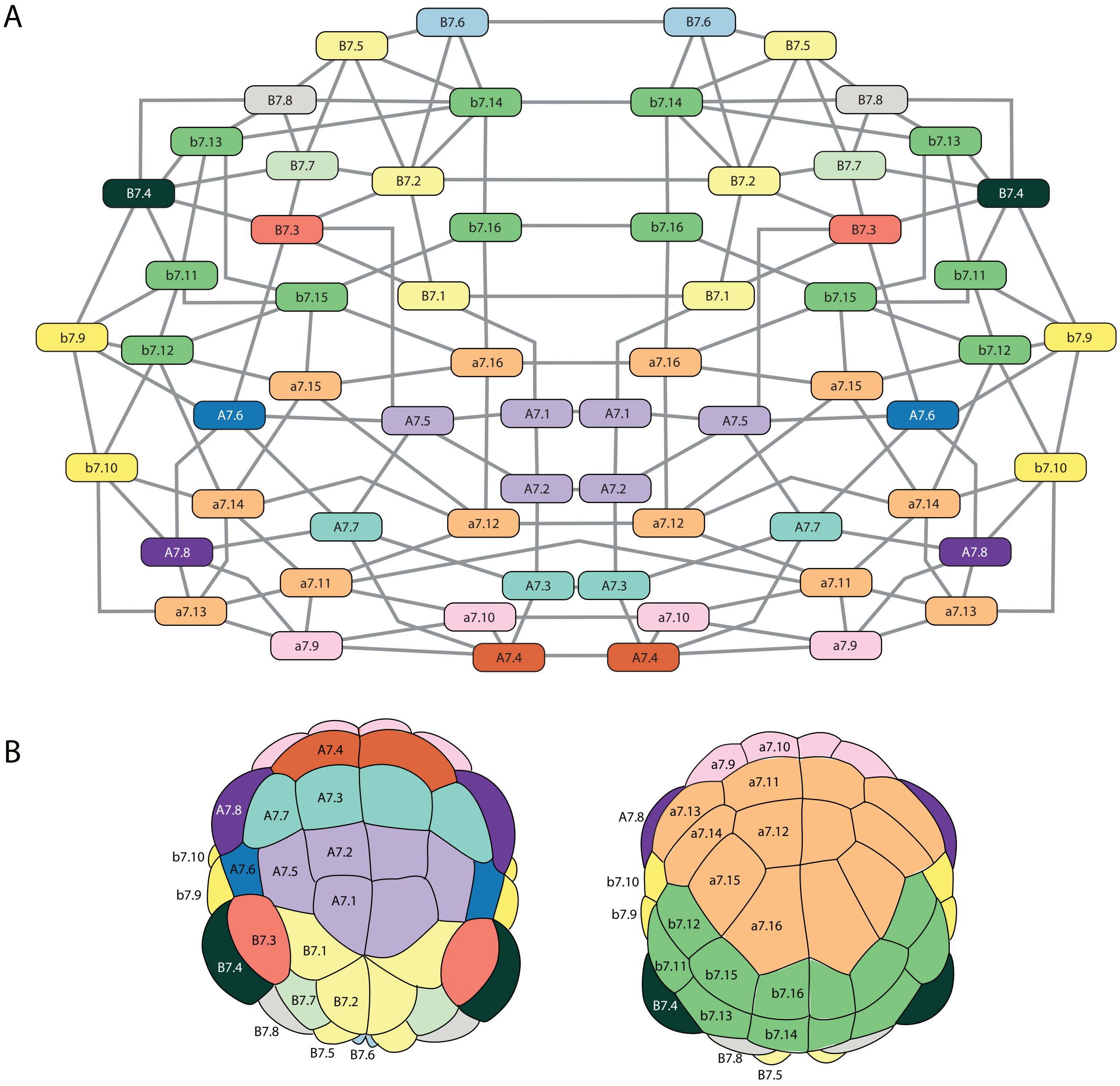


FIG. S11. (A) The cell-cell contact network for the 64-cell stage colored by the cell types identified by clustering. (B) The overall spatial arrangement of cells at the 64-cell stage, again colored by identified cell type.

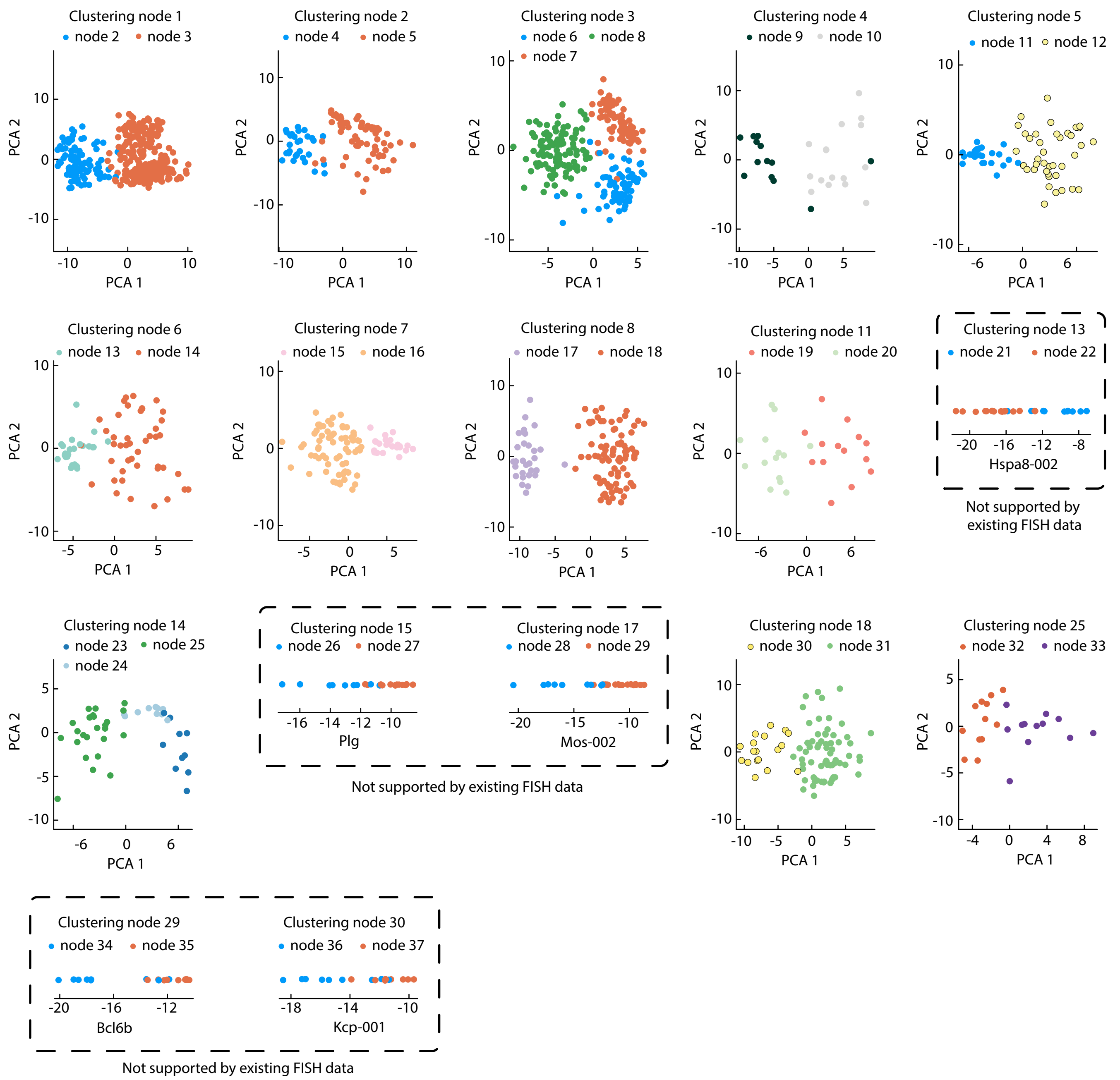


FIG. S12. Hierarchical clustering at the 64-cell stage continued from Fig. S10. The first two principal components are shown from the space of gene expression used when clustering except in the case when only a single gene was used. Rounds of clustering that are not supported by existing in-situ hybridization data are placed within a dotted line.

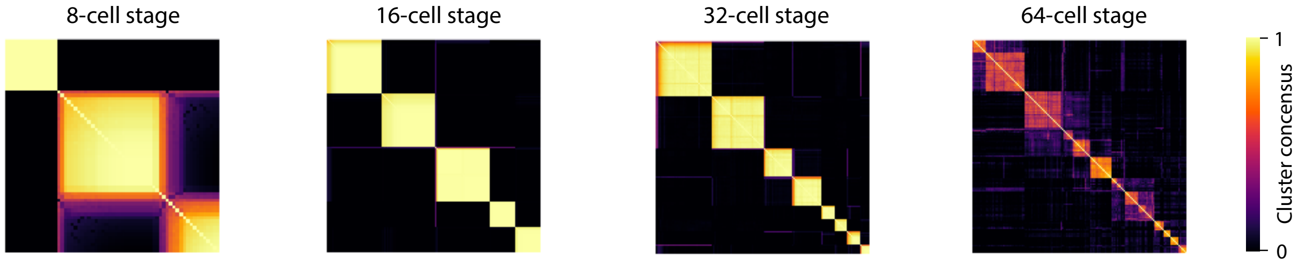


FIG. S13. Overall clustering consistency across each cell stage. In each case, samples were drawn from the posterior distribution and reclustered. The reclustering uses the same hierarchical cluster structure that was found in the main clustering algorithm, i.e. it proceeds hierarchically as before, using the known genes at each stage to split into known cluster sizes. Having done this repeatedly, we color how frequently cells $i$ and $j$ get put into the same cluster, with a perfectly consistent clustering always having values 0 or 1. The 16 and 32-cell stages achieve a very consistent overall clustering. The 8-cell stage clearly identifies the germ line cells, but determining the a4.2, b4.2 cells from A4.1 cells is challenging because difference is the weakly expressed *Foxa.a* gene. The clustering at the 64-cell stage is, while still fairly consistent, much less consistent than the others due to incomplete embryos and noisier expression at the 64-cell stage.

#### F. Overall clustering consistency

Having found an overall recursive clustering, we can ask how consistent is it? Much in the same way that one can rerun a K-means algorithm with the same $K$ for resampled data, we can rerun our combined algorithm for resampled data. For instance, at stage 8, we resample the data, use *Pen1* and the modified K-means to split the data into a cluster of size 2 and a cluster of size 6. We then use the modified K-means to split the cluster of size 6 into a cluster of size 4 and a cluster of size 2 using *Foxa.a*, Fig. S6. Repeating this many times for resampled data gives us a clustering consistency for the overall algorithm. Importantly, while we use the data to discover which genes to use and the sizes of clusters to choose, having found this information, the original data is then not used when the algorithm is run again on resampled data. Therefore, the cluster consistency test gives us a measure of how robust the overall clustering procedure is. The results of the clustering consistency are shown in Fig. S13.

#### G. Comparison with standard pipeline

We can apply a standard preprocessing then clustering [20], as implemented in widely used packages like scanpy [21] and seurat [15], and compare against the clusters found above, although as stressed earlier, a standard approach does not account for the structure of the ascidian embryo. In short, the data is normalized with TPM, highly variable genes are selected [15], these genes are projected onto 10 principal components and then a graph-based clustering algorithm is applied [20]. Taking this data to the 16-cell stage as an example, this standard pipeline finds 9 clusters, more than the 5 that our approach finds, Fig. S14. However, not only do these clusters not correspond with known marker genes, some of them correspond to a specific embryo or pair of embryos, Fig. S14B. This is not surprising as some maternal genes vary significantly between different embryos and an algorithm which is not aware of the nature of cell types will then identify these embryos as separate clusters. Additionally, upon subsampling the data, these clusters are less consistent than the clusters our approach finds, even though when we compute our clustering consistency we also resample from the posterior acknowledging additional uncertainty about the expression measurement, Fig. S14C.

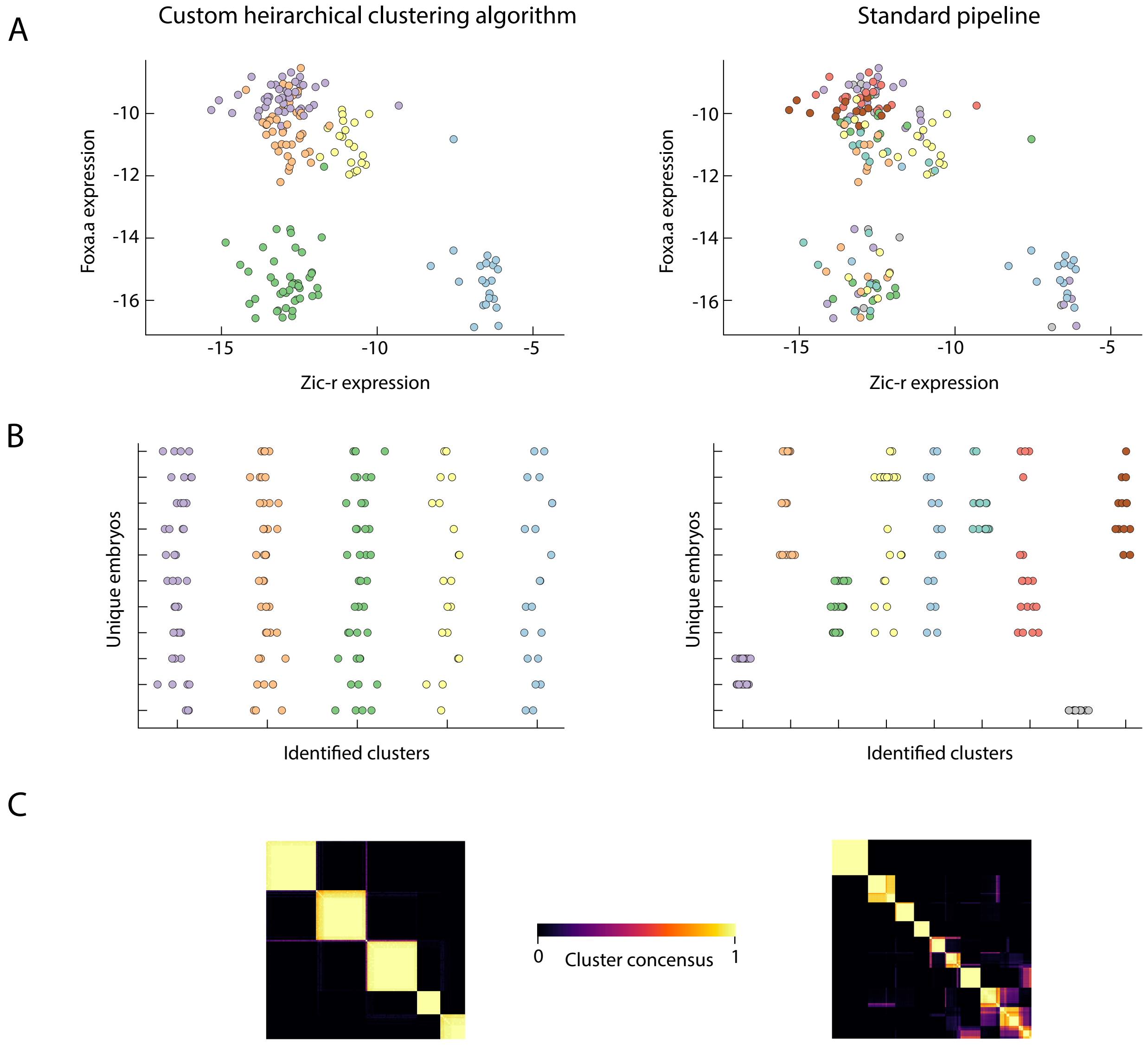


FIG. S14. Comparison of our custom clustering approach (left) with a standard clustering pipeline (right). The 16-cell stage is shown as an example (A) Scatter plot of two key marker genes *Foxa.a* and *Zic-r*, colored by the identified clusters. (B) Cluster identity plotted against embryo identity (with random jitter in x-axis for clarity). Our clustering algorithm, by construction, finds clusters that have cells spread across embryos, whereas the in the standard approach some clusters can correspond entirely to one or two embryos. (C) Overall clustering consistency after subsampling the data by a factor of 0.9. Resampling from the posterior is applied for our clustering approach (left) but not for the standard pipeline (right).

### IV. ANALYSIS OF DATA POST CLUSTERING

For each cell stage, a set of maternal genes, zygotic genes, and differentially expressed genes was determined (Methods). Table S2 shows how many genes were found at each cell stage. As expected, the number of zygotic genes increases with cell stage. The number of differentially expressed genes increases with stage, except at the 64-cell stage where it is smaller than the 32-cell stage. This reflects that many fewer embryos and cells are available at the 64-cell stage and additionally, many more pairwise comparisons between clusters are made, requiring a larger adjustment to the p-value and hence a more stringent requirement to be identified as differentially expressed. For this reason we take the same set of maternal genes at each stage: 2268 genes that are expressed at the 4-cell stage but not differentially expressed at any later stage (Methods). Note that at the 8-cell stage, in addition to the lone zygotic gene (*Foxa.a*), and around 30 maternal factors which are localized to the germ line cells (B4.1), there appear to be around 10 maternal factors that are expressed higher in the somatic cells (A4.1, a4.2, b4.2) than the germ line cells, although the effect is weak.

TABLE S2: Number of identified genes versus stage

| Cell stage | 8 | 16 | 32 | 64 |
|---|---|---|---|---|
| Differentially expressed genes | 46 | 239 | 762 | 634 |
| Zygotic genes | 1 | 92 | 264 | 463 |
| Differentially expressed zygotic genes | 1 | 30 | 138 | 234 |

#### A. Singular values

At each stage, we have matrices $\hat{X}_{cg}$, $\tilde{X}_{cg}$ of maternal, and zygotic genes genes respectively. Fig. S15, shows their singular values after subtracting the mean expression for each gene, $X_{cg} \mapsto Y_{cg} = X_{cg} - \sum_d X_{dg} / \sum_d 1$. Note that we do not plot the zygotic genes at the 8-cell stage as there is only one zygotic gene and hence one singular value.

As discussed in the main text, for the maternal genes, we take $\hat{Y}_{cg} \mapsto \hat{Z}_{cg} = \hat{Y}_{cg} - \nu_{m(c)g}$, with $m(c)$ the mother of the embryo that cell $c$ belongs to, and $\nu_{mg} = \sum_{c \in m} \hat{Y}_{cg} / \sum_{c \in m} 1$ to remove the effect of mother-to-mother variation, Fig. S15. Note that this is redundant for the 8-cell stage, as all embryos come from the same mother. Next, we remove embryo-to-embryo variation, $\hat{Z}_{cg} \mapsto \hat{A}_{cg} = \hat{Z}_{cg} - \eta_{e(c)g}$, with $e(c)$ the embryo that cell $c$ belongs to, and $\eta_{eg} = \sum_{c \in e} \hat{Z}_{cg} / \sum_{e \in m} 1$. Note that removing the mother specific effect was made redundant by removing an embryo specific effect, however it is still interesting to ask how much variation was accounted for by each step and whether the mother specific effect is sufficient.

As for the zygotic genes, we subtract the cell-type specific expression of genes by $\tilde{Y}_{cg} \mapsto \tilde{Z}_{cg} = \tilde{Y}_{cg} - \mu_{t(c)g}$, where $t(c)$ is the cell type that cell $c$ belongs to, and $\mu_{tg} = \sum_{c \in t} \tilde{Y}_{cg} / n_t$ where $n_t = \sum_{c \in t} 1$, Fig. S15.

To remove the effects of time along the WT trajectory, we assume temporal variations enter through $\tilde{Z}_{cg} \approx \tau_{e(c)} \alpha_{t(c)g}$, where $e(c)$ is the embryo corresponding to cell $c$, $t(c)$ the cell type, and $\tau_e$ is an embryo specific time, with $\alpha$ giving the direction of temporal dynamics. We therefore wish to minimize

$$M = \sum_{g,c} (\tilde{Z}_{cg} - \tau_{e(c)} \alpha_{t(c)g})^2. \tag{S6}$$

We can do so by alternately factorizing,

$$M = \sum_e \left[ \tau_e^2 \sum_{g,c \in e} \alpha_{t(c)g}^2 - 2\tau_e \sum_{g,c \in e} \tilde{Z}_{cg} \alpha_{t(c)g} \right] + const. \tag{S7}$$

$$= \sum_{t,g} \left[ \alpha_{tg}^2 \sum_{c \in t} \tau_{e(c)}^2 - 2\alpha_{tg} \sum_{g,c \in e} \tilde{Z}_{cg} \tau_{e(c)} \right] + const. \tag{S8}$$

suggesting an iterative strategy of setting

$$\tau_e = \sum_{g,c \in e} \tilde{Z}_{cg} \alpha_{gt(c)} / \sum_{g,c \in e} \alpha_{t(c)g}^2 \tag{S9}$$

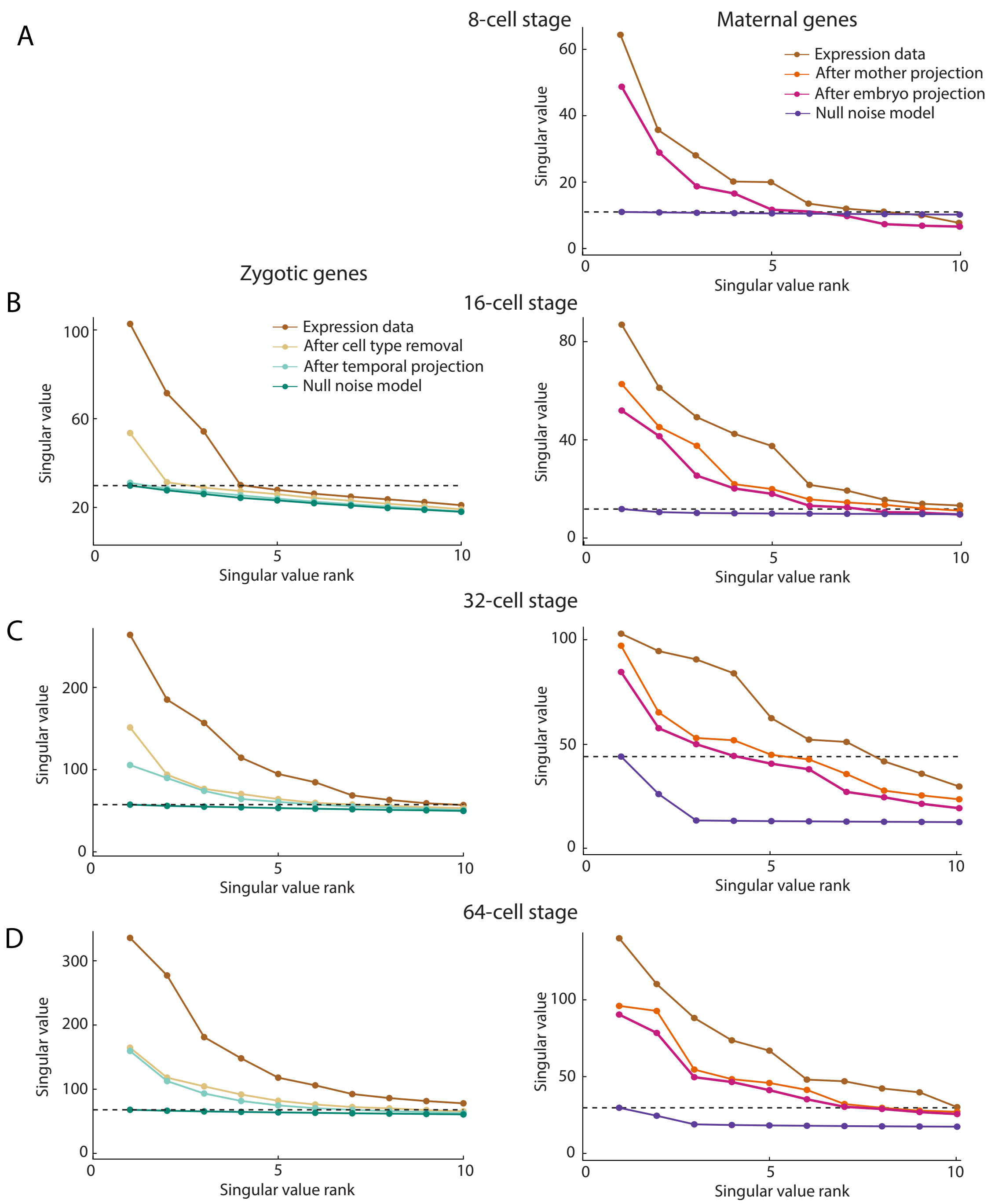


FIG. S15. The largest 10 singular values of zygotic (left) and maternal (right) gene expression matrices for (A) the 8-cell, (B) 16-cell, (C) 32-cell, and (D) 64-cell stages. Singular values are shown for the expression data (with overall expression mean for each gene subtracted), after removing cell type and then temporal projection (zygotic genes) or after projecting to remove the effect of an embryos mother and embryo specific effects (maternal genes). In all cases, accounting for these sources of signal decrease the largest singular values. The data is also shown against a null model, applied by permuting values for each gene after projection has occurred. The largest singular value of the null model is shown (dotted line) and effectively determines the number of singular values that should be kept. Note that all embryos at the 8-cell stage have the same mother, so no mother projection can occur. Also there is only one zygotic gene at the 8-cell stage, the variation in that gene can plausibly be explained by cell type alone, and hence this stage is not included here. Finally, note that while the singular values for the null distribution may not look like that of a random matrix i.e., a Wishart distribution, this is due to the fact that different genes have different variances, after scaling each gene to have unit variance we get random-matrix like statistics, as expected. Part (C) appears in the main text as Fig. 4A-B.

and then

$$\alpha_{tg} = \sum_{g,c\in t} \tilde{Z}_{cg}\tau_{e(c)} / \sum_{g,c\in t} \tau^2_{e(c)}, \tag{S10}$$

together with normalization step of $\sum_{g,t}\alpha^2_{tg} = 1$, to resolve the undetermined scaling degree of freedom. This iterative minimization converges well in practice. Having found $\tau$ and $\alpha$ they are subtracted through $\tilde{Z}_{cg} \mapsto \tilde{A}_{cg} = \tilde{Z}_{cg} - \tau_{e(c)}\alpha_{t(c)g}$. Removing this temporal variation has the effect of removing the largest singular value, in both the 16-cell stage and 32-cell stage, although the effect is less clearly interpretable at the 64-cell stage, Fig. S15.

#### B. Null model for singular values

After subtracting known sources of signal, leaving the matrices $\hat{A}_{cg}$, $\tilde{A}_{cg}$ for the maternal and zygotic genes respectively, there remains variation, Fig. S15. As discussed in the main text, we can estimate how many principal components to keep, assessing how many components represent significant variation and how much can be explained by uncorrelated noise. To do so, we draw a data matrix from the posterior, and remove the embryo effect in the maternal case, and the cell-type and temporal effects in the zygotic case. We compute the singular values of this matrix. We then shuffle this matrix, permuting the values of a given gene between different cells, which preserves the marginal distribution for each gene, but removes correlations between different entries [5]. We then compute the largest singular component and count how many components from the unshuffled data matrix are larger than the largest shuffled component. We do this for many draws from the posterior, getting many values for number of significant components. For the final number we take the $5^{th}$ percentile of these trials, erring on the side of attributing variation to uncorrelated noise.

For Zygotic genes, this gives 0, 0, 5, and 6 components for the 8, 16, 32, and 64-cell stages respectively. For maternal genes, this gives 6, 7, 4, and 7 components for the 8, 16, 32, and 64-cell stages respectively.

#### C. Covariances

To measure covariances, there are two sources of uncertainty we need to account for. The first is the uncertainty in the gene expression measurement itself, which we account for using the Sanity error bars. The second is the uncertainty due to finite measurements, which we account for using bootstrapping.

Suppose we want to compute a covariances from data of the form

$$C^t_{ij} = \frac{1}{n_t}\sum_{c\in t}\langle x_{ci}x_{cj}\rangle \tag{S11}$$

for principal components $i$ and $j$, and cell types $t$ and $t'$, with $t \neq t'$, both variables are assumed to be zero mean. Given experimental data, $x_{ci}$, which has had temporal projection and PCA applied, the empirical estimator for this covariance is

$$\gamma^{tt'}_{ij} = \frac{1}{\sum_e n^e_t n^e_{t'}}\sum_e \sum_{c\in t, c'\in t'} x_{ci}\, x_{c'j}, \tag{S12}$$

where $n^e_t$ is the number of cells of type $t$ in embryo $e$, which may not be a constant across $e$ due to missing cells.

However, as stated earlier, not only do we want the estimate $\gamma^{tt'}_{ij}$, to be robust to the uncertainty in $x$ and the finite data. To do so, we sample a new data set, by drawing embryos with replacement from our data, and then for each embryo drawing from the posterior of gene expression. We then compute the covariance of this new data set. Specifically, we first remove the known sources of signal, for zygotic genes by removing the empirical cell type mean and applying empirical temporal projection, for maternal genes applying the embryo projection. Then we project onto the PCA axis, and compute an empirical covariance, $\gamma^{tt'}_{ij}$. Importantly however, we keep the original PC directions fixed and do not resample those. We do this as similarly sized PCs may switch order under resampling, overestimating our uncertainty at measuring correlations. We can do this resampling procedure hundreds or thousands of times to obtain a posterior distribution for $\gamma^{tt'}_{ij}$. The median of this posterior gives us our best estimate of $\gamma^{tt'}_{ij}$, and the quantiles of the posterior give us an estimate of error bars on this measurement.

After removing the variance and projecting onto the PCs, we have data that should be zero mean. As the mean is estimated from the sample, we could apply the appropriate Bessel correction to the covariance estimate, although for the data set used here it is small, especially in relation to the error bars. The full algorithm is described in Algorithm 6.

Similarly, we can generalize this procedure to compute the measurable $4^{th}$ moments

$$\frac{1}{n_{t_1}n_{t_2}n_{t_3}n_{t_4}} \sum_{c_1\in t_1, c_2\in t_2, c_3\in t_3, c_4\in t_4} \langle x_{c_1 i}\, x_{c_2 j}\, x_{c_3 k}\, x_{c_4 l}\rangle, \tag{S13}$$

where $t_1, t_2, t_3, t_4$ are distinct, by using the empirical estimator

$$\gamma_{ijkl}^{t_1 t_2 t_3 t_4} = \frac{1}{\sum_e n_{t_1}n_{t_2}n_{t_3}n_{t_4}} \sum_e \sum_{c_1\in t_1, c_2\in t_2, c_3\in t_3, c_4\in t_4} x_{c_1 i}\, x_{c_2 j}\, x_{c_3 k}\, x_{c_4 l}, \tag{S14}$$

which we will apply later in Fig. S23.

**Algorithm 6** Procedure for computing covariances. By replacing the median with a $k^{th}$ percentile, effective error bars can be found on the covariances

**Input:** Expression $X$, uncertainty $\epsilon$, principal component projection matrix $P$, $N = 1000$.
**function** COVARIANCEVECTOR($X, \epsilon, P$)
  **for** $s = 1 : N$ **do**
    $X^*, \epsilon^* \leftarrow$ bootstrap $X$, $\epsilon$ by resampling embryos.
    $Y_{ij} \leftarrow$ draw from posterior $X^*_{ij} + \mathcal{N}(0, \epsilon^*_{ij})$
    $Y_{cg} \leftarrow Y_{cg} - \sum_d Y_{dg} / \sum_d 1$
    **if** zygotic genes **then**
      $Z_{cg} \leftarrow Y_{cg} - \mu_{t(c)g}$ \\ remove cell type mean
      $A_{cg} \leftarrow Z_{cg} - \tau_{e(c)}\alpha_{t(c)g}$ \\ remove temporal variation
    **else**
      $Z_{cg} \leftarrow Y_{cg} - \nu_{m(c)g}$ \\ remove maternal variation
      $A_{cg} \leftarrow Y_{cg} - \eta_{e(c)g}$ \\ remove embryo-wide variation
    **end if**
    $B_{ci} \leftarrow \sum_g A_{cg} P_{gi}$
    ${}^s\gamma_{ij}^{tt'} \leftarrow \sum_e \sum_{c\in t, c'\in t'} [B_{ci} B_{c'j}] / \sum_e n_t^e n_{t'}^e,$
  **end for**
  $\gamma_{ij}^{tt'} \leftarrow \text{median}({}^s\gamma_{ij}^{tt'})$
  **return** $\gamma_{ij}^{tt'}$
**end function**

#### D. Shuffle tests

In order to perform the shuffle tests, we now introduce two classes of permutations on the set of all cell identities. The first set $\Sigma$, is defined by $\sigma \in \Sigma$ if $\sigma$ is a permutation and $e(c) = e(\sigma(c))$ for all $c$, i.e. $\sigma$ permutes cells within the same embryo. The second set $\mathcal{E}$ is defined by $\varepsilon \in \mathcal{E}$ if $\varepsilon$ is a permutation and $t(c) = t(\varepsilon(c))$, i.e. $\varepsilon$ permutes cells across embryos within the same cell type.

To apply a permutation $\xi$ we remove the known variance and project onto principle components resulting in a matrix $A_{ci}$, and then shuffle $A_{ci} \mapsto A_{\xi(c)i}$. To fully account for potential overfitting of the temporal projection or cell type identification, we then add back on the sources of variation and treat this as if it were a new data set. All cell-cell correlations either within or across cells have been destroyed by the shuffling in a manner that is independent of the known variance, which has been removed by projection. If this shuffling significantly alters the covariance statistics, specifically the deviation from the average $\Delta$, we know that there are genuine cell-cell couplings remaining after the known variance was removed. The procedures of shuffling before removing the variance and shuffling after removing the variance is precisely described in Algorithm 7.

**Algorithm 7** Procedure for shuffling cells. The function CovarianceVector is found in Algorithm 6.

**Input:** Expression $X$, uncertainty $\epsilon$, permutation set $\Xi$, principal component projection matrix $P$, trials $N = 100$.
$\gamma \leftarrow \textsc{CovarianceVector}(X, \epsilon, P)$
**for** $s = 1 : N$ **do**
  $\xi \leftarrow \xi \in \Xi$ \\ select random shuffle
  $\epsilon^s_{cg} \leftarrow \epsilon_{\xi^{-1}(c)g}$
  $Y_{cg} \leftarrow X_{cg} - \sum_d X_{dg} / \sum_d 1$
  **if** zygotic genes **then**
    $Z_{cg} \leftarrow Y_{cg} - \mu_{t(c)g}$ \\ remove cell type mean
    $A_{cg} \leftarrow Z_{cg} - \tau_{e(c)}\alpha_{t(c)g}$ \\ remove temporal variation
  **else**
    $Z_{cg} \leftarrow Y_{cg} - \nu_{m(c)g}$ \\ remove maternal variation
    $A_{cg} \leftarrow Z_{cg} - \eta_{e(c)g}$ \\ remove embryo-wide variation
  **end if**
  $V_{cg} = X_{cg} - A_{cg}$ \\ total variation removed
  $A_{cg} \leftarrow A_{\xi^{-1}(c)g}$ \\ shuffle
  $X^s_{cg} \leftarrow A_{cg} + V_{cg}$ \\ add back variation
  $\gamma^s \leftarrow \textsc{CovarianceVector}(X^s, \epsilon^s, P)$
**end for**
$\bar{\gamma}^{tt'}_{ij} \leftarrow \left(\gamma^{tt'}_{ij} + \sum_s (\gamma^{tt'}_{ij})^s\right) / (N+1)$
$\sigma^{tt'}_{ij} \leftarrow \sqrt{\left[(\gamma^{tt'}_{ij} - \bar{\gamma}^{tt'}_{ij})^2 + \sum_s ((\gamma^{tt'}_{ij})^s - \bar{\gamma}^{tt'}_{ij})^2\right] / N}$
$\Delta \leftarrow \sum_{i,j,t,t'} \left(\frac{\gamma^{tt'}_{ij} - \bar{\gamma}^{tt'}_{ij}}{\sigma^{tt'}_{ij}}\right)^2$
**for** $s = 1 : N$ **do**
  $\Delta^s \leftarrow \sum_{i,j,t,t'} \left(\frac{(\gamma^{tt'}_{ij})^s - \bar{\gamma}^{tt'}_{ij}}{\sigma^{tt'}_{ij}}\right)^2$
**end for**
**return** $\Delta$, $\Delta^s$

The results of these shuffling tests are shown in Fig. S16 for all cell stages stage. Recall that as the 8 and 16-cell stages for zygotic genes can be explained by uncorrelated variation in expression and temporal variation, there are no significant principle components and hence these stages are not shown.

For the maternal genes, if we only assume mother-to-mother variability, shuffling cells between embryos becomes significant again, Fig. S16. To understand why mother-to-mother variability is not sufficient, we found that while most maternal genes are either not differentially expressed between embryos, or not differentially expressed between embryos with the same mother, there are a small number that are, for examples see Fig. S17. Since embryo-to-embryo variation is sufficient, the fact that mother-to-mother is not suggests the effect is not due to the zygotic program, but rather to these maternal genes which are differentially expressed between embryos with the same mother.

### V. MAXIMUM ENTROPY FRAMEWORK

Recall from the main text, we model a gene expression measurement as a matrix $\boldsymbol{x} = (x_{ci})$, where $i$ indexes the principal components, and $c$ indexes over cells within an embryo. Each experimental realization will give rise to a different $\boldsymbol{x}$, and we wish to understand the distribution $\rho(\boldsymbol{x})$.

We will not have enough data to fully sample $\rho(\boldsymbol{x})$ empirically. Instead, we remain agnostic to the exact underlying mechanism of the system, building a model that preserves the key physical observables, but otherwise assumes as little as possible. Specifically, we construct a model of the distribution $\rho(\boldsymbol{x})$ by maximizing the entropy of $\rho$ subject to constraints that the observables of $\rho$ must match to their experimentally measured values.

The maximum entropy approach has two main components, the first of which is a purely statistical statement.

1. Given some set of observables $\mathcal{C}$, for instance mean or second moments, there are infinitely many distributions consistent with those observables. If we have no further knowledge of the form of the distribution, out of all consistent distributions, we should choose the distribution which is maximally agnostic. We get this distribution by maximizing entropy subject to the constraints that the distribution must agree with the measured observables, $\mathcal{C}$.

2. The underlying mechanisms of our system fix some set of important observed statistics directly. All other statistics follow indirectly from these. A good approximation to the overall distribution can be obtained by maximizing entropy with these important statistics held fixed.

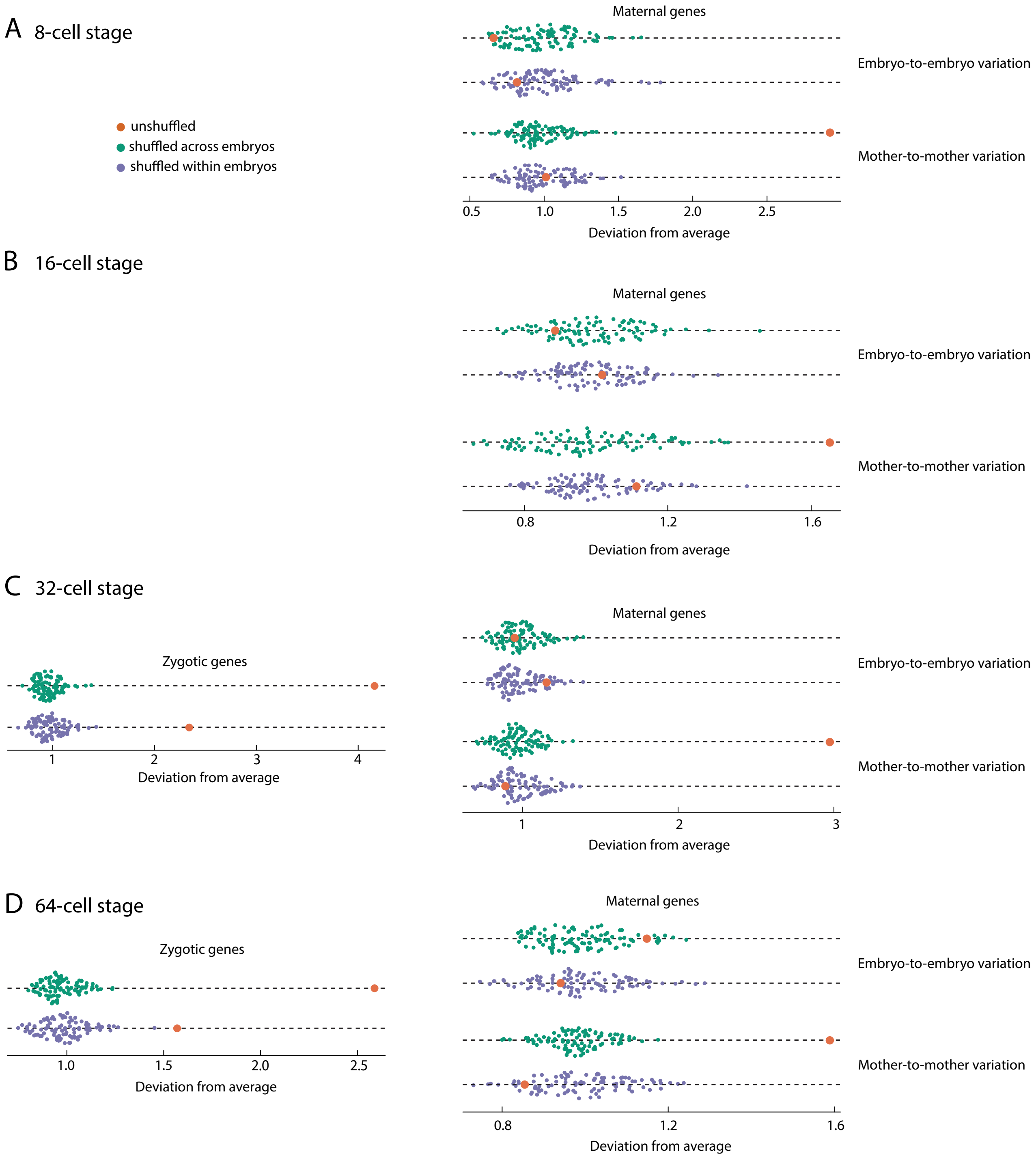


FIG. S16. Shuffle tests for the (A) 8-cell stage, (B) 16-cell stage, (C) 32-cell stage, and (D) 64-cell stage for both maternal and zygotic genes and shuffling within and across embryos. (B) also appears in main text Fig. 4, but reproduced here for consistency. Deviation from average covariance statistics is shown for unshuffled and shuffled data on x-axis (see Alg. 7). For the maternal genes, shuffling is shown after embryo-to-embryo variation was subtracted, and also after only mother-to-mother variation was subtracted. The shuffling is not significant if embryo-to-embryo variation has been accounted for, but is significant if only mother-to-mother variation is accounted for.

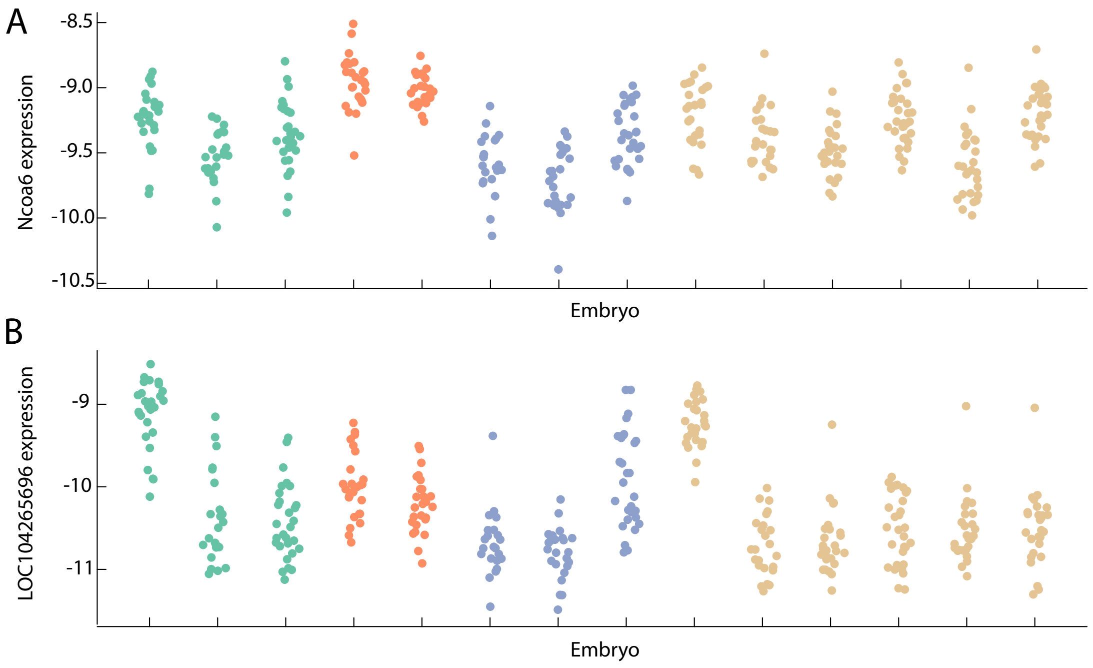


FIG. S17. Examples of maternal and embryo specific effects on gene expression of maternal mRNA at the 32-cell stage. (A) *Ncoa6* expression across different embryos, color corresponds to embryo mother. *Ncoa6* was chosen at random amongst the maternal genes, and shows maternal specific variation as well as some small embryo to embryo variation. (B) *LOC104265696* shows some maternal variation but also shows strong embryo specific variation. A number of genes with embryo-to-embryo variation, like *LOC104265696*, means that mother-to-mother variation is not sufficient to explain variation observed in the data.

Together these can be used as a way to understand which statistics are important and explain the behavior of the system. Constraining a parsimonious choice of variables that are of key physical importance should result in a distribution that closely approximates the real distribution even for statistics that are not directly constrained, and this has been used as a way to test physical hypothesis. For example in a system with multiple firing neurons, constraining simply the rate at which neurons fire did not explain the joint distribution of neurons, whereas constraining the pairwise interactions was sufficient, showing that pairwise interactions are of key importance [2].

#### A. Maximum entropy for embryos

To apply the maximum entropy approach, we must first decide on the observables which we hypothesize are fixed by the underlying mechanism. As discussed in the main text, we will fix covariances of the form

$$C^t_{ij} = \frac{1}{n_t} \sum_{c \in t} \langle x_{ci} x_{cj} \rangle \tag{S15}$$

$$\Lambda^k_{ij} = \frac{1}{|\Omega|} \sum_{p,q} \Omega^k_{pq} \langle x_{pi} x_{qj} \rangle, \tag{S16}$$

$$\Gamma^k_{ij} = \frac{1}{|\Psi|} \sum_{p,q} \Psi^k_{pq} \langle x_{pi} x_{qj} \rangle, \tag{S17}$$

and, after entropy maximization, constraints of this form will result in a Gaussian distribution

$$\rho(\boldsymbol{x}) = (2\pi)^{-n/2} (\det J)^{1/2} \exp\left[ -\frac{1}{2} x_a J_{ab} x_b \right] \tag{S18}$$

$$J_{(pi)(qj)} = \sum_t M^t_{ij} I^t_{pq} + \sum_{k=1}^{n_\Psi} S^k_{ij} \Psi^k_{pq} + \frac{1}{2} \sum_{k'=1}^{n_\Omega} \left( U^{k'}_{ij} \Omega^{k'}_{pq} + U^{k'}_{ji} \Omega^{k'}_{qp} \right), \tag{S19}$$

where here $x_{ci}$ has been flattened into a single vector $x_a$.

We now derive this formula for $J$, starting with a simpler case of only intra-cell covariances fixed before moving to the general case.

#### B. No cell-cell interactions

To start with, and to rederive a known formula from Ref. [8], imagine fixing the correlation matrix between genes within each cell $c$, so $C^c_{ij} = \langle x_{ci} x_{cj} \rangle$, and assume that this quantity is something that can be measured from experiments. In order to maximize the entropy subject to these constraints, we must minimize the following functional,

$$L = \int \mathrm{d}\boldsymbol{x}\, \rho(\boldsymbol{x}) \log \rho(\boldsymbol{x}) \;+ \frac{1}{2} \sum_c \sum_{i,j} M^c_{ij} \int \mathrm{d}\boldsymbol{x}\, \rho(\boldsymbol{x}) x_{ci} x_{cj} + \mu \int \mathrm{d}\boldsymbol{x}\, \rho(\boldsymbol{x}). \tag{S20}$$

We can solve this by taking

$$\frac{\delta L}{\delta \rho} = \log \rho(\boldsymbol{x}) + 1 + \frac{1}{2} \sum_c \sum_{i,j} M^c_{ij} x_{ci} x_{cj} + \mu, \tag{S21}$$

resulting in

$$\rho(\boldsymbol{x}) = \exp\left[ -\frac{1}{2} \sum_c \sum_{i,j} M^c_{ij} x_{ci} x_{cj} \right] / Z, \tag{S22}$$

where the partition function $Z$ is given by

$$Z = \int \mathrm{d}\boldsymbol{x}\; \exp\left[ -\frac{1}{2} \sum_c \sum_{i,j} M^c_{ij} x_{ci} x_{cj} \right]. \tag{S23}$$

In the most general form, flattening $\boldsymbol{x}$ to be indexed as a vector, $\boldsymbol{x} = (x_a)$, we have that

$$Z = \int \mathrm{d}\boldsymbol{x}\; \exp\left[ -\frac{1}{2} x_a J_{ab} x_b \right] = (2\pi)^{n/2} (\det J)^{-1/2}, \tag{S24}$$

and moments can be calculated as

$$\langle x_a x_b \rangle = -2 \frac{\partial}{\partial J_{ab}} \log Z = \frac{\partial}{\partial J_{ab}} \log \det J = (J^{-1})_{ab}, \tag{S25}$$

with

$$J_{(qi)(rj)} = M^q_{ij} \delta_{qr}. \tag{S26}$$

where we have used the matrix identity

$$\frac{\partial}{\partial X_{ij}} \log \det X = (X^{-1})_{ji}, \tag{S27}$$

and the fact that $J$ is symmetric.

In this simple situation, an analytic formula for $J^{-1}$ exists, namely $J^{-1}_{(qi)(rj)} = (M^r)^{-1}_{ij} \delta_{qr}$, and hence

$$C^c_{ij} = \langle x_{ci} x_{cj} \rangle = (M^c)^{-1}_{ij}. \tag{S28}$$

Hence, given the measured $C^c_{ij}$, we can construct $\rho$ through a simple matrix inversion, similar to Ref. [8]. For more complex constraints, a closed form solution is not known.

#### C. Full generality

Previously, we had constrained $C^c_{ij} = \langle x_{ci} x_{cj} \rangle$, for each cell $c$, and maximized entropy. For the sequencing data, as discussed in the main text, the specific identity of each cell can not be determined exactly, but cell types exist and can be identified. Thus, we can measure

$$C^t_{ij} = \frac{1}{n_t} \sum_{c \in t} \langle x_{ci} x_{cj} \rangle, \tag{S29}$$

which we also consider to be fixed by the system, where $t$ indexes over cell types. To enforce this correlation in the Lagrangian, we introduce a term

$$\frac{1}{2} \sum_{i,j,t,p,q} M^t_{ij} I^t_{pq} \langle x_{pi} x_{qj} \rangle, \tag{S30}$$

where $I^t$ is an indicator matrix with $I^t_{pq} = \delta_{pq} \delta_{q \in t}$, and we have introduced the Lagrange multipliers $M^t_{ij}$, with $M^t$ a symmetric matrix.

As for correlations between cells, consider first the symmetric interactions. This could be, for instance, the correlation between an A4.1 cell on the left side and an A4.1 cell on the right side of the embryo. Alternatively, it could be a generic neighbor-neighbor correlation that is not directed across any cell-cell contact. Let $\Psi$ be the adjacency matrix, so that $\Psi_{pq} = 1$ if $p$ and $q$ are cells which share a cell-cell contact relevant for this particular correlation, and $\Psi_{pq} = 0$ otherwise. The measured symmetric correlation between genes $i$ and $j$, given this adjacency matrix is

$$\Gamma_{ij} = \frac{1}{2|\Psi|} \sum_{p,q} \Psi_{pq} \langle x_{ip} x_{jq} \rangle, \tag{S31}$$

where $|\Psi|$ counts the number of edges. To enforce this, we introduce a term into the Lagrangian,

$$\frac{1}{2} \sum_{i,j,p,q} S_{ij} \Psi_{pq} \langle x_{pi} x_{qj} \rangle, \tag{S32}$$

where $S_{ij}$ is a symmetric matrix.

Now consider asymmetric interactions. For instance, consider the correlation between PC $i$ in cell $p$ and PC $j$ in cell $q$, $\langle x_{pi} x_{qj} \rangle$. Due to the left-right symmetry, we also have to include the correlation between gene $i$ in the mirror of cell $p$ and gene $j$ in the mirror of cell $q$. Introducing an asymmetric adjacency matrix, $\Omega_{rs}$ where $\Omega_{rs} = 1$ for a directed contact between cell $r$ and cell $q$, and $\Omega_{rs} = 0$ else, the correlation is

$$\Lambda_{ij} = \frac{1}{|\Omega|} \sum_{p,q} \Omega_{pq} \langle x_{pi} x_{qj} \rangle, \tag{S33}$$

and we enforce this correlation by adding a term to the Lagrangian of

$$\frac{1}{2} \sum_{i,j,p,q} U_{ij} \Omega_{pq} \langle x_{pi} x_{qj} \rangle, \tag{S34}$$

where $U$ need not be symmetric.

In total, if we are enforcing $n_\Psi$ symmetric correlations, and $n_\Omega$ asymmetric correlations, the total Lagrangian becomes

$$\begin{aligned} L = &\int \mathrm{d}\boldsymbol{x}\, \rho(\boldsymbol{x}) \log \rho(\boldsymbol{x}) \;\; + \mu \int \mathrm{d}\boldsymbol{x}\, \rho(\boldsymbol{x}) \\ &+ \frac{1}{2} \sum_{i,j,p,q} \left( \sum_t M^t_{ij} I^t_{pq} + \sum_{k=1}^{n_\Psi} S^k_{ij} \Psi^k_{pq} + \sum_{k'=1}^{n_\Omega} U^{k'}_{ij} \Omega^{k'}_{pq} \right) \times \int \mathrm{d}\boldsymbol{x}\, \rho(\boldsymbol{x}) x_{pi} x_{qj}. \end{aligned} \tag{S35}$$

This leads to

$$\rho(\boldsymbol{x}) = (2\pi)^{-n/2} (\det J)^{1/2} \exp \left[ -\frac{1}{2} \sum_{a,b} J_{ab} x_a x_b \right], \tag{S36}$$

with

$$J_{(pi)(qj)} = \sum_t M^t_{ij} I^t_{pq} + \sum_{k=1}^{n_\Psi} S^k_{ij} \Psi^k_{pq} + \frac{1}{2} \sum_{k'=1}^{n_\Omega} \left( U^{k'}_{ij} \Omega^{k'}_{pq} + U^{k'}_{ji} \Omega^{k'}_{qp} \right), \tag{S37}$$

where we have symmeterized the final term to ensure $J$ is symmetric. There is no known analytic formula for the inverse of $J$ in terms of the $M, S$ and $U$ matrices. For this model, as earlier, we have that

$$\langle x_{pi} x_{qj} \rangle = (J^{-1})_{(pi)(qj)}. \tag{S38}$$

We arrived at this form of $J$ by maximizing the entropy with Lagrange multipliers. We must choose these Lagrange multipliers so that the correlations of the model match those of the data. We could think of our task of finding the parameters $\{M^t, S^k, U^k\}$ as a minimization problem where we must minimize the functional

$$\begin{aligned} \tilde{L} = & \sum_{i,j,t} \left( C^t_{ij} - \frac{1}{n_t} \sum_{r \in t} (J^{-1})_{(rj)(ri)} \right)^2 \\ & + \sum_{i,j,k} \left( \Gamma^k_{ij} - \frac{1}{2|\Psi|} \sum_{q,r} \Psi_{qr} (J^{-1})_{(ri)(qj)} \right)^2 \\ & + \sum_{i,j,k'} \left( \Lambda^{k'}_{ij} - \frac{1}{|\Omega|} \sum_{q,r} \Omega_{qr} (J^{-1})_{(ri)(qj)} \right)^2 . \end{aligned} \tag{S39}$$

There is an issue with this formulation, we do not measure $\Gamma$ or $\Lambda$ as we can not order the cells in space. Instead we measure correlations like $\Gamma^{tt'}_{ij} = (\sum_{r \in t, q \in t'} \langle x_{ri} x_{qj} \rangle)/n_t n_{t'}$, where we have defined $\Gamma^{tt'}_{ij}$, and $t$ and $t'$ are cell types. To match these observables, we minimize the functional

$$\begin{aligned} \tilde{L}(\{M^t, S^k, U^k\}) = & \sum_{i,j,t} \left( C^t_{ij} - \frac{1}{n_t} \sum_{r \in t} (J^{-1})_{(rj)(ri)} \right)^2 \\ & + \sum_{i,j,t<t'} \left( \Gamma^{tt'}_{ij} - \frac{1}{n_t n_{t'}} \sum_{r \in t, q \in t'} (J^{-1})_{(ri)(qj)} \right)^2 , \end{aligned} \tag{S40}$$

where we minimize over $\{M^t, S^k, U^k\}$. Assuming, for now, that a solution exists with minimum $\tilde{L} = 0$, then we have found that a maximum entropy model, fit to correlation matrix $C$ and some *a priori* unknown $\Lambda$, $\Gamma$, which we can calculate from $M$, $S$, and $U$.

#### D. Initial guess

Whilst we do not know the true $\{M^t, S^k, U^k\}$, to find them numerically, we must start with an initial guess and perform gradient descent to minimize $\tilde{L}$. We derive an initial guess from the limit of weak coupling, assuming that $\Gamma$ and $\Lambda$ are in some sense small. If $\Lambda$ and $\Gamma$ are in fact zero, we see that $J_{(pi)(qj)} = \sum_t (C^t)^{-1}_{ij} I^t_{pq}$ satisfies

$$(J^{-1})_{(pi)(qj)} = \sum_t C^t_{ij} I^t_{pq}, \tag{S41}$$

and sets $\tilde{L} = 0$, meaning that we have found an exact (known) solution in this case. If necessary, more sophisticated ways of initializing are available [14].

### VI. INFERENCE

For the models with interactions between spatial neighbors, interactions between only sister cells, and embro-wide cell-cell interaction, main text Fig. 6 shows how well inter-cell covariances are fit by the model. Since the model is fitted to both the inter-cell and intra-cell covariances, the fit to both is shown here for reference in Fig. S18.

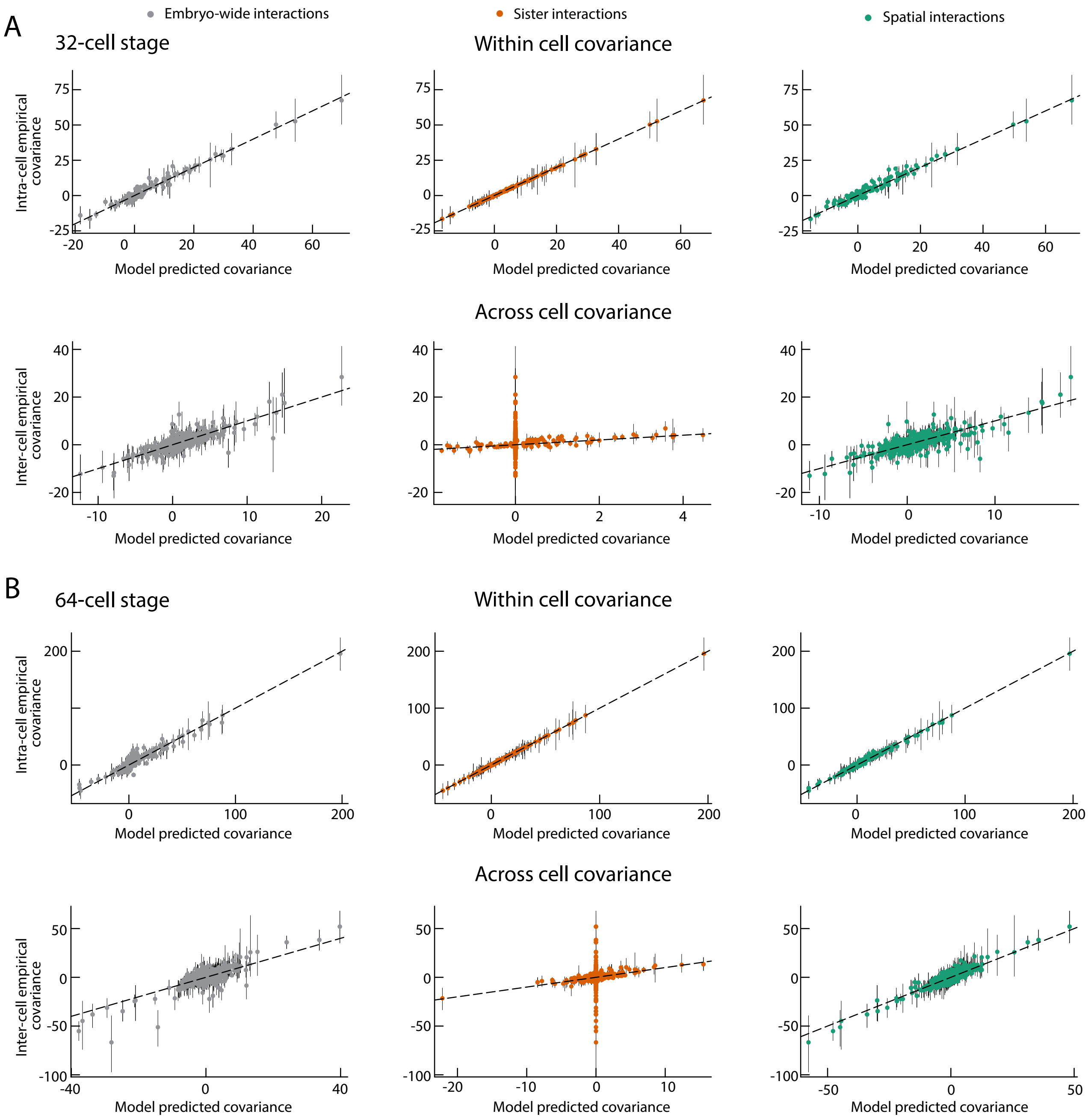


FIG. S18. Comparing the model predicted covariances, computed from a maximum entropy model, against empirical covariances measured from data. Here, we compare the intra-cell covariances between principal components in the same cell (top) as well as the inter-cell covariances, measured between cell types (bottom). Comparison shown for the (A) 32-cell stage, and (B) 64-cell stage. The maximum entropy models assume that all cells within an embryo directly interact (gray) only sisters interact (orange), or only spatial neighbors interact (green). Error bars are from the $25^{th}$ to $75^{th}$ percentile of covariance estimates, black dashed line is $x = y$. The inter-cell covariances comparison also appears in main text Fig. 6. Both inter- and intra-cell covariances are used to fit the maximum entropy models.

For the full model, we allow for all plausible interaction terms. Specifically, we take a $\Psi^k$ for a generic embryo wide interaction between all cells, one for an interaction between all sisters, and one for all spatial neighbors, all with corresponding $S^k$. We then take a specific $\Omega^k$ and corresponding $U^k$ matrix for every pair of spatial neighbors where $\Omega^k$ also couples the corresponding left-right mirror symmetric pair on the other side of the embryo. In the case where one neighbor is on the left and the other is on the right side of the embryo, we use a symmetric adjacency matrix $\Psi^k$ to enforce the left-right symmetry. Thus every pair of neighboring cells can have a unique interaction in addition to the generic sister, neighbor, and embryo wide interactions. However, we have limited experiments, and including all of these parameters would result in an overfitted model, therefore we will use regularization to find the sparsest model which can explain the data.

Here, we adapt the procedure of Ref. [19], to our specific problem. Note that we start with a model with free parameters that are to be fit, $\boldsymbol{p} = \{M^t, S^k, U^k\}$, where $M^t$ can be dense, but the others should be sparse. We will end with a ranking of the sparse terms by importance. The steps are as follows:

1. For a given $L_1$ regularization parameter, $\lambda$, generate covariances from a sample of the data (bootstraped + Sanity error bars) and fit a max ent model by minimizing the objective

$$L_{reg}(\mathbf{p}) = \tilde{L}(\mathbf{p}) + \lambda \left( \sum_{i \leq j,k} |S_{ij}^k| + \sum_{i,j,k} |U_{ij}^k| \right). \tag{S42}$$

   Do this $k$ times.

2. For each parameter value $p_i$ which is being regularized ($S$ and $U$ matrices) compute the coefficient of variation,

$$CV = \frac{iqr(p_i)}{median(p_i)}, \tag{S43}$$

   across the $k$ trials.

3. For this particular value of $\lambda$ the values of the CV provide a ranking for the importance of terms, where a small CV value represents a term that is consistently non-zero and is an important term to keep.

4. Do this for a number of different $\lambda$ values. Aggregate the different rankings with the Schulze method [16].

5. Starting with the sparsest model of no terms, we can train with increasing numbers of terms, as defined by the ranking. Based on how well each model fits the data, we can decide how many terms to keep.

Specifically, we choose $k = 100$, and take 8 regularization parameters $\lambda$ for each stage. We choose the value of the regularization parameters by first fitting models to the non-bootstrapped and not resampled covariance values with various regularization parameters, $\lambda$. As $\lambda \to 0$, the objective function plateaus as effectively there is no regularization, whereas when $\lambda \to \infty$, the objective function also plateaus as the $S$ and $U$ terms are all set to zero. On a log scale in lambda, this looks like a sigmoid curve, and we choose the $\lambda$ values for the above algorithm to be spaced evenly on a log scale and approximately spanning the range in which the sigmoid varies.

#### A. Inference results

The first 25 most important terms are listed in Table. S3 for the 32- and 64-cell stages. The inferred model for the 64-cell stage, along with the first 3 corresponding collective modes is shown in Fig. S19. The first 4 collective modes for a model without interactions at the 32-cell stage is shown in Fig. S20.

### VII. SYNTHETIC DATA AND INFERENCE TESTING

#### A. Synthetic data

In order to assess the inference procedure, we seek to test it on data with a ground truth. We can do so by generating a synthetic data set with known sparsity. For instance, we can take the sparse model that explains $> 90\%$ of the variance in the covariance measurements ($\eta > 0.9$). This model corresponds to a probability distribution $\rho(\mathbf{x})$ from which we can sample. We take finite samples of this distribution in order to generate a realistic synthetic data set. We will then apply the inference procedure to the synthetic data set and assess whether we recover the true sparse interactions that were known to generate it.

Our inference pipeline starts with a expression data set $X$, together with error bars $\epsilon$, allowing us to sample from the posterior. As outlined in Section IV, first the known variation, such as cell types and temporal variation, is subtracted, $A = X - V$, then the remainder $A$ is projected onto principal components, $A \mapsto PA$ from which covariances can be computed. The distribution $\rho$ models variation in this principal component space. To generate a realistic data set,

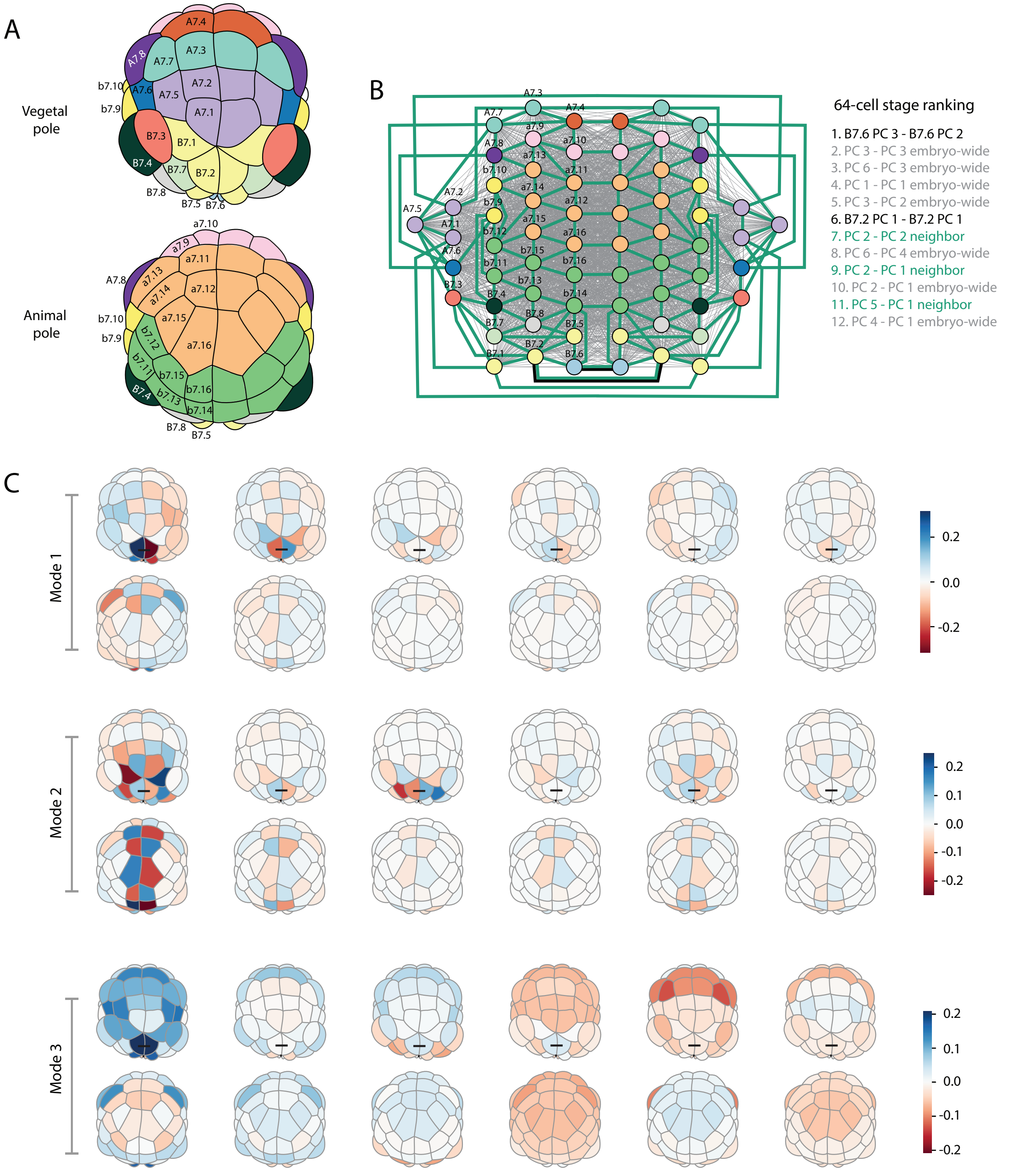


FIG. S19. Collective modes of variation and inferred interactions for the 64-cell stage. (A) Animal and vegetal views of the embryo at the 64-cell stage colored by cell types. (B) Direct interactions of regularized statistical physics model at the 64-cell stage. Using regularization we rank all possible interaction terms by importance. A 12 term model can explain 90% of the variation in the covariance values, with embryo-wide (grey lines) and neighbor (green) cell-cell interactions amongst the most important terms, as well as specific cell-cell interactions (black lines). For ranking of terms see Table S3. (C) Principal modes of variation from inferred regularized model. The first three principal directions are shown, which together capture over 90% of the variance in the model. Black lines show direct interactions between specific cells.

| | 32-cell stage | 64-cell stage |
|---|---|---|
| 1. | PC 4 - PC 1 neighbor | B7.6 PC 3 - B7.6 PC 2 |
| 2. | B6.3 PC 1 - B6.3 PC 1 | PC 3 - PC 3 embryo-wide |
| 3. | B6.1 PC 1 - B6.1 PC 1 | PC 6 - PC 3 embryo-wide |
| 4. | B6.1 PC 4 - B6.1 PC 2 | PC 1 - PC 1 embryo-wide |
| 5. | PC 5 - PC 3 embryo-wide | PC 3 - PC 2 embryo-wide |
| 6. | a6.7 PC 1 - a6.8 PC 2 | B7.2 PC 1 - B7.2 PC 1 |
| 7. | PC 5 - PC 4 embryo-wide | PC 2 - PC 2 neighbor |
| 8. | PC 4 - PC 1 embryo-wide | PC 6 - PC 4 embryo-wide |
| 9. | a6.7 PC 3 - a6.8 PC 2 | PC 2 - PC 1 neighbor |
| 10. | PC 4 - PC 2 embryo-wide | PC 2 - PC 1 embryo-wide |
| 11. | b6.8 PC 3 - b6.7 PC 4 | PC 5 - PC 1 neighbor |
| 12. | a6.8 PC 1 - a6.8 PC 1 | PC 4 - PC 1 embryo-wide |
| 13. | PC 2 - PC 2 embryo-wide | PC 6 - PC 1 embryo-wide |
| 14. | B6.1 PC 3 - B6.1 PC 3 | b7.11 PC 1 - b7.12 PC 4 |
| 15. | A6.1 PC 1 - A6.2 PC 4 | PC 3 - PC 3 neighbor |
| 16. | A6.3 PC 1 - B6.2 PC 5 | PC 1 - PC 1 neighbor |
| 17. | PC 5 - PC 2 embryo-wide | B7.6 PC 1 - B7.5 PC 4 |
| 18. | b6.6 PC 1 - b6.7 PC 3 | B7.6 PC 2 - B7.6 PC 1 |
| 19. | A6.2 PC 2 - A6.4 PC 2 | B7.1 PC 1 - B7.3 PC 2 |
| 20. | A6.3 PC 3 - B6.1 PC 1 | A7.4 PC 2 - A7.4 PC 2 |
| 21. | PC 5 - PC 1 embryo-wide | B7.6 PC 5 - B7.6 PC 1 |
| 22. | PC 1 - PC 1 neighbor | PC 5 - PC 3 embryo-wide |
| 23. | a6.6 PC 1 - a6.8 PC 4 | B7.6 PC 5 - B7.6 PC 2 |
| 24. | a6.8 PC 2 - a6.8 PC 2 | a7.15 PC 1 - a7.14 PC 3 |
| 25. | a6.7 PC 2 - a6.8 PC 2 | B7.6 PC 4 - B7.6 PC 4 |

TABLE S3. Ranking of interaction terms, as inferred by regularization procedure, the first 25 terms for the 32- and 64-cell stages are shown. For the 32-cell stage, a 17-term model was taken to infer collective modes, for the 64-cell stage a 12-term model was taken.

we sample from $\rho$ as many times we have experimental embryos, deleting cells from the synthetic data in the same places where cells are missing in the experimental data. We take the resulting data matrix $B_s$, and transform it to $X_s = (P^+ B_s) + V$, where $P^+$ is the pseudoinverse, inverting the projection from the latent space to the full space, and $V$ is the matrix of removed variation computed from the full data set. We pair $X_s$ with the error bars from the experimental data in order to get a synthetic data set. Note that the temporal projection and principal component projections are not trivially undone as any resampling happens before they are applied.

We then put this synthetic data through the same inference pipeline to get a ranking of terms. For data coming from a model with $k$ terms, those terms should ideally be identified as the most important $k$ terms by the inference procedure, or at least identified as amongst the most important. Broadly speaking we find the procedure identifies the true sparse terms as amongst the most important ones, even if it does not do so perfectly, Fig. S21. This suggests that we can trust that should a true sparse set of interactions exist, these terms will be identified as amongst the most important, even if some additional spurious terms rank highly also. Moreover, we can ask how the robustness of the inference changes with the size of data, generating a synthetic data set with, say, ten times as many embryos as the experimental data, in which case the ability to recover the true sparse terms is improved, Fig. S21.

#### B. Robustness of inferred modes

We have established that the inferred interaction terms are robust, but the exact ranking may change and it is possible that a few spurious interactions are included among the top ranked terms. The question then becomes whether the inferred modes are robust to a change of the interaction terms. We can test this by taking the top 3 modes from the 17-term model, which explain around 80% of that model's variance, and ask how much of the variance do these modes explain for different models. Doing this for the first 25 models of varying numbers of interaction terms from 0 to 24, shows that for many models which fit the empirical data well, the 3 modes explain around 20% of the variance, Fig. S22. We can repeat this for models using the ranking from the synthetic data, which does not agree perfectly with the ranking from the experimental data (Fig. S21). Nevertheless, the three modes from the original 17-term model still explain around 20% of the variance of these new models, Fig. S22. Moreover, while the exact direction of the mode vector changes, inspecting modes coming from different models often finds left-right and animal-vegetal like modes appearing in the top three, suggesting these are robust patterns of variation.

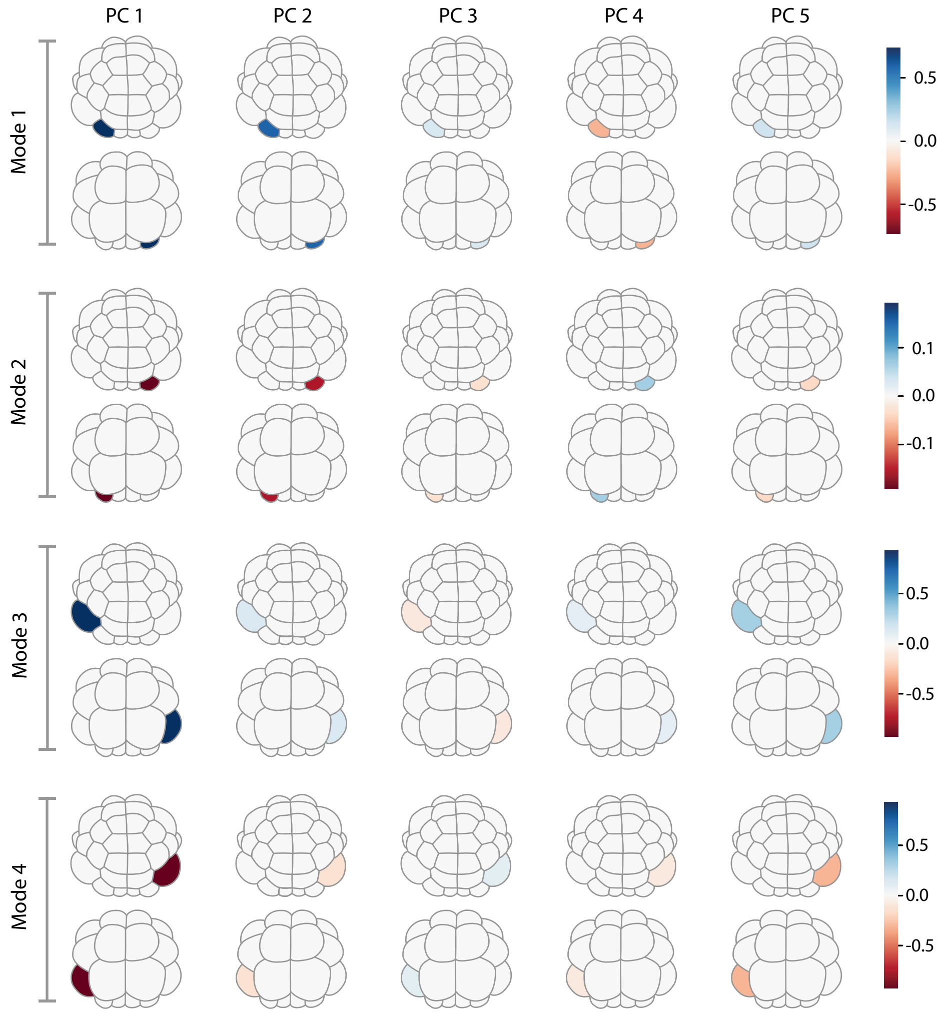


FIG. S20. Principal modes of variation inferred from a model with no cell-cell interaction at the 32-cell stage. Without cell-cell interactions, the modes of variation represent variation in one cell at a time, with modes coming in left-right pairs. The top 4 modes explain around 30% of the variance in the model.

#### C. Gaussian assumption

Another way to test whether our model has predictive power is to compare how it performs on statistics that we did not use to fit with. Since we used all of the $2^{nd}$ order moments that we could measure in fitting, we must turn to higher order moments, such as the measurable $4^{th}$ order moments. Further, in any Gaussian model the measurable $4^{th}$ order moments can be computed from the measurable $2^{nd}$ order moments through Isserlis' theorem. Thus any Gaussian model which fits the $2^{nd}$ order moments well will have the same prediction for the $4^{th}$ order moments. Therefore, to test whether our models perform well at predicting unknown $4^{th}$ order moments, we are really asking whether the $4^{th}$ order moments can be explained by a Gaussian model. In Fig. S23, we see that at every stage, a Gaussian assumption explains the $4^{th}$ order moments reasonably well, at least up to uncertainty in measurement.

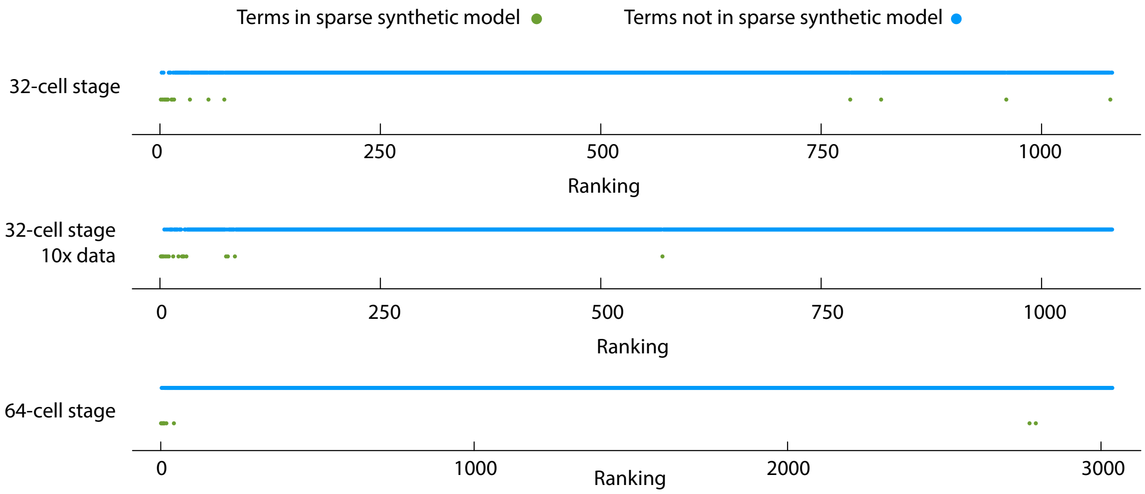


FIG. S21. Inference procedure approximately recovers the true sparse interactions from a synthetic data set. For each cell stage, we sample from the best fit sparse model (see main text Fig. 6), to construct a data set of the same size and quality as the experimental data. We apply the inference procedure to the 32-cell stage synthetic data set and ask where the genuine sparse interaction terms appear in the ranking (top). Further, we can construct a synthetic data set with more data points than the true experimental data to explore how more data affects the robustness of the inference (middle). Terms that appear in the true sparse model are in green, terms that are not in the sparse model are in blue. Out of the 17 true terms in the 32-cell stage synthetic data set, 10 appear in the top 17 inferred terms, and 13 appear in the top 100. For the 64-cell stage (bottom) out of the 12 true terms in the synthetic data set, 8 appear in the top 12 and 10 appear in the top 100.

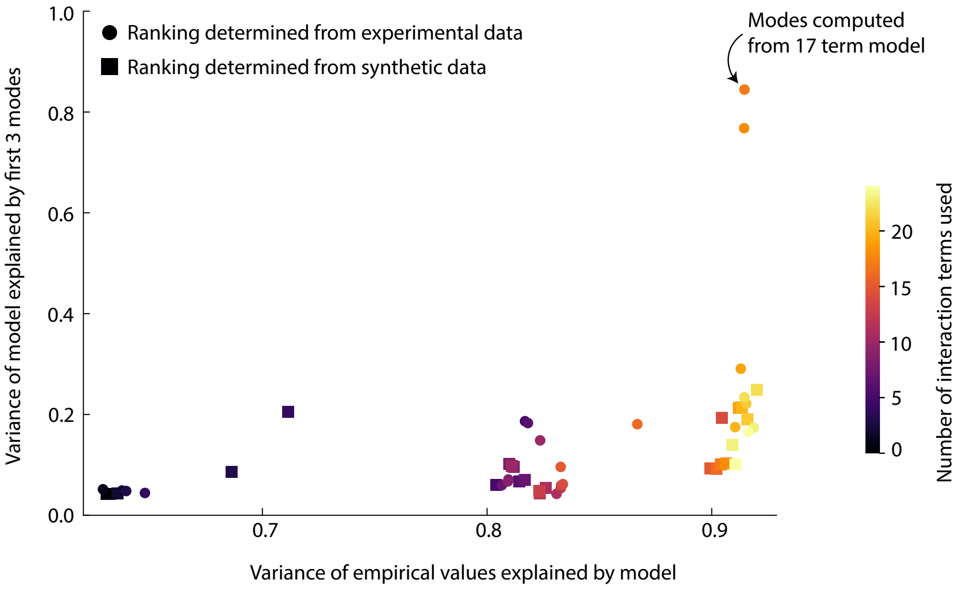


FIG. S22. Inferred modes are robust to the interaction terms in a model. The more interaction terms that are included in a model, the better that model fits the empirical measurements, as measured by the variance explained in covariance values (x-axis). In the 17-term model from which the modes in Fig. 7 were computed, the top 3 modes explain around 80% of the variance in that model. For any other model, we can ask how much variance the 3 modes from the 17-term model explain (y-axis). Despite the fact that vectors in high dimensional spaces are generically orthogonal, for models that fit the empirical data well the modes from the 17-term model explain around 20% of that models variance. Moreover, taking models with interaction terms coming from the synthetic ranking, which do not agree perfectly with the actual ranking (Fig. S21), the modes from the 17-term model still explain around 20% of the model variance. This shows that the inferred modes of the model have a degree of robustness to the exact interaction terms in that model.

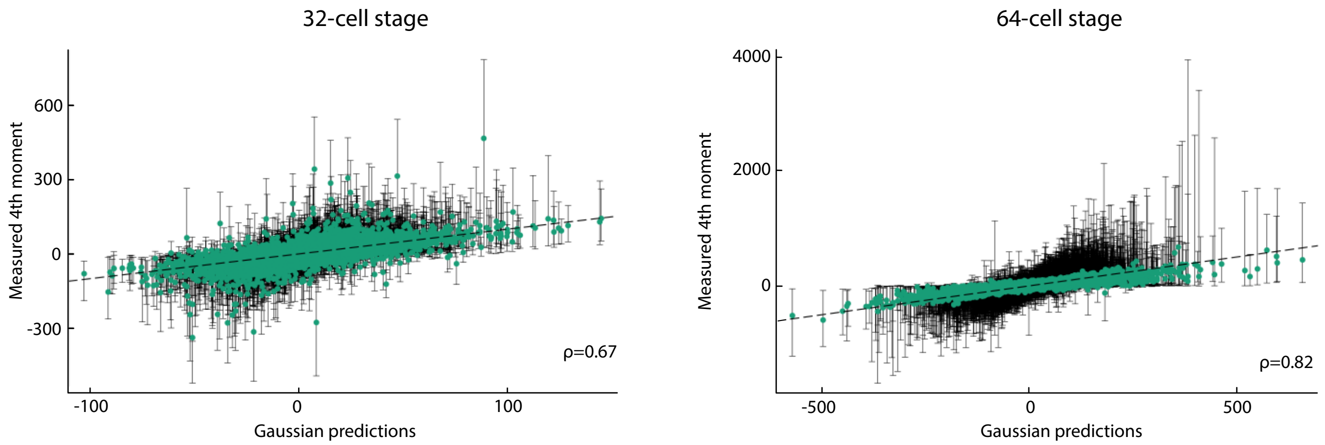


FIG. S23. Comparison of the measured $4^{th}$ order moments compared to the predictions made from $2^{nd}$ order moments assuming a Gaussian distribution. All moments are computed as described in Section. IV. Error bars are from the $25^{th}$ to $75^{th}$ percentile of moment estimates. Also shown is the Pearson correlation coefficient, $\rho$. The agreement, up to error bars, suggests that a Gaussian model is reasonable.

### Appendix A: Maximum entropy Gradients

In order to efficiently fit a maximum entropy model we make use of an analytic gradient during optimization. We outline the calculation of this gradient here. Starting from

$$J_{(pi)(qj)} = \sum_t M^t_{ij} I^t_{pq} + \sum_{k=1}^{n_\Psi} S^k_{ij} \Psi^k_{pq} + \frac{1}{2} \sum_{k'=1}^{n_\Omega} \left( U^{k'}_{ij} \Omega^{k'}_{pq} + U^{k'}_{ji} \Omega^{k'}_{qp} \right), \tag{S1}$$

We can compute $J^{-1}$ using

$$\frac{\partial (J^{-1})_{ab}}{\partial \alpha} = -\sum_{c,d} (J^{-1})_{ac} \frac{\partial J_{cd}}{\partial \alpha} (J^{-1})_{db}, \tag{S2}$$

we find that

$$\begin{aligned}
\frac{\partial}{\partial M^t_{ij}} (J^{-1})_{ab} &= -\sum_{p \in t} (J^{-1})_{a(pi)} (J^{-1})_{(pj)b} \\
\frac{\partial}{\partial S^k_{ij}} (J^{-1})_{ab} &= -\sum_{p,s} (J^{-1})_{a(pi)} \Psi^k_{ps} (J^{-1})_{(sj)b}, \\
\frac{\partial}{\partial U^k_{ij}} (J^{-1})_{ab} &= -\frac{1}{2} \sum_{p,s} \left[ (J^{-1})_{a(pi)} \Omega^k_{ps} (J^{-1})_{(sj)b} + (J^{-1})_{a(pj)} \Omega^k_{sp} (J^{-1})_{(si)b} \right],
\end{aligned}$$

where we have not yet assumed $M$ or $S$ to be symmetric. In total, we can calculate the gradient as, e.g.,

$$\frac{\partial \tilde{L}}{\partial M^t_{ij}} = \sum_{a,b} \frac{\partial (J^{-1})_{ab}}{\partial M^t_{ij}} \frac{\partial \tilde{L}}{\partial (J^{-1})_{ab}}. \tag{S3}$$

Once we have found the matrix $Y_{ab} = \partial \tilde{L} / \partial (J^{-1})_{ab}$, we can construct the gradient as

$$\frac{\partial \tilde{L}}{\partial M_{ij}^t} = -\sum_{p\in t}\sum_{a,b}(J^{-1})_{(pi)a}Y_{ab}(J^{-1})_{b(pj)} \tag{S4}$$
$$= -\sum_{p\in t}(J^{-1}YJ^{-1})_{(pi)(pj)}$$
$$\frac{\partial \tilde{L}}{\partial S_{ij}^k} = -\sum_{p,s}(J^{-1}YJ^{-1})_{(pi)(sj)}\Psi_{ps}^k$$
$$\frac{\partial \tilde{L}}{\partial U_{ij}^k} = -\frac{1}{2}\sum_{p,s}(J^{-1}YJ^{-1})_{(pi)(sj)}\Omega_{ps}^k + (J^{-1}YJ^{-1})_{(pj)(si)}\Omega_{sp}^k.$$

For notational convenience, let $X = J^{-1}$, so that

$$\tilde{L} = \tilde{L}(X) = \sum_{i,j,t}\left(C_{ij}^t - \frac{1}{n_t}\sum_{r\in t}X_{(ri)(rj)}\right)^2 \tag{S5}$$
$$+ \sum_{i,j,t\le t'}\left(\Gamma_{ij}^{tt'} - \frac{1}{n_t n_{t'}}\sum_{r\in t,q\in t'}X_{(ri)(qj)}\right)^2,$$
$$= \sum_{i,j,t}\omega_{ij}^t\omega_{ij}^t + \sum_{i,j,t\le t'}\mu_{ij}^{tt'}\mu_{ij}^{tt'}$$

Then,

$$\frac{\partial \omega_{ij}^t}{\partial X_{(pk)(ql)}} = -\frac{1}{n_t}\delta_{ki}\delta_{lj}\delta_{pq}1_{q\in t}, \tag{S6}$$
$$\frac{\partial \mu_{ij}^{tt'}}{\partial X_{(pk)(ql)}} = -\frac{1}{n_t n_{t'}}\delta_{ki}\delta_{lj}1_{p\in t}1_{q\in t'},$$

so that

$$Y_{(pk)(ql)} = \frac{\partial \tilde{L}}{\partial X_{(pk)(ql)}} = -\frac{2}{n_{t_p}}\delta_{pq}\omega_{kl}^{t_p} - \frac{2}{n_{t_p}n_{t_q}}\mu_{kl}^{t_p t_q}1_{t_p\le t_q}, \tag{S7}$$

where $t_p$, $t_q$ are the cell types of indices $p$ and $q$ respectively.

---

[14] Y. Roudi, E. Aurell, and J. Hertz. Statistical physics of pairwise probability models. *Front. Comput. Neuro.*, 3, 2009.

[15] R. Satija, J. A. Farrell, D. Gennert, A. F. Schier, and A. Regev. Spatial reconstruction of single-cell gene expression data. *Nat. Biotech.*, 33(5):495, 2015.

[16] M. Schulze. A new monotonic, clone-independent, reversal symmetric, and condorcet-consistent single-winner election method. *Social Choice and Welfare*, 36(2):267303, 2011.

[17] Y. Senbabaoğlu, G. Michailidis, and J. Z. Li. Critical limitations of consensus clustering in class discovery. *Sci. Reps*, 4(1):6207, 2014.

[18] H. L. Sladitschek, U.-M. Fiuza, D. Pavlinic, V. Benes, L. Hufnagel, and P. A. Neveu. Morphoseq: Full single-cell transcriptome dynamics up to gastrulation in a chordate. *Cell*, 181(4):922.e21, 2020.

[19] G. Stepaniants, A. D. Hastewell, D. J. Skinner, J. F. Totz, and J. Dunkel. Discovering dynamics and parameters of nonlinear oscillatory and chaotic systems from partial observations, 2023.

[20] V. A. Traag, L. Waltman, and N. J. van Eck. From louvain to leiden: guaranteeing well-connected communities. *Sci. Rep.*, 9(1):5233, 2019.

[21] F. A. Wolf, P. Angerer, and F. J. Theis. Scanpy: large-scale single-cell gene expression data analysis. *Genome Biol.*, 19(1):15, 2018.
